# Evolutionary dynamics of bacteria in the gut microbiome within and across hosts

**DOI:** 10.1101/210955

**Authors:** Nandita R. Garud, Benjamin H. Good, Oskar Hallatschek, Katherine S. Pollard

**Affiliations:** Gladstone Institutes, San Francisco, California, United States of America; Department of Physics, University of California, Berkeley, California, United States of America; Department of Bioengineering, University of California, Berkeley, California, United States of America; Department of Integrative Biology, University of California, Berkeley, California, United States of America; Kavli Institute for Theoretical Physics, University of California, Santa Barbara, California, United States of America; Department of Epidemiology & Biostatistics, Institute for Human Genetics, Quantitative Biology Institute, and Institute for Computational Health Sciences, University of California, San Francisco, California, United States of America; Chan-Zuckerberg Biohub, San Francisco, California, United States of America

**Author notes:** These authors contributed equally and are ordered alphabetically (Nandita R. Garud, Benjamin H. Good). (B.H.G.), (N.R.G.).

## Abstract

Gut microbiota are shaped by a combination of ecological and evolutionary forces. While the ecological dynamics have been extensively studied, much less is known about how species of gut bacteria evolve over time. Here we introduce a model-based framework for quantifying evolutionary dynamics within and across hosts using a panel of metagenomic samples. We use this approach to study evolution in ∼30 prevalent species in the human gut. Although the patterns of between-host diversity are consistent with quasi-sexual evolution and purifying selection on long timescales, we identify new genealogical signatures that challenge standard population genetic models of these processes. Within hosts, we find that genetic differences that accumulate over ∼6 month timescales are only rarely attributable to replacement by distantly related strains. Instead, the resident strains more commonly acquire a smaller number of putative evolutionary changes, in which nucleotide variants or gene gains or losses rapidly sweep to high frequency. By comparing these mutations with the typical between-host differences, we find evidence that some sweeps are seeded by recombination, in addition to new mutations. However, comparisons of adult twins suggest that replacement eventually overwhelms evolution over multi-decade timescales, hinting at fundamental limits to the extent of local adaptation. Together, our results suggest that gut bacteria can evolve on human-relevant timescales, and they highlight the connections between these short-term evolutionary dynamics and longer-term evolution across hosts.

## Introduction

The gut microbiome is a complex ecosystem comprised of a diverse array of microbial organisms. The abundances of different species and strains can vary dramatically based on diet [1], host-species [2], and the identities of other co-colonizing taxa [3]. These rapid shifts in community composition suggest that individual gut microbes may be adapted to specific environmental conditions, with strong selection pressures between competing species or strains. Yet while these ecological responses have been extensively studied, much less is known about the evolutionary forces that operate within populations of gut bacteria, both within individual hosts, and across the larger host-associated population. This makes it difficult to predict how rapidly strains of gut microbes will evolve new ecological preferences when faced with environmental challenges, like drugs or diet, and how the genetic composition of the community will change as a result.

The answers to these questions depend on two different types of information. At a mechanistic level, one must understand the functional traits that are under selection in the gut, and how they may be modified genetically. Recent work has started to address this question, leveraging techniques from comparative genomics [4–6], evolution in model organisms [7–9], and high-throughput genetic screens [10, 11]. Yet in addition to the targets of selection, evolution also depends on population genetic processes that describe how mutations spread through a population of gut bacteria, both within individual hosts, and across the larger population. These dynamical processes can strongly influence which mutations are likely fix within a population, and the levels of genetic diversity that such populations can maintain. Understanding these processes is the goal of our present work.

Previous studies of pathogens [12], laboratory evolution experiments [13], and some environmental communities [14–17] have shown that microbial evolutionary dynamics are often dominated by rapid adaptation, with new variants accumulating within months or years [7, 14, 18–25]. However, it is not clear how this existing picture of microbial evolution extends to a more complex and established ecosystem like the healthy gut microbiome. On the one hand, hominid gut bacteria have had many generations to adapt to their host environment [26], and may not be subject to the continually changing immune pressures faced by many pathogens. The large number of potential competitors in the gut ecosystem may also provide fewer opportunities for a strain to adapt to new conditions before an existing strain expands to fill the niche [27, 28] or a new strain invades from outside the host. On the other hand, it is also possible that small-scale environmental fluctuations, either driven directly by the host or through interactions with other resident strains, might increase the opportunities for local adaptation [29]. If immigration is restricted, the large census population size of gut bacteria could allow residents to produce and fix adaptive variants rapidly before a new strain is able to invade. In this case, one could observe rapid adaptation on short timescales, which is eventually arrested on longer timescales as strains are exposed to the full range of host environments. Additional opportunities for adaptation can occur if the range of host environments also shifts over time (e.g., due to urbanization, antibiotic usage, etc.). Determining which of these scenarios apply to gut communities is critical for efforts to study and manipulate the microbiome.

While traditional amplicon sequencing provides limited resolution to detect within-species evolution [30], whole-genome shotgun metagenomic sequencing is starting to provide the raw polymorphism data necessary to address these questions [31]. In particular, several reference-based approaches have been developed to detect genetic variants within individual species in larger metagenomic samples [31–36]. While these approaches enable strain-level comparisons between samples, they have also documented substantial within-species variation in individual metagenomes [31, 35, 37]. This makes it difficult to assign an evolutionary interpretation to the genetic differences between samples, since they arise from unobserved mixtures of different bacterial lineages.

Several approaches have been developed to further resolve these mixed populations into individual haplotypes or “strains”. These range from simple consensus approximations [35,37,38], to sophisticated clustering algorithms [39,40] and the incorporation of physical linkage information [41]. However, while these methods are useful for tracking well-defined strains across samples, it is not known how their assumptions and failure modes might bias inferences of evolutionary dynamics, particularly among closely related strains. As a result, the evolutionary processes that operate within species of gut bacteria remain poorly characterized.

In this study, we take a different approach to the strain detection problem that is specifically designed for inferring evolutionary dynamics in a large panel of metagenomes. Building on earlier work by [4, 35], we show that many prevalent species have a subset of hosts for which a portion of the dominant lineage is much easier to identify. By focusing only on this subset of “quasi-phaseable” samples, we develop methods for resolving small differences between the dominant lineages with a high degree of confidence.

We use this approach to analyze a large panel of publicly available human stool samples [42–46], which allows us to quantify evolutionary dynamics within and across hosts in ∼30 prevalent bacterial species. We find that the long-term evolutionary dynamics across hosts are broadly consistent with models of quasi-sexual evolution and purifying selection, with relatively weak geographic structure in many prevalent species. However, our quantitative approach also reveals interesting departures from standard population genetic models of these processes, which suggests that new models are required to fully understand the evolutionary dynamics that take place across the larger population.

We also use our approach to detect examples of within-host adaptation, in which nucleotide variants or gene gains or losses rapidly sweep to high frequency within ∼6 month intervals. We find within-host sweeps may be seeded by recombination, in addition to *de novo* mutations, as might be expected for complex ecosystems with large census population sizes and frequent horizontal exchange. However, by analyzing differences between adult twins, we find that short-term evolution can eventually be overwhelmed by the invasion of distantly related strains on multi-decade timescales. This suggests that resident strains are rarely able to become so well-adapted to a particular host that they prevent future replacements. Together, these results show that the gut microbiome is a promising system for studying the dynamics of microbial evolution in a complex community setting. The framework we introduce may also be useful for characterizing evolution of microbial communities in other environments.

## Methods

### Resolving within-host lineage structure in a panel of metagenomic samples

To investigate evolutionary dynamics within species in the gut microbiome, we analyzed shotgun metagenomic data from a panel of stool samples from 693 healthy individuals sequenced in previous work (Table S1). This panel includes 250 North American subjects sequenced by the Human Microbiome Project [42, 44], a subset of which were sampled at 2 or 3 timepoints roughly 6-12 months apart. To probe within-host dynamics on longer timescales, we also included data from a cohort of 125 pairs of adult twins from the TwinsUK registry [45], and 4 pairs of younger twins from Ref. [46]. Previous work has demonstrated that twins are often colonized by the nearly identical strains in childhood [46], so that differences between twins may serve as a proxy for the temporal changes that accumulate over the decades since childhood. Finally, to further control for geographic structure, we also included samples from 185 Chinese subjects sequenced at a single timepoint [43].

We used a standard reference-based approach to measure single nucleotide variant (SNV) frequencies and gene copy number across a panel of prevalent species for each metagenomic sample (see SI Section S1 Text for details on the bioinformatic pipeline, including mapping parameters and other filters). Descriptive summaries of this genetic variation have been reported elsewhere [31, 33–35, 37, 44]. Here, we revisit these patterns to investigate how they emerge from the lineage structure set by the host colonization process. Using these results, we then show how certain aspects of this lineage structure can be inferred from the statistics of within-host polymorphism, which enable measurements of evolutionary dynamics across samples.

As an illustrative example, we first focus on the patterns of polymorphism in *Bacteroides vulgatus*, which is among the most abundant and prevalent species in the human gut. These properties ensure that the *B. vulgatus* genome has high-coverage in many samples, which enables more precise estimates of the allele frequencies in each sample (Fig. 1A-D). The overall levels of within-host diversity for this species are summarized in Fig. 1E, based on the fraction of synonymous sites in core genes with intermediate allele frequencies (white region in Figs. 1A-D). This measure of within-host genetic variation varies widely across the samples: some metagenomes have only a few variants along the *B. vulgatus* genome, while others have mutations at more than 1% of all synonymous sites (comparable to the differences between samples, Fig. S2).

**Fig 1.**
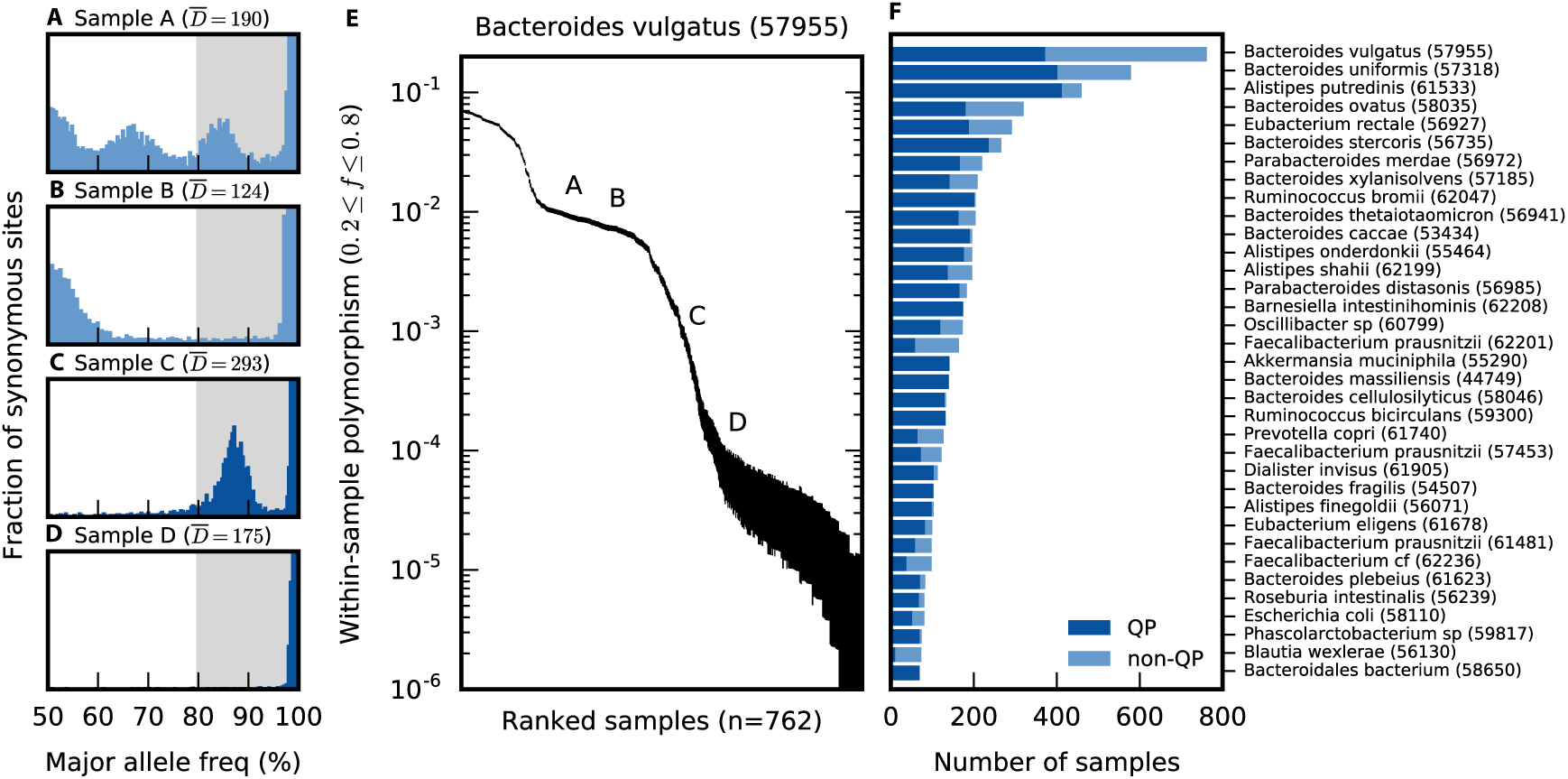
Genetic diversity within hosts. *Bacteroides vulgatus* is shown as an example in panels a-e. (a-d) The distribution of major allele frequencies at synonymous sites in the core genome for four different samples, with the median read depth 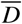 listed above each panel. To emphasize the distributional patterns, the vertical axis is scaled by an arbitrary normalization constant in each panel, and it is truncated for visibility. The white region denotes the intermediate frequency range used for the polymorphism calculations below. (e) The average fraction of synonymous sites in the core genome with major allele frequencies ≤80% (white region in a-d), for all samples with 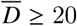 Vertical lines denote 95% posterior confidence intervals based on the observed number of counts (Text S2 Text). For comparison, the corresponding values for the samples in panels (a-d) are indicated by the numbers (1-4). (f) The distribution of quasi-phaseable (QP) samples among the 35 most prevalent species, arranged by descending prevalence; the distribution across hosts is shown in Fig. S4. For comparison, panels (c) and (d) are classified as quasi-phaseable, while panels (a) and (b) are not.

We first asked whether these patterns are consistent with a model in which each host is colonized by a single *B. vulgatus* clone, so that the intermediate frequency variants represent mutations that have arisen since colonization. Assuming this model and using conservatively high estimates for per site mutation rates (µ ∼10^-9^ [47]), generation times (∼10 per day [48]), and time since colonization (< 100 years), we estimate a neutral polymorphism rate < 10^-3^ at each synonymous site (Text S2.1), which is at odds with the higher levels of diversity in many samples (Fig. 1E). Instead, we conclude that the samples with higher synonymous diversity must have been colonized by multiple divergent bacterial lineages that accumulated mutations for many generations before coming together in the same gut community.

As a plausible alternative, we next asked whether the data are consistent with a large number of colonizing lineages (*n*_*c*_ ≫ 1) drawn at random from the broader population. However, this process is expected to produce fairly consistent polymorphism rates and allele frequency distributions in different samples, which is at odds with the variability we observe even among the high-diversity samples (e.g., Figs. 1A,B). Instead, we hypothesize that many of the high-diversity hosts have been colonized by just a few diverged lineages [i.e., (*n*_*c*_ -1) ∼ 𝒪 (1)]. Consistent with this hypothesis, the distribution of allele frequencies in each host is often strongly peaked around a few characteristic frequencies (Fig. 1A-D), suggesting a mixture of several distinct lineages. Similar findings have recently been reported in a number of other host-associated microbes, including several species of gut bacteria [4, 35, 49, 50]. Figures 1A-C show that hosts can vary both in the apparent number of colonizing lineages, and the frequencies at which they are mixed together. As a result, we cannot exclude the possibility that even the low diversity samples (e.g. Fig. 1D) are colonized by multiple lineages that happen to fall below the detection threshold set by the depth of sequencing. We will refer to samples with a small number of diverged lineages as an *oligo-colonization* model, in order to contrast with the single-colonization (*n*_*c*_ = 1) and multiple-colonization (*n*_*c*_ ≫ 1) alternatives above.

#### Quasi-phaseable (QP) samples

Compared to the single- and multiple-colonization models, the oligo-colonization model makes it more difficult to identify evolutionary changes between lineages. In this scenario, individual hosts are not clonal, but the within-host allele frequencies derive from idiosyncratic colonization processes, rather than a large random sample from the population. To disentangle genetic changes between lineages from these host-specific factors, we must estimate phased haplotypes (or “strains”) from the distribution of allele frequencies within individual hosts. This is a complicated inverse problem, and we will not attempt to solve the general case here. Instead, we adopt an approach similar to Ref. [35] and others, and leverage the fact that the lineage structure in *certain* hosts is sufficiently simple that we can assign alleles to the dominant lineage with a high degree of confidence.

Our approach is based on the simple observation that two high-frequency variants must co-occur in an appreciable fraction of cells (Text S3.1). This “pigeonhole principle” suggests that we can estimate the genotype of one of the lineages in a mixed sample by taking the major alleles present above some threshold frequency *f* ^*^≫ 50%, and treating the remaining sites as missing data. Although the potential errors increase with the length of the inferred haplotype, we will not actually require genome-length haplotypes for our analysis here. Instead, we leverage the fact that significant evolutionary information is already encoded in the marginal distributions of one- and two-site haplotypes, so that these “quasi-phased” lineages will be sufficient for our present purposes.

The major challenge with this approach is that we do not observe the true allele frequency directly, but must instead estimate it from a noisy sample of sequencing reads. This can lead to phasing errors when the true major allele is sampled at low frequency by chance and is assigned to the opposite lineage (Fig. S1). The probability of such “polarization” errors can vary dramatically depending on the sequencing coverage and the true frequency of the major allele (Text S3.2). Previous approaches based on consensus alleles [35, 37] can therefore induce an unknown number of errors that make it difficult to confidently detect a small number of evolutionary changes between samples.

In Text S3.3, we show that by explicitly modeling the sampling error process, the expected probability of a polarization error in our cohort can be bounded to be sufficiently low if we take *f* ^*^ = 80%, and if we restrict our attention to samples with sufficiently high coverage and sufficiently low rates of intermediate-frequency polymorphism. We will refer to these as *quasi-phaseable* (QP) samples. In the *B. vulgatus* example above, Figs. 1C,D are classified as quasi-phaseable, while Figs. 1A,B are not. Note that quasi-phaseability is separately defined for each species in a metagenomic sample, rather than for the sample as a whole. For simplicity, we will still refer to these species-sample combinations as QP samples, with the implicit understanding that they refer to a particular focal species.

In Fig. 1F, we plot the distribution of QP samples across the most prevalent gut bacterial species in our panel. The fraction of QP samples varies between species, ranging from ∼50% in the case of *P. copri* to nearly 100% for *B. fragilis* [4], and it accounts for much of the variation in the average polymorphism rate (Fig. S3). Most individuals carry a mixture of QP and non-QP species (Fig. S4), which suggests that quasi-phaseability arises independently for each species in a sample, rather than for the sample as a whole. Thus, although many species-sample combinations are non-QP, in a cohort of a few hundred samples it is not uncommon to find ≥ 50 QP samples in many prevalent species, which yields ∼ 3000 quasi-phased haplotypes when aggregated across the different species. Consistent with previous studies of the stability of personal microbiomes [31, 35, 51], a majority of the longitudinally sampled species maintain their QP classification at both timepoints, though this pattern is not universal (Fig. S5). We will revisit the peculiar properties of this within-host lineage distribution in the Discussion. For the remainder of the analysis, we will take the distribution in Fig. 1F as given and focus on leveraging the QP samples to quantify the evolutionary changes that accumulate between lineages in different samples.

We investigate two types of evolutionary changes between lineages in different QP samples. The first class consists of single nucleotide differences, which are defined as SNVs that segregate at frequencies ≤ 1*-f* ^*^in one sample and *≥f* ^*^ in another, with *f* ^*^*≈*80% as above (Fig. S1). These thresholds are chosen to ensure low genome-wide false positive rate given the typical coverage and allele frequency distributions among the QP samples in our panel (Text S3.4). The second class consists of differences in gene presence or absence, in which the relative copy number of a gene, *c*, is below the threshold of detection (*c* < 0.05) in one sample, and is consistent with a single-copy gene (0.6 < *c* < 1.2, see Fig. S6) in the other sample. These thresholds are chosen to ensure a low genome-wide false positive rate across the QP samples, given the typical variation in sequencing coverage along the genome (Text S3.5), and to minimize mapping artifacts (Text S1.3).

Note that these SNV and gene changes represent only a subset of the potential differences between lineages, since they neglect other evolutionary changes (e.g., indels, genome rearrangements, or changes in high copy number genes) that are more difficult to quantify in a metagenomic sample, as well as more subtle changes in allele frequency and gene copy number that do not reach our stringent detection thresholds. We will revisit these and other limitations in more detail in the Discussion.

## Results

### Long-term evolution across hosts

By focusing on the QP samples for each species, we can measure genetic differences between lineages in different hosts, as well as within hosts over short time periods. Descriptive summaries of this variation have been reported elsewhere [31, 33–35, 37, 44]. Here, we aim to leverage these patterns (and the increased resolution of the QP samples) to quantify the evolutionary dynamics that operate within species of gut bacteria, both within and across hosts.

To interpret within-host changes in an evolutionary context, it will be useful to first understand the structure of genetic variation between lineages in different hosts. This variation reflects the long-term population genetic forces that operate within each species, presumably integrating over many rounds of colonization, growth, and dispersal. To investigate these forces, we first analyzed the average nucleotide divergence between strains of a given species in different pairs of QP hosts (Fig. 2A). In the case of twins, we included only a single host from each pair, to better approximate a random sample from the population.

**Fig 2.**
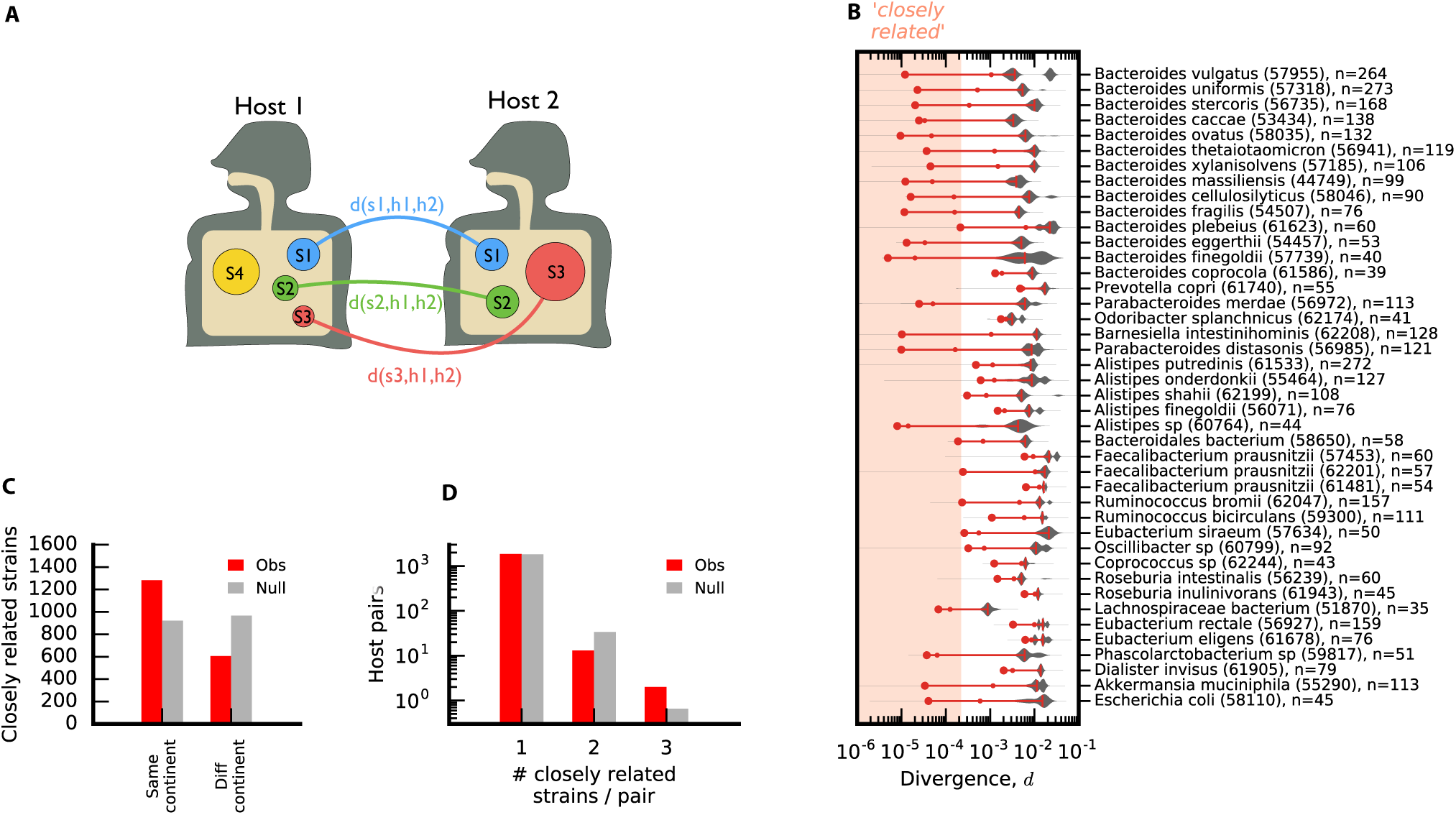
Between-host divergence across prevalent species of gut bacteria. (a) Schematic illustration. For a given pair of hosts (h1, h2), core-genome nucleotide divergence (*d*) is computed for each species (s1, s2, etc.) that is quasi-phaseable in both hosts. (b) Distribution of *d* across all pairs of unrelated hosts for a panel of prevalent species. Species are sorted according to their phylogenetic distances [33], with the number of QP hosts indicated in parentheses; species were only included if they had at least 33 QP hosts (> 500 QP pairs). Symbols denote the median (dash), 1-percentile (small circle), and 0.1-percentile (large circle) of each distribution, and are connected by a red line for visualization; for distributions with < 10^3^ data points, the 0.1-percentile is estimated by the second-lowest value. The shaded region denotes our ad-hoc definition of “closely related” divergence, *d* ≤2 × 10^-4^. (c) The distribution of the number of species with closely related strains in distinct hosts present in the same or different continents. The null distribution is obtained by randomly permuting hosts within each species. Though the observed values are significantly different than the null (*P* < 10^-4^), the large contribution from different continents shows that closely related strains are not solely a product of geographic separation. (d) The distribution of the number of species with closely related strains for each pair of hosts. The null distribution is obtained by randomly permuting hosts independently within each species (*n* = 10^3^ permutations, *P* ≈ 0.9). This shows that there is no tendency for the same pairs of hosts to have more closely related strains than expected under the null distribution above.

Figure 2B shows the distribution of pairwise divergence, averaged across the core genome, for ∼ 40 of the most prevalent bacterial species in our cohort. In a panmictic, neutrally evolving population, we would expect these distances to be clustered around their average value, *d* ≈ 2µ*T*_*c*_, where *T*_*c*_ is the coalescent timescale for the across-host population [52]. By contrast, Fig. 2 shows striking differences in the degree of relatedness for strains in different hosts. Even at this coarse, core-genome-wide level, the genetic distances vary over several orders of magnitude.

Some species show multiple peaks of divergence for high values of *d*, consistent with the presence of subspecies [36], ecotypes [53, 54], or other strong forms of population structure. These coarse groupings have been observed previously and are not our primary focus here. Rather, we seek to understand the population genetic forces that operate at finer levels of taxonomic resolution.

From this perspective, the more surprising parts of Fig. 2 are the thousands of pairs of lineages with extremely low between-host divergence (e.g. *d*; S 0.01%), more than an order of magnitude below the median values in most species. Similar observations have recently been reported by Ref. [35], and are often interpreted as *strain sharing* across hosts. However, the evolutionary interpretation of these closely strains remains unclear.

#### Closely related strains reflect population genetic processes, rather than cryptic host relatedness

The simplest explanation for a long tail of closely related strains is cryptic relatedness [55], arising from a breakdown of random sampling. For microbes, this can occur when two cells are sampled from the same clonal expansion, e.g., when strains are transferred between mothers and infants [33, 56], between cohabitating individuals [46], or within a hospital outbreak [57]. While these transmission events have been observed in other studies, they are unlikely to account for the patterns here. All of the lineages in Fig. 2 are sampled from individuals in different households, and more than a third of the closely related pairs derive from individuals on different continents (Fig. 2B).

Of course, there could still be some other geographic variable, beyond household or continent of origin, that could explain an elevated probability of transmission between two individuals. Fortunately, our metagenomic approach allows us to rule out these additional sources of cryptic host relatedness by leveraging multiple species comparisons for the same pair of hosts. If there was a hidden geographic variable, then we would expect that individuals with closely related strains in one species would be much more likely to share closely related strains in other species as well. However, we observe only a small fraction of hosts that share multiple closely related strains (Fig. 2C), consistent with a null model in which these strains are randomly and independently distributed across hosts. This suggests that host-wide sampling biases are not the primary driver of the closely related strains in Fig. 2. Though the rates of nucleotide divergence are low, the vast majority of these strains are still genetically distinguishable from each other. The number of SNV differences typically exceeds our estimated false positive rate (Fig. S7A, Text S3.4), and these single nucleotide variants are typically accompanied by ≳ 10 differences in gene content between the two strains (Fig. S7B). Furthermore, closely related strains frequently differed in their collections of private marker SNVs (Fig. S8), which are often used to track strain transmission events [33, 46]. Together, these lines of evidence suggest that closely related strains are often genetically distinct, and do not arise from a simple clonal expansion. Instead, the data suggest that there are additional population genetic timescales beyond *T*_*c*_ that are relevant for microbial evolution.

This hypothesis is bolstered by the large number of species, particularly in the *Bacteroides* genus, with anomalously low divergence rates between some pairs of hosts. However, we note that this pattern is not universal: some genera, like *Alistipes* or *Eubacterium*, show more uniform rates of divergence between hosts. Apart from these phylogenetic correlations, we cannot yet explain why some species have low divergence host pairs and others do not. Natural candidates like sample size, abundance, vertical transmissibility [33], or sporulation score [58] struggle to explain the differences between *Bacteroides* and *Alistipes*.

#### Closely related strains have distinct signatures of natural selection

We next examined how natural selection influences the genetic diversity observed between hosts. Previous work has suggested that genetic diversity in many species of gut bacteria is strongly constrained by purifying selection, which purges deleterious mutations that accumulate between hosts [31]. However, the temporal dynamics of this process remain poorly understood. We do not know whether purifying selection acts quickly enough to prevent deleterious mutations from spreading to other hosts, or if deleterious mutations typically spread across multiple hosts before they are purged. In addition, it is plausible that the dominant mode of natural selection could be different for the closely related strains above (e.g. if they reflect recent ecological diversification [15]).

To address these questions, we analyzed the relative contribution of synonymous and nonsynonymous mutations that comprise the overall divergence rates in Fig. 2A. We focused on the ratio between the per-site divergence at nonsynonymous sites (*d*_*N*_) and the corresponding value at synonymous sites (*d*_*S*_). Under the assumption that synonymous mutations are effectively neutral, the ratio *d*_*N*_/*d*_*S*_ measures the average action of natural selection on mutations at nonsynonymous sites.

In Fig. 3, we plot these *d*_*N*_/*d*_*S*_ estimates across every pair of QP hosts for each of the prevalent species in Fig. 2A. The values of *d*_*N*_/*d*_*S*_ are plotted as a function of *d*_*S*_, which serves as a proxy for the average divergence time across the genome. We observe a consistent negative relationship between these two quantities across the prevalent species in Fig. 2.

**Fig 3.**
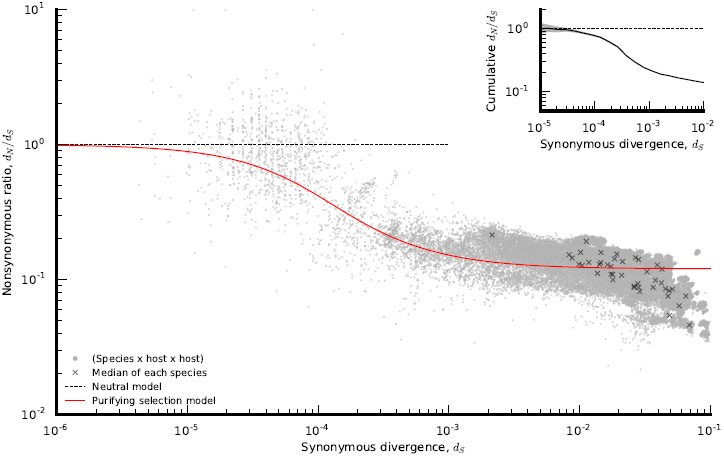
Signatures of selective constraint within species as a function of core-genome divergence. Ratio of divergence at nondegenerate nonsynonymous sites (*d*_*N*_) and fourfold degenerate synonymous sites (*d*_*S*_) as a function of *d*_*S*_ (SI Section S4 Text) for all species x host1 x host2 combinations in Fig. 2 (grey circles). Crosses (**x**) denote species-wide estimates obtained from the ratio of the median *d*_*N*_ and *d*_*S*_ within each species. The red line denotes the theoretical prediction from the purifying selection null model in SI Section S4 Text. (inset) Ratio between the cumulative private *d*_*N*_ and *d*_*S*_ values for all QP host pairs with core-genome-wide synonymous divergence less than *d*_*S*_. The narrow shaded region denotes 95% confidence intervals estimated by Poisson resampling (Text S4 Text), which shows that *d*_*N*_/*d*_*S*_ ≳ 1 even for low *d*_*S*_.

For large divergence times (*d*_*S*_ ∼1%), we observe only a small fraction of nonsynonymous mutations (*d*_*N*_/*d*_*S*_ ∼0.1), indicating widespread purifying selection on amino-acid replacements [31]. Yet among more closely related strains, we observe a much higher fraction of nonsynonymous changes, with *d*_*N*_/*d*_*S*_ approaching unity when *d*_*S*_ ∼0.01%. (We observe a similar trend if we restrict our attention to singleton SNVs, Fig. S9). Moreover, this negative relationship between *d*_*N*_/*d*_*S*_ and *d*_*S*_ is much more pronounced than the between-species variation in the typical values of *d*_*N*_/*d*_*S*_ (black crosses in Fig. 3). While between-species variation may be driven by mutational biases, the strong within-species signal indicates that there are consistent differences in the action of natural selection as a function of time.

In principle, the *d*_*N*_/*d*_*S*_ increases in the recent past could be driven by interesting biological processes, like enhanced adaptation or ecological diversification on short timescales, or a recent global shift in selection pressures caused by host-specific factors (e.g., the introduction of agriculture). However, the data in Fig. 3 appear to be well explained by an even simpler null model of purifying selection, where deleterious mutations are purged over a timescale inversely proportional to their cost (SI Section S4 Text). This dynamical model can explain the varying signatures of natural selection without requiring that the selective pressures themselves vary over time. We find reasonable quantitative agreement for a simple distribution of fitness effects, in which 10% of nonsynonymous sites are neutral, and the remaining 90% have fitness costs on the order of *s*/µ ∼10^5^. Though the true model is likely more complicated, we argue that this simple null model should be excluded before more elaborate explanations are considered.

For example, unambiguous proof of recent adaptation could be observed if *d*_*N*_/*d*_*S*_ consistently exceeded one among the most closely related strains, since this can only occur by chance under purifying selection. While a few of the individual comparisons in Fig. 3A have *d*_*N*_/*d*_*S*_ > 1, the cumulative version in Fig. 3B shows that *d*_*N*_/*d*_*S*_ does not significantly exceed one, even for the lowest values of *d*_*S*_. This suggests that, if positive selection is present, it is not sufficiently widespread to overpower the signal of purifying selection in these global *d*_*N*_/*d*_*S*_ measurements. However, there is also substantial variation around the average trend in Fig. 3, which could hide important biological variation among species (or among different genomic regions in the same species). Resolving the signatures of natural selection at these finer scales remains an important avenue for future work.

#### Quasi-sexual evolution on intermediate timescales

In principle, the large range of genome-wide divergence in Figs. 2 and 3 could arise in a model with strong population structure, in which all but the most closely related strains are genetically isolated from each other [59]. Such isolation can be driven by geography as well as ecological diversification [15]. Here, we leverage our quasi-phasing approach to show that genetic isolation cannot account for the patterns in Figs. 2 and 3. Instead, we find that the core genomes of many prevalent gut bacterial species evolve in a quasi-sexual manner, with frequent genetic exchange among individual strains.

Recombination alters the genealogical relationships between strains in different portions of the genome [52]. We therefore sought evidence for recombination by searching for inconsistencies between the genealogies encoded in individual SNVs, and those encoded in the genome-wide divergences in Fig. 2. To minimize the inherent uncertainties involved in phylogenetic inference, we developed a new method for quantifying the phylogenetic inconsistency of a given SNV directly from the pairwise divergence distribution in Fig. 2B (see Fig. S10, Text S5.1). This method also provides an estimate of the maximum age of each SNV, assuming purely clonal evolution. By combining these estimates, we can quantify the extent of phylogenetic inconsistency of SNVs in each species as a function of time (Fig. 4A).

**Fig 4.**
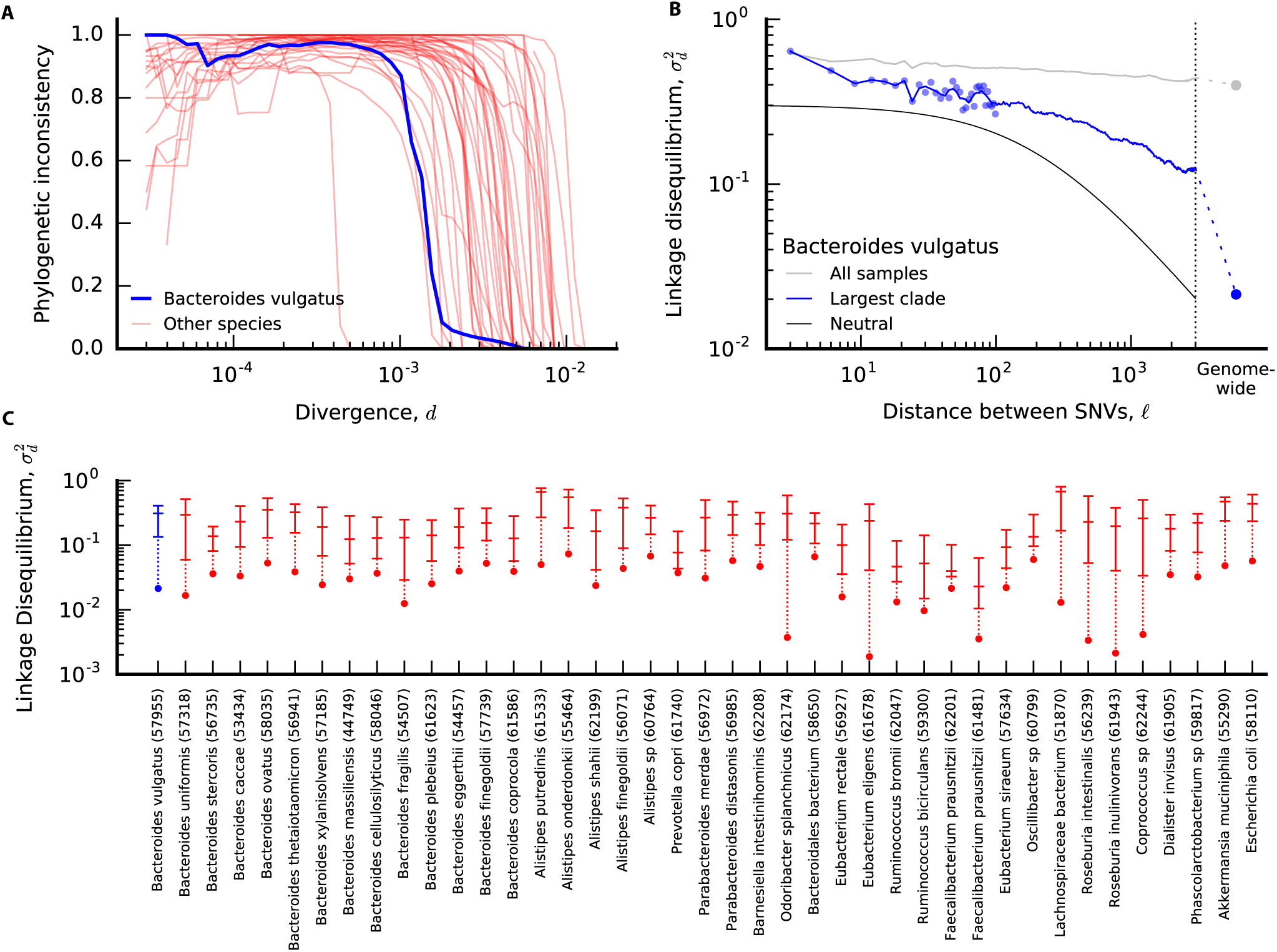
Recombination between strains across hosts. (a) Fraction of phylogenetically inconsistent SNVs as a function of core-genome-wide divergence *d* for each of the species in Fig. ≥ 2 (see Text S5.1). Phylogenetic inconsistency is assessed for the subset of fourfold degenerate synonymous sites in the core genome with 2 minor alleles. *Bacteroides vulgatus* is highlighted in blue, while the remaining species are shown in light red. (b) Linkage disequilibrium 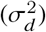 as a function of distance (*𝓁*) between pairs of fourfold degenerate synonymous sites in the same core gene in *B. vulgatus* (SI Section S6 Text). Individual data points are shown for distances < 100bp, while the solid line shows the average in sliding windows of 0.2 log units. The grey line indicates the values obtained without controlling for population structure, while the blue line is restricted to QP hosts in the largest top-level clade (Table S2, Text S5.2). The solid black line denotes the neutral prediction from SI Section S6 Text; the two free parameters in this model are 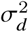 and *𝓁* scaling factors, which are shifted to enhance visibility. For comparison, the core-genome-wide estimate for SNVs in different genes is depicted by the dashed line and circle. (c) Summary of linkage disequilibrium for QP hosts in the largest top-level clade for all species with ≥ 10 QP hosts, sorted phylogenetically as in Fig. 2B. For each species, the three dashes denote the value of 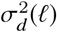 for intragenic distances of *𝓁* = 9, 99, and 2001 bp, respectively, while the core-genome-wide values are depicted by circles. Points belonging to the same species are connected by vertical lines for visualization.

An illustrative example is provided by *Bacteroides vulgatus*, one of the most abundant and prevalent species in our cohort. At the highest divergence values, we observe little phylogenetic inconsistency for this species (blue line in Fig. 4A), consistent with the strong population structure suggested by the multi-modal divergence distribution in Fig. 2B. For intermediate values of divergence, in contrast, we find that a large majority of all SNVs are inconsistent with the genome-wide divergence estimates. Similarly high values of phylogenetic inconsistency are observed in most of the other species as well (light red lines in Fig. 4A).

While these signals are suggestive of recombination, phylogenetic inconsistencies can also arise from purely clonal mechanisms (e.g. recurrent mutation), or from statistical uncertainties in the genome-wide tree. We therefore sought additional evidence of recombination by examining how phylogenetic inconsistency varies for pairs of SNVs in different locations in the genome. We quantified phylogenetic inconsistency between pairs of SNVs using a standard measure of linkage disequilibrium (LD), 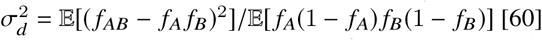, with an unbiased estimator to control for varying sample size (SI Section S6 Text). The overall magnitude of 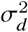 is often uninformative, since it depends on demographic factors, the extent of recurrent mutation, etc. However, relative differences in 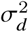 between different pairs of SNVs reflect differences in the effective recombination rate. If 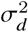 consistently decreases for SNVs that are separated by greater genomic distances, then we can conclude that recombination, rather than recurrent mutation, is responsible for the phylogenetic inconsistency that we observe [61].

With traditional metagenomic approaches, it is difficult to measure 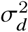 unless the SNVs co-occur on the same sequencing read. By focusing on QP samples, we can now estimate 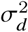 between SNVs that are separated by greater distances along the reference genome. However, since the synteny of individual lineages may differ substantially from the reference genome, we only assigned coordinate distances (*𝓁*) to pairs of SNVs in the same gene, which are more likely (but not guaranteed) to be nearby in the genomes in other samples; all other pairs of SNVs are grouped together in a single category (“core-genome-wide”). We then estimated 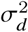 as a function of *𝓁* for each of these distance categories (SI Section S6 Text), and analyzed the shape of this function.

As an example, Fig. 4B illustrates the estimated values of σ^2^ (*𝓁*) for synonymous SNVs in the core genome of *B. vulgatus*. Similar curves are shown for several other species in Fig. S12. As anticipated by our analysis in Fig. 4A, it is crucial to account for the presence of strong population structure. The LD curve among all samples decays only slightly with distance, as expected from a mixture of genetically isolated sub-populations. However, if we restrict our attention to the lineages in the largest sub-population, we observe a pronounced decay in 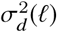. To account for these confounding effects, we manually annotated top level clades for each species using the genome-wide divergence distribution (Text S5.2), using standard criteria for identifying ecotype clusters [62].

In Fig. 4C, we plot summarized versions of the σ^2^ (*𝓁*) curves across a panel of ∼ 40 prevalent species. In almost all cases, we find that core-genome-wide LD is significantly lower than for pairs of SNVs in the same core gene, suggesting that much of the phylogenetic inconsistency in Fig. 2 is caused by recombination. Qualitatively similar results are obtained if we repeat our analysis using isolate genomes from some of the more well-characterized species (Fig. S13, SI Section S7 Text). In principle, signatures of recombination between genes could be driven by the exchange of intact operons or other large clusters of genes (e.g. on an extra-chromosomal plasmid). However, Figs. 4 and S13 also show a significant decay in LD within individual genes, suggesting a role for homologous recombination within genes as well.

The magnitude of the decay of LD within core genes is somewhat less than has been observed in other bacterial species [16], and only rarely decays to genome-wide levels by the end of a typical gene. Moreover, by visualizing the data on a logarithmic scale, we see that the shape of 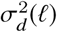 is inconsistent with the predictions of the neutral model (Fig. 4A), decaying much more slowly with *𝓁* than the ∼1/*𝓁* dependence expected at large distances [60]. Thus, while we can obtain rough estimates of *r*/µ by fitting the data to a neutral model (which generally support 0.1; ≲ *r*/µ ≲ 10, see Fig. S14), these estimates should be regarded with caution because they vary depending on the length scale on which they are measured (SI Section S6 Text). This suggests that new theoretical models will be required to fully understand the patterns of recombination that we observe.

### Short-term succession within hosts

So far, we have focused on evolutionary changes that accumulate over many host colonization cycles. In principle, evolutionary changes can also accumulate within hosts over time. Longitudinal studies have shown that strains and metagenomes sampled from the same host are more similar to each other on average than to samples from different hosts [31, 33, 35, 44, 63, 64]. This suggests that resident populations of bacteria persist within hosts for at least a year (∼300 -3000 generations), which is potentially enough time for evolutionary adaptation to occur [7]. However, the limited resolution of previous polymorphism- [31] or consensus-based comparisons [35, 44] has made it difficult to quantify the individual changes that accumulate within hosts, and to interpret these changes in an evolutionary context.

#### Within-host dynamics reflect a mixture of replacement and modification

To address this issue, we focused on the species in longitudinally sampled HMP subjects that were quasi-phaseable at consecutive timepoints. This yields a total of 801 *resident populations* (host/species timepoint pairs) across 45 of the most prevalent species (Fig. S5). Our calculations show that the false positives caused by sampling noise should be sufficiently rare that we can resolve a single nucleotide difference between two of these timepoints in a genome-wide scan (Text S3.4). In contrast to existing reference-based approaches, we have also imposed additional filters to minimize false positives from mapping artifacts (Text S1 Text).

We first examined the SNV differences that accumulated within each *resident population* (i.e., a particular host/species pair) over time. We considered SNVs in both core and accessory genes on the reference genome, since the latter are plausibly enriched for host-specific targets of selection [65]. Consistent with previous work [31, 44], the average number of within-host differences is ∼ 100-fold smaller than the average number of differences between unrelated hosts (Fig. S15). However, the within-host changes are distributed across the different resident populations in a highly skewed manner (Figs. 5 and S16). Visualized on a logarithmic scale, the data reveal a striking multi-modal pattern, suggesting that the within-host differences arise from two separate processes.

**Fig 5.**
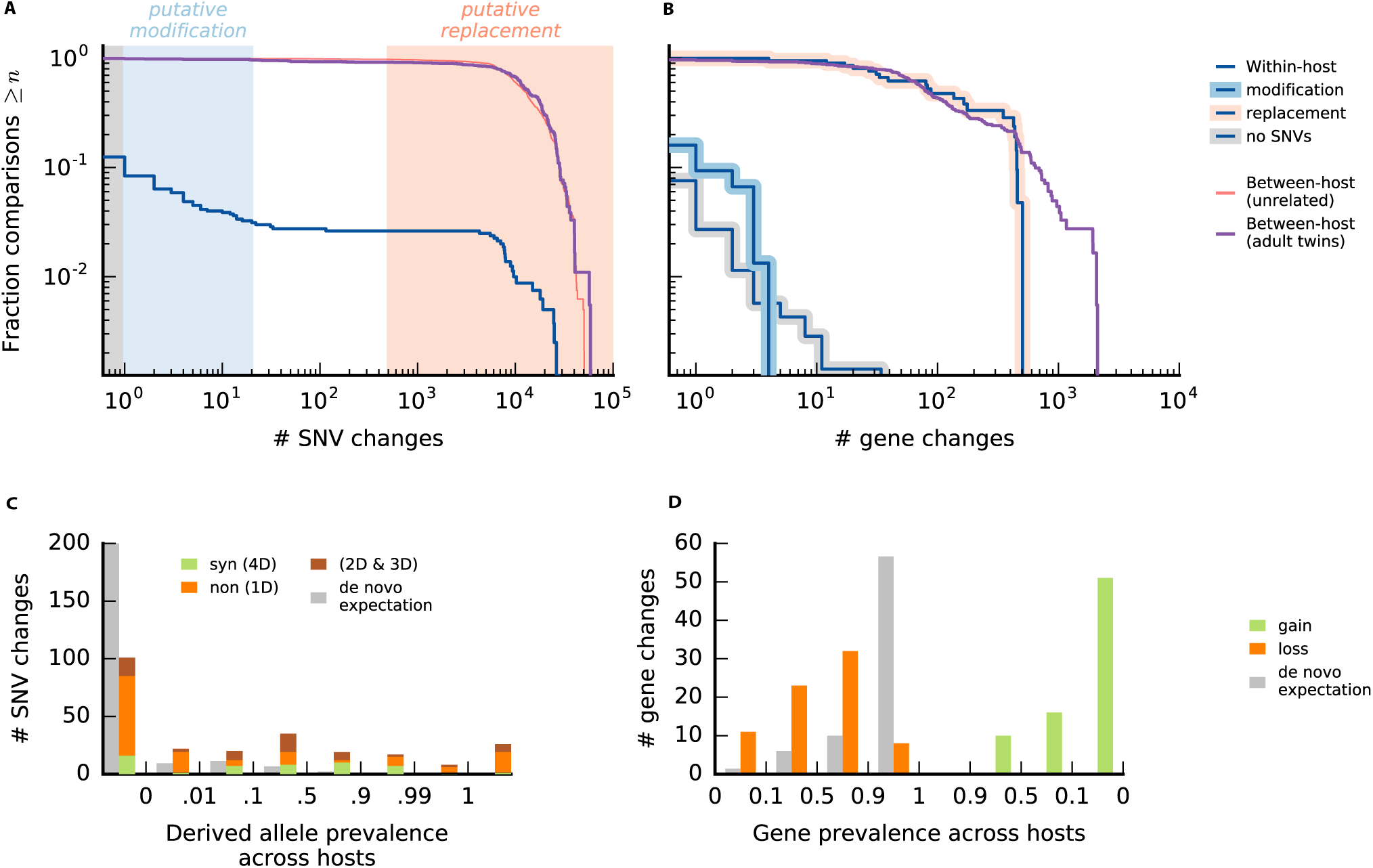
Within-host changes across prevalent species of gut bacteria. (a) Within-host nucleotide differences over ∼6 months. The blue line shows the distribution of the number of SNV differences between consecutive QP timepoints for different combinations of species, host, and non-overlapping time interval (if more than two samples are available), for the 45 prevalent species in Fig. S16. For comparison, the red line shows a matched distribution of the number of SNV differences between each initial timepoint and a randomly selected HMP host. Finally, the purple line shows the distribution of the number of SNV differences between QP lineages in pairs of adult twins. The shaded regions indicate replacement events (light red), modification events (light blue), and no detected changes (grey); these ad-hoc thresholds were chosen to be conservative in calling modifications. (b) Within-host gene content differences (gains+losses). The blue lines show the distribution of the number of gene content differences within hosts for the samples in (a), with the putative modifications highlighted in light blue, the putative replacements highlighted in light red, and the samples with no SNV changes highlighted in grey. For comparison, the corresponding between-host and twin distributions are shown as in (a). (c) The total number of nucleotide differences at non-degenerate nonsynonymous sites (1D), fourfold degenerate synonymous sites (4D), and other sites (2D and 3D) aggregated across the modification events in (a). Sites are stratified based on their prevalence across hosts (SI Section S8 Text). For comparison, the grey bars indicate the expected distribution for random *de novo* mutations (Text S8.1). (d) The total number of gene loss and gain events among the gene content differences in (b), stratified by the prevalence of the gene across hosts. The *de novo* expectation for gene losses is computed as in (c); by definition, there are no *de novo* gene gains.

Most of the resident populations did not have any detectable SNV differences over the ∼6 month sampling window (i.e., the median is zero). Yet in a small minority of cases (∼3%), the resident populations accumulated several thousand mutations, comparable to the typical number of differences between hosts (Fig. 5A). This is consistent with previous notions of strain *replacement* [35], in which the dominant resident strain is succeeded by an effectively unrelated strain from the larger metapopulation. This operational definition includes both the invasion of a new strain (e.g. from other hosts or body sites), or a sudden rise in frequency of a previously-colonized strain that had been segregating at low frequency.

In addition to rare replacement events, a larger fraction of resident populations in Fig. 5A (∼10% of the total) have a moderate number of SNV differences (on the order of 20 or fewer). We will refer to these as *modification* events, in order to distinguish them from the replacement events above. In contrast to replacements, modifications preserve most of the genetic information in a lineage when a new genetic change is added. This is true at the level of nucleotide divergence, but also for gene content (Fig. 5B) and the sharing of private marker SNVs (Fig. S8B). We therefore hypothesize that the modification events in Fig. 5 reflect heritable evolutionary changes that have risen to high frequency within the host.

Given the large census population sizes in the gut [66], we conclude that these rapid allele frequency changes must be driven by natural selection, rather than genetic drift. However, this does not imply that the observed SNVs are the direct target of selection: given the limitations of our reference-based approach, and our aggressive filtering scheme, the observed mutations may simply be passengers hitchhiking alongside an unseen selected locus. In either case, given the size of the frequency change (Δ*f* ∼0.5) and the length of the sampling period (Δ*t* ∼6 months), we infer that the selected haplotype must have had an average fitness benefit of at least *S* ∼1% per day.

To further probe the dynamics of within-host evolution, we pooled the 248 SNV differences observed across the 75 modification events in our cohort, and we stratified them according to two additional criteria. We first partitioned the SNVs according to how prevalent the sweeping allele was among the other hosts in our cohort (Figs. 5C and S17A). By comparing this distribution against the null expectation for randomly selected sites, we find that there are significantly more intermediate- and high-prevalence mutations than expected for random *de novo* mutations (*P* < 10^-4^, Text S8.1).

One potential explanation for this signal could be parallel evolution [67], e.g. if the same strongly beneficial mutations independently arose and fixed in different hosts. However, we can rule out this recurrent sweep hypothesis by further partitioning the SNVs into synonymous and nonsynonymous mutations (Fig. 5C). The relative fractions of the two types are distributed across the different prevalence classes in a highly nonuniform manner (*P* < 10^-4^, Text S8.2). Among rare alleles (< 1% prevalence), we observe an excess of nonsynonymous mutations [*dN*/*dS* ≈1.3 (0.8, 2.4)], consistent with positive selection and hitchhiking. By contrast, nonsynonymous mutations are depleted and synonymous mutations enriched for alleles with intermediate prevalence (0.1 < *f* < 0.9), precisely where the recurrent sweep hypothesis requires the strongest selection pressures. This low value is surprising even for pure passenger mutations, since purifying selection should be rendered inefficient over these short timescales [68], similar to what we observed in Fig. 3.

Together, these observations suggest an alternate hypothesis, in which some of the within-host sweeps are driven by much older DNA fragments that were acquired through recombination. This could explain the intermediate prevalence of some sweeping alleles, since standing variants can arrive through recombination. And it can simultaneously explain their low *dN*/*dS* values, since there is more time for deleterious mutations to be purged (and for synonymous mutations to accumulate) before the fragment is transferred.

Consistent with this hypothesis, we also found evidence for a small number of gene content differences between the two timepoints in many of the non-replacement samples (Fig. 5B). Gene content differences were twice as likely to occur in populations where we observed one or more SNV differences (*P* ≈ 0.025, Fisher’s exact test), though the overall rates are still modest under our current filtering criteria (∼10%). We observed a roughly equal contribution from gains and losses (Fig. 5D). The gene losses could be consistent with purely clonal processes (e.g. a large deletion mutation) as well as recombination (e.g. if the incorporated homologous fragment lacks the gene in question). Gene gains, on the other hand, can only be acquired through recombination. The genes that are gained and lost tend to be drawn from the accessory portion of the genome (Figs. 5D and S17C), consistent with the expectation that these genes are more likely to be gained or lost over time.

#### Replacement dominates over longer within-host timescales

The successional dynamics in the HMP cohort raise a number of key questions about how these dynamics play out over longer timescales. For example, does the probability of a replacement accumulate uniformly with time, so that we would expect most strains to be replaced after ∼20 years? Or are replacements concentrated in a few replacement-prone individuals, with a negligible rate among the larger population? Alternatively, do resident populations eventually acquire enough evolutionary changes that they become so well-adapted the host that replacements become less likely to succeed?

To fully address these questions, we would require a large longitudinal cohort with metagenomes collected over a period of decades. However, we can approximate this design in a crude way by comparing metagenomes collected from a cohort of ∼200 adult twins from the TwinsUK project [45]. Previous work has shown that twins are often colonized by nearly identical strains in early childhood [46] (Fig. S18). By comparing quasi-phasable samples in adult twins, we can gain insight into the changes that have occurred in the ∼20 -40 years that the hosts have spent in separate households.

The number of SNV and gene changes between the resident populations in each twin pair are illustrated in Fig. 5A,B. We observe striking departures from the within-host distribution: while ∼3% of the resident populations experienced a replacement event in the ∼6 month HMP study, more than 90% of the resident populations in twins have more than ∼1000 SNV differences between them. Compared to the modification events we observed in the HMP study, these highly diverged twin strains have much lower rates of private marker SNV sharing (Fig. S8), along with a higher proportion of SNVs with intermediate prevalence (Fig. S19). Together, these lines of evidence suggest that the highly diverged strains in twins are true replacement events, rather than an accumulation of many evolutionary changes. The 16 resident populations with fewer than 1000 SNV differences were scattered across 13 twin pairs. All had at least one SNV or gene difference between the twins (median 29 SNVs and 1.5 genes), which is significantly higher than the within-host distribution from the HMP cohort. However, a larger sample size is required to determine what fraction of these SNVs accumulated since colonization.

Together, these data suggest that a vast majority of the resident populations have experienced a replacement over the ∼20 -40 years that their hosts have spent in different households. This observation is consistent with a straightforward extrapolation of the short-term estimates from the HMP cohort, which predicts that replacement should dominate over modification in a typical population after ∼20 years. In other words, replacement is not confined to a few special hosts, but will eventually occur for most (Western) individuals given enough time. This suggests that the potential benefits of local adaptation do not compound indefinitely.

The high prevalence of twin replacements also provides insight into the two replacement mechanisms described in the previous section. If replacements are primarily drawn from a set of strains that colonized both twins during childhood, then the replacement probability should saturate at 1 – 1/*n*_*c*_, where *n*_*c*_ is the number of colonizing strains. The observed replacement probability of 90% would then imply that the number of low frequency colonizing strains for each species must be as large as *n*_*c*_ ∼10, or that most of the replacements are caused by the invasion of new strains that arrive after initial colonization. It will be interesting to test these alternative mechanisms with deeper sequencing and longer time courses.

## Discussion

Evolutionary processes can play an important role in many microbial communities. Yet despite increasing amounts of sequence data, our understanding of these processes is often limited by our ability to resolve evolutionary changes in populations from complex communities. In this work, we quantify the evolutionary forces that operate within bacteria in the human gut microbiome by characterizing in detail the lineage structure of ∼30 species in metagenomic samples from individual hosts.

Building on previous work by Ref. [35] and others, we found that the within-host lineage structure of many prevalent species is consistent with colonization by a few distinct strains from the larger population, with the identities and frequencies of these strains varying from person to person (Fig. 1). The distribution of strain frequencies in this “oligo-colonization model” is itself quite interesting: in the absence of fine tuning, it is not clear what mechanisms would allow for a second or third strain to reach intermediate frequency, while preventing a large number of other lineages from entering and growing to detectable levels at the same time. A better understanding of the colonization process, and how it might vary among the species in Fig. 1F, is an important avenue for future work.

Given the wide variation among species and hosts, we chose to focus on a subset of samples with particularly simple strain mixtures for a given species, in which we can resolve evolutionary changes in the dominant lineage with a high degree of confidence. Our quasi-phasing approach can be viewed as a refinement of the consensus approximation employed in earlier studies [4, 35, 37, 38], but with more quantitative estimates of the errors associated with detecting genetic differences between lineages in different samples.

By analyzing genetic differences between lineages in separate hosts, we found that long-term evolutionary dynamics in many gut bacteria are consistent with quasi-sexual evolution and purifying selection, with relatively weak geographic structure. Earlier work had documented extensive horizontal transfer between distantly related species in the gut [69, 70], but our ability to estimate rates of recombination within species was previously limited by the small number of sequenced isolates for many species of gut bacteria [71]. The high rates of homologous recombination we observed with our quasi-phasing approach are qualitatively consistent with previous observations in other bacterial species [16, 71–75]. However, our quantitative characterization of linkage disequilibrium revealed interesting departures from the standard neutral prediction that cannot be captured by simply lowering the recombination rate. Understanding the origin of this discrepancy is an interesting topic for future work. It is also interesting to ask how these long-term rates of recombination could emerge from the oligo-colonization model above, since it would seem to limit opportunities for genetic exchange among strains of the same species.

In a complex community like the gut, a key advantage of our metagenomic approach is that it can jointly measure genetic differences in multiple species for the same pair of hosts. By leveraging this feature, we found that previous observations of highly similar strains in different hosts [35, 44] are not driven by cryptic host relatedness. Instead, the presence of these closely related strains, and the genetic differences that accumulate between them, may be driven by more general population genetic processes in bacteria that operate on timescales much shorter than the typical coalescent time across hosts. It is difficult to produce such closely related strains in traditional population genetic models of loosely linked loci [76] (or “bags of genes” [77]), though recent hybrid models of vertical and horizontal inheritance [75, 78] or fine-scale ecotype structure [79] could potentially provide an explanation for this effect. Further characterization of these short-term evolutionary processes will be vital for current efforts to quantify strain sharing across hosts [33, 46, 56], which often require implicit assumptions about how genetic changes accumulate on short timescales. Our results suggest that these short-term dynamics of across-host evolution may not be easily extrapolated by comparing average pairs of strains.

The other main advantage of our quasi-phasing approach is its ability to resolve a small number of evolutionary changes that could accumulate within hosts over short timescales. Previous work has shown that on average, longitudinally sampled metagenomes from the same host are more similar to each other than metagenomes from different hosts [31, 33, 63, 64], and that some within-host changes can be ascribed to replacement by distantly related strains [35, 44]. However, the limited resolution of previous polymorphism- [31] or consensus-based comparisons [35, 44] had made it difficult to determine whether resident strains also evolve over time.

Our quasi-phasing approach overcomes this limitation, enabling finely resolved estimates of temporal change within individual species in individual hosts. This increased resolution revealed an additional category of within-host variation, which we have termed *modification*, in which resident strains acquire modest numbers of SNV and gene changes over time. This broad range of outcomes shows why it is essential to understand the distribution of temporal variation across hosts: even though modification events were ∼3*x* more common than replacements in our cohort, their contributions to the total genetic differences are quickly diluted as soon as a single replacement is included (Fig. S15). As a result, we expect that previous metagenome-wide [31] or species-averaged [44] estimates of longitudinal variation largely reflect the rates and genetic differences associated with replacement events, rather than evolutionary changes.

Though we have interpreted modifications as evolutionary events (i.e., mutations to an existing genome), it is possible that they could also reflect replacement by extremely closely related strains, as in Fig. 2. The present data seem to argue against this scenario: modifications are not only associated with different patterns of SNV sharing (Fig. S8), but we also observe significant asymmetries in the prevalence distributions in Fig. 5C,D that depend on the temporal ordering of the two samples (see Text S8.3). This temporal directionality is a natural product of evolution, but it is less likely to emerge from competition between a fixed set of strains. Unambiguous proof of evolution could also be observed in a longer time course, since subsequent evolutionary changes should eventually accumulate in the background of earlier substitutions. Further investigation of these nested substitutions remains an interesting topic for future work.

The signatures of the sweeping SNVs, along with the presence of gene gain events, suggest that some of the within-host sweeps we observed were seeded by recombination, rather than de novo mutation. In particular, many of the alleles that swept within hosts were also present in many other hosts, yet their *dN*/*dS* values indicated strong purifying selection, consistent with an ancient polymorphism (Fig. 3). Sweeps of private SNVs, by contrast, were associated with a much higher fraction of nonsynonymous mutations, consistent with adaptive de novo evolution. Interestingly, we also observe a slight excess of *private* nonsynonymous mutations between closely related strains in different hosts (Fig. S9). This suggests that some of the differences observed between hosts may reflect a record of recent within-host adaptation.

The fact that some sweeps are seeded by recombination stands in contrast to the *de novo* mutations observed in microbial evolution experiments [13] and some within-host pathogens [21, 22]. Yet in hindsight, it is easy to see why recombination could be a more efficient route to adaptation in a complex ecosystem like the gut microbiome, given the large strain diversity [42], the high rates of DNA exchange [69, 70], and the potentially larger selective advantage of importing an existing functional unit that has already been optimized by natural selection [11]. Consistent with this hypothesis, adaptive introgression events have also been observed on slightly longer timescales in bacterial biofilms from an acid mine drainage system [14], and they are an important force in the evolution of virulence and antibiotic resistance in clinical settings [80].

While the data suggest that some within-host sweeps are initiated by a recombination event, it is less clear whether recombination is relevant during the sweep itself. Given the short timescales involved, and our estimates of the recombination rate (Fig. S14), we would expect many of the observed sweeps to proceed in an essentially clonal fashion, since recombination would have little time to break up a megabase-sized genome. If this were the case, it would provide many opportunities for substantially deleterious mutations (with fitness costs of order *S*_*d*_ ∼1% per day) to hitchhike to high frequencies within hosts [68], thereby limiting the ability of bacteria to optimize to their local environment. The typical fitness costs inferred from Fig. 2D lie far below this threshold, and would therefore be difficult to purge within individual hosts. In this scenario, the low values of *d*_*N*_/*d*_*S*_ observed between hosts (as well as the putative introgression events) would crucially rely on the competition process across hosts [81]. Although the baseline recombination rates suggest clonal sweeps, there are also other vectors of exchange (e.g. transposons, prophage, etc.) with much higher rates of recombination. Such mechanisms could allow within-host sweeps to behave in a quasi-sexual fashion, preserving genetic diversity elsewhere in the genome. These “local” sweeps are predicted in certain theoretical models [82,83] and have been observed in a few other bacterial systems [15, 17, 84]. If local sweeps were also a common mode of adaptation in the gut microbiome, they would allow bacteria to purge deleterious mutations more efficiently than in the clonal scenario above.

Although evolution was more common than replacement on ∼6 month timescales, our analysis of adult twins suggest that the rare replacement events eventually dominate on multi-decade timescales. This suggests that resident strains are limited in their ability to evolve to become hyper-adapted to their host, since most strains were eventually susceptible to replacement. Though our results indicate that the long-term probability of replacement is largely uniform across hosts, it remains an open question whether these events occur more or less uniformly in time, or whether they occur in punctuated bursts during major ecosystem perturbations (e.g. antibiotic treatment). This would be an interesting question to address with denser and longer time-series data.

While we have identified many interesting signatures of within-host adaptation, there are several important limitations to our analysis. One class concerns the events that we cannot see with our approach (i.e., *false negatives*). For example, our reference-based method only tracks SNVs and gene copy numbers in the genomes of previously sequenced isolates of a given species. Within this subset, we have also imposed a number of stringent bioinformatic filters, further limiting the sequence space that we consider. Thus, it is likely that we are missing many of the true targets of selection, which might be expected to be concentrated in the host-specific portion of the microbiome, multi-copy gene families, or in genes that are shared across multiple prevalent species. A further limitation is that we can only analyze the evolutionary dynamics of QP samples (though the consistency of our results for species with different QP fractions suggests that this might not be a major issue). Finally, a potentially more important false negative is that our current method can only identify complete or nearly complete sweeps within individual hosts. While we observed many within-host changes that matched this criterion, we may be missing many other examples of within-host adaptation where variants do not completely fix. Given the large population sizes involved, such sweeps can naturally arise from phenotypically identical mutations at multiple genetic loci [67, 85], or through additional ecotype partitioning between the lineages of a given species [23, 25]. Both mechanisms have been observed in experimental populations of *E. coli* adapting to a model mouse microbiome [7].

In addition to these false negatives, the other limitation of our approach concerns potential false positives inherent in any metagenomic analysis. With short-read data, it is difficult to truly know whether a paticular DNA fragment is linked to a particular species, or whether it resides in the genome of another species (perhaps an uncultured one), that is fluctuating in abundance. False SNV and gene changes can therefore occur due to these *read donating* effects. The temporally asymmetric prevalence distributions in Fig. 5C,D suggest that our filters were successful in eliminating many of these events (Text S8.3). However, isolate or long-read sequences are required to unambiguously prove that these variants are linked to the population of interest.

Fortunately, two concurrent studies have also documented short-term evolution of gut bacteria within healthy human hosts using an isolate-based approach [86, 87]. Each study focused on a single bacterial species, *E. coli* in Ref. [86] and *B. fragilis* in Ref. [87]. Although *E. coli* was not sufficiently abundant in our cohort to be included in our within-host analysis, the observations in *B. fragilis* are largely consistent with our findings that within-host evolution can be rapid, and that it can be mediated by recombination in addition to new mutations. Crucially, since these observations were obtained using an isolate-based approach, they are not subject to the same methodological limitations described above, and they therefore serve as an independent verification of our results. However, since our statistical approach provides simultaneous observations across more than ∼30 prevalent species, our results show that these general patterns of within-host evolution are shared across many species of gut bacteria, and they demonstrate a general approach for investigating these forces in widely available metagenomic data. Future efforts to combine metagenomic- and isolate-based approaches, e.g. by incorporating long-range linkage information [41, 88, 89], will be crucial for building a more detailed understanding of these evolutionary processes.

## Supporting Information

**S1 Fig. Schematic depiction of phasing and substitution errors caused by sampling noise.**

**S2 Fig. Average genetic distance between *B. vulgatus* metagenomes.**

**S3 Fig. Correlation between within-host diversity and the fraction of non-QP samples per species.**

**S4 Fig. Distribution of the number of QP species per sample.**

**S5 Fig. Distribution of quasi-phaseable (QP) samples in longitudinal samples and adult twin pairs.**

**S6 Fig. Distribution of estimated gene copy numbers for the four samples in Fig. 1.**

**S7 Fig. SNV and gene content differences among closely related strains.**

**S8 Fig. Private marker SNV sharing within and between hosts.**

**S9 Fig. Signatures of selective constraint within private SNVs.**

**S10 Fig. Schematic illustration of phylogenetic inconsistency between individual SNVs and core-genome-wide divergence.**

**S11 Fig. Top-level clade structure among lineages in different QP hosts.**

**S12 Fig. Decay of linkage disequilibrium in three example species.**

**S13 Fig. Recapitulating patterns of between-host evolution from sequenced isolates.**

**S14 Fig. Recombination rate estimates based on the decay of linkage disequilibrium.**

**S15 Fig. Average number of SNV differences within and between hosts.**

**S16 Fig. Comparable rates of within-host SNV and gene changes across prevalent species.**

**S17 Fig. Prevalence distributions of within-host SNV and gene content differences without binning.**

**S18 Fig. SNV and gene content differences between younger twins.**

**S19 Fig. Prevalence of SNV and gene content differences between adult twins.**

**S1 Table. Metagenomic samples used in study.**

**S2 Table. Top-level clade definitions for prevalent species. S1 Text. Metagenomic pipeline.**

**S2 Text. Quantifying within-species diversity in individual samples.**

**S3 Text. Quasi-phasing metagenomic samples.**

**S4 Text. Population genetic null model of purifying selection for pairwise divergence across hosts.**

**S5 Text. Phylogenetic inconsistency and clade structure across hosts.**

**S6 Text. Population genetic null model for the decay of linkage disequilibrium.**

**S7 Text. Validation of between-host patterns using isolate sequences.**

**S8 Text. Quantifying prevalence of within-host SNV and gene changes.**

## Data and code availability

The raw sequencing reads for the metagenomic samples used in this study were downloaded from public repositories listed in Refs. [42–45]. All necessary metadata, as well as the source code for the sequencing pipeline, downstream analyses, and figure generation, will available at GitHub (https://github.com/benjaminhgood/microbiome_evolution) upon publication.

## Acknowledgments

We thank S. Wyman for assistance with sample metadata and S. Greenblum, S. Venkataram, A. Harpak, J. Ladau, I. Cvijovic, Tami Lieberman, Joshua Plotkin and members of the Pollard, Hallatschek, and Petrov labs for feedback. This research was supported in part by the National Science Foundation (PHY-1125915 and DMS-1563159), the National Institutes for Health (R25GM067110), and the Gordon and Betty Moore Foundation (No. 2919.01). BHG acknowledges support from the Miller Institute for Basic Research in Science at the University of California Berkeley. OH acknowledges support from a National Science Foundation Career Award (No. 1555330) and a Simons Investigator award from the Simons Foundation (No. 327934). KSP acknowledges support from Gladstone Institutes, Chan-Zuckerberg Biohub, and the San Simeon Fund.

---

**Table S1. Metagenomic samples used in study.** We analyzed a total of 1013 samples from 693 individuals. This included samples from 250 individuals from the Human Microbiome Project (HMP) [42, 44], 250 individuals from Ref. [45], 185 individuals from Ref. [43], and 8 individuals from Ref. [46]. Listed are the subject ids, sample ids, run accessions, country of the study, continent of the study, visit number, and study (HMP, Xie *et al*, Korpela *et al*, or Qin *et al*, 2012).

**Table S2. Top-level clade definitions.** This table contains the manually-defined top-level clades described in Text S5.2. Rows list the various combinations of species and hosts plotted in Fig. 2, along with its corresponding numeric clade label.

## S1 Text

### Metagenomic pipeline

#### S1.1 Overview

We analyzed whole-genome sequence data from a panel of stool samples from 693 healthy human subjects (Table S1). As described in the main text, this panel includes 250 North American subjects sequenced by the Human Microbiome Project [42, 44], a subset of which were sampled at 2 or 3 timepoints roughly 6-12 months apart. We also included a cohort of 125 pairs of adult twins from the TwinsUK registry [45], 4 pairs of younger twins from Ref. [46], and 185 Chinese subjects sequenced in Ref. [43].

Previous work has shown that there is little genomic variability between technical and sample replicates in HMP data [33, 44], so we merged fastq files for technical and sample replicates from the same time point to increase coverage to resolve within-host allele frequencies. We analyzed the gene and SNV content of these samples using the MIDAS software package [v1.2.2 [33]] as a foundation, with multiple additional layers of filtering implemented in custom postprocessing scripts, described below. This postprocessing pipeline was designed to be as inclusive as possible in the early steps, when hard thresholds are required, so that we can adaptively estimate thresholds from the data to use in later postprocessing steps. Later rounds of postprocessing impose a set of progressively more conservative filters, which are designed to rule out mapping artifacts and other metagenomic ambiguities, at the expense of reduced genome and species coverage. We ultimately apply this pipeline to estimate SNV and gene content changes in species with sequencing coverage of 20x or more, so our filters are designed with these numbers in mind.

#### S1.2 Estimating the panel of reference species for each host

The first step in the pipeline is to determine which species to include in the personalized reference panel for each host. The goal is to include as many truly present species as possible (to prevent their reads from being *donated* to other reference genomes) while leaving out species that are truly absent (to prevent their reference genomes from *stealing* reads from other species). To determine this set, MIDAS first quantifies the relative abundances of species in different metagenomic samples by mapping sequencing reads to a database of universal single-copy “marker” gene sequences for each of the species in the default MIDAS database (version 1.2, downloaded on November 21, 2016 [33]). We include a species in the reference panel for a given sample if it has an average marker gene coverage ≥ 3 in that sample. This definition leaves out many species that are present at lower abundances. We note, however, that their coverage would be too low for them to be included in our downstream analyses, and any donated reads would add only fractional contributions to the polymorphism frequencies for species in our target coverage range.

For longitudinally sampled individuals, we defined a single reference panel for each host by including all species with marker coverage ≥ 3 in at least one timepoint. This choice is designed to reduce potential mapping artifacts by ensuring that all longitudinal comparisons are performed with the same mapping parameters.

#### S1.3 Quantifying gene content

We next quantified the gene content for each species present in each sample. In downstream analyses, gene content information was used to estimate the prevalence of genes in the broader population, and to quantify gene content differences between QP samples (SI Section S3.5).

MIDAS estimates gene copy number for each species by mapping reads to a database of gene families (or *pangenome*) constructed from genes in sequenced isolates [33]. This approach has been adopted in a number of related methods [90, 91], along with similar methods based on co-occurence or binning [43, 92, 93]. Briefly, pre-computed pangenomes are supplied for each species in the default MIDAS database, and a host-specific database is constructed by concatenating pangenomes from each species in the personalized reference panel. Sequencing reads are aligned to this host-specific database using Bowtie2 [94] with default MIDAS settings (local alignment, MAPID ≥ 94.0%, READQ ≥ 20, and ALN_COV ≥ 0.75), and the average coverage is estimated by dividing the total number of mapped reads in a gene family by the total target size. We note that with these settings, reads with multiple best-hit alignments will be distributed among these targets according to their proportional representation on the pangenome reference sequence; these reads were retained to ensure consistent estimates of average coverage within a gene family, which might contain multiple highly similar genes [33].

For each species, average coverage was reported for each gene, as well as for a panel of universal, single-copy marker genes [33, 95]. The copynumber of a gene (*c*) is then estimated as the ratio between its coverage and the (median) marker gene coverage. We used these raw copynumber values to estimate the prevalence of genes in the broader population, defined as the fraction of samples with *c* ≥ 0.3 (conditioned on the marker coverage being ≥5x). Ref. [33] have previously shown that these thresholds yield accurate gene presence estimates. We then used these prevalence estimates to define a *core genome* for each species, defined as the set of genes with prevalence ≥ 0.9.

In addition to quantifying gene prevalence, we also used MIDAS’s copynumber estimates to detect changes in gene content between QP samples (SI Section S3.5). The QP methodology was designed to eliminate spurious gene content differences that arise from sampling noise, e.g. when a host is colonized by multiple strains of the same species. However, another well-known limitation of the pangenome approach used by MIDAS and others is that linkage between a gene and its species is not observed directly, but is only inferred by the presence of that gene in a previously sequenced isolate. This can lead to spurious copynumber changes if a target gene is actually linked to a different species in a particular host, and the relative abundance of the species are simply changing over time. To guard against this scenario, we implemented a number of additional filters described below.

First, we only considered gene content differences that were consistent with a single copy gene transitioning to zero copynumber, or vice versa. We used a permissive definition of potential single copy genes (0.6 ≤ *c* ≤ 1.2, with marker coverage ≥ 20*x*) in order to capture normal coverage variation along the genome in growing cells [96], see Fig. S6. Similarly, we defined a zero copynumber to be *c* ≤ 0.05, so that a small fraction of cells could still retain the gene. (For simplicity, these copynumber thresholds are also used for the sampling error calculation in SI Section S3.5.) We implemented this copynumber restriction because, if a gene is truly linked to a different species, it is less likely to have both a “normal” and “absent” copynumber by chance. For this to happen, it would require that the two species that share the gene in a given host (a rare event) have similar relative abundance at one timepoint and ≥10-fold different abundance at the other (another rare event). Although this approach omits many biologically interesting copynumber differences among multi-copy genes (e.g. transporter genes in *Bacteroides* [97]), we do not study them here because they are much harder to disentangle from mapping artifacts.

To supplement these copynumber filters, we also created a blacklist of genes that are potentially shared across species. This is helpful for some highly promiscuous genes, e.g. transposons in *Bacteroides* [98], where the probability of cross-species sharing cannot be assumed to be low. We constructed this blacklist by searching for gene families in the MIDAS database that had sequence similarity ≥ 95% with a gene family in another species (SI Section S1.5). These families constitute a gold standard for gene sharing events, since they imply that highly similar genes have been observed in isolates from different species. However, this approach can also miss cases of gene sharing for species with poor phylogenetic coverage in the MIDAS isolate database. We therefore supplemented the isolate-based blacklist with gene sharing candidates that were identified directly from the metagenomic data. In particular, we defined a putatively shared gene to be one with c ≥ 3 in at least one sample in our cohort, since this could indicate read donating by a shared gene in a more abundant species. This does not constitute proof of gene sharing, but it is conservative for the purposes of constructing a blacklist. The metagenomic and isolate-based methods identified many common gene sharing candidates, but for many species, there were also many genes that were only identified by one of the two approaches.

All genes in the combined blacklist were excluded from downstream analyses of gene content estimation and SNV calling. As with our copynumber filters above, this likely omits many biologically interesting regions of the genome, since shared genes are arguably more likely to play a role in short-term evolutionary dynamics. We ignore them here in order to minimize false positives created by read donating.

Even with these various filters, it is important to note that the pangenome approach employed by us and others is at best an inferential method, which relies on an out-of-sample estimate of linkage to the correct species background. While we have included these gene content differences to supplement our SNV-based analysis, isolate sequences [98] or long read data [41] are required to definitely prove that any specific gene content difference is linked to the species of interest.

#### S1.4 Quantifying SNVs

We next quantified single nucleotide variants (SNVs) for each species in each sample. In downstream analyses, these calls were used to quantify SNV prevalence across our cohort, to identify QP samples, and to quantify SNV differences between QP samples.

Similar to our pipeline for identifying gene content, MIDAS uses a standard reference based approach to identify SNVs in metagenomic data. Briefly, sequencing reads were first aligned to the host-specific panel of reference genomes using Bowtie2, with default MIDAS mapping thresholds: global alignment, MAPID ≥94.0%, READQ ≥20, ALN_COV ≥0.75, and MAPQ ≥ 20. Species were immediately excluded from further analysis if ≤40% of the genome (the typical core genome fraction) recruited any reads; these excluded cases likely correspond to scenarios where the species is not truly present, but reads from some accessory genes are instead recruited from a different species. Gene annotations for each reference genome were lifted over from the PATRIC database [99], and protein coding sites were classified as 1-fold, 2-fold, 3-fold, or 4-fold degenerate based on the codon reading frame of each annotated gene.

Based on these raw alignments, MIDAS reports the total read coverage *D* for each site in the reference genome for each given sample [100]. We used <u>t</u>his distribution of coverage across the genome to obtain a measure of the “typical” coverage, 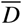, defined as the median of all protein coding sites with nonzero coverage. All samples with 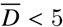 were excluded from further analyses. Additional coverage requirements for QP samples <u>a</u>re imposed below.

We then used the sample-specific estimates of 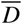 to refine the alignment step above, since 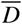 helps to calibrat<u>e</u> our expectation for the coverage at a given site in the genome (*D*). In particular, sites with 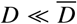 could arise due to mapping errors or read donating from less abundant species, if the refer<u>en</u>ce genome contains regions that are not present in a given sample. Similarly, sites with 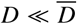 could arise from multi-copy genes, or read donating from m<u>o</u>re abundan<u>t s</u>pecies. To exclude these cases, we masked sites in a given sample if 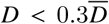 or 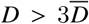 This constitutes a slightly more permissive version of our single-copy criterion above, due to the larger uncertainties inherent in estimating *D*. As above, we only considered sites in coding sequences of annotated genes, and sites that were unmasked in fewer than 4 samples were excluded from all further analyses.

For each of the retained sites, MIDAS reports reference and alternate allele counts using samtools mpileup [100]. (In a minority of cases where multiple alternative alleles were present, these are merged into a single class.) We used these raw allele counts to estimate the prevalence of SNVs in the broader population, defined as the fraction of samples where the alternate allele comprises the majority of reads. Since the reference allele is arbitrarily defined by the choice of reference genome, we used these prevalence estimates to polarize each SNV based on the consensus across the cohort. Polarized within-sample allele frequencies were then defined as the fraction of reads supporting the allele with the lower prevalence across the cohort.

These allele frequencies are used to identify QP samples (SI Section S3.3) and to ultimately quantify SNV differences between QP samples (SI Section S3.4). As with the gene content estimates above, these SNV differences are also susceptible to false positives that occur if the two alleles are actually linked to different species that are simply fluctuating in abundance. We have implemented a number of filters to guard against these events.

First, the global alignment and the MAPQ settings in Bowtie2 already ensure that reads must have a particularly unique match to their assigned reference genome. We only considered sites in protein coding genes, and we excluded all putatively shared genes in the blacklist above. In contrast to previous polymorphism- [31] or consensus-based approaches [35, 44], we considered only extreme changes in allele frequency (≤ 0.2 to ≥ 0.8, or vice versa) to ensure that the SNV difference is supported by the vast majority of the reads at both timepoints, rather than a fraction of reads donated from other species. Combined with the coverage requirement in both samples (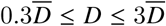 and 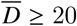), this eliminates most opportunities for SNV differences to arise from abundance fluctuations: large fluctuations will typically violate the coverage requirement, while small fluctuations will not produce a sufficient change in allele frequency.

In cases where we compare longitudinal samples from the same host, we imposed an even stronger version of this filter to be more conservative with respect to calling SNV changes. Under the reasonable assumption that genome synt<u>en</u>y is preserved among very closely related strains, we expect the relative coverage of a site (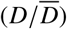 to be more similar in longitudinal samples than the maximum 10-fold range allowed by the coverage condition 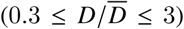. Thus, in addition to the requirements above, we only called a SNV difference between two samples if the successive values of 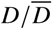 were within a factor of 3.

#### S1.5 Identifying orthologous genes in different pangenomes

The species-specific pangenomes in the MIDAS database were constructed by clustering all genes found in the isolate genomes of each species using a 95% identity threshold [33]. However, this clustering approach leaves open the possibility that a gene in one species’ pangenome may have sequence similarity ≥ 95% to a gene in another species’ pangenome. We identified these cross-pangenome orthologs as follows.

First, for computational efficiency, we focused on human-relevant bacterial species in the MIDAS database by identifying those isolates with the keywords ‘human’ or ‘Homo sapiens’ in the host column of the PATRIC database. We also included species that had a universal single copy gene marker coverage ≥1x in at least one sample in our cohort. This resulted in 1002 human-relevant species.

Next, we ran USEARCH [101] on the set of genes belonging to the pangenomes of these human-relevant bacterial species. Based on this approach, we identified a total of 890,058 genes across these 1002 species that had ≥95% sequence identity with at least one other gene in a different species’ pangenome. These genes were excluded from further analysis as described above.

## S2 Text

### Quantifying within-species diversity in individual samples

#### Estimating rates of core-genome polymorphism

To estimate the overall levels of nucleotide polymorphism for a given species in a given sample (Fig. 1E), we calculate the fraction of synonymous sites in core genes with intermediate allele frequencies (0.2 ≤ *f* ≤ 0.8). In other words, the polymorphism rate *r* is defined by

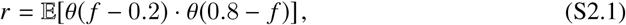

where θ(*z*) is the Heaviside step function. This measure is similar to the traditional population genetic measure of heterozygosity, *H* = 𝔼[2*f*(1 – *f*)], which places the most weight near intermediate allele frequencies. The thresholded version in Eq. (S2.1) is preferable in our case, as it is more robust to low-frequency sequencing errors that can overwhelm the average in *H*.

To obtain the approximate confidence intervals for the rates in Fig. 1E, we used a standard Bayesian procedure based on a poisson approximation. If we let *L* denote the total number of sites examined and let *n* denote the number of “successes” (i.e., the number of intermediate frequency polymorphisms), then we assume that *n* is drawn from a Poisson

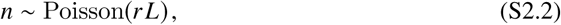

where *r* is the per site rate plotted in Figs. 1E. Since *r* is a positive quantity that varies over many orders of magnitude, we use a uniform prior over log *r*. After applying Bayes’ rule, this yields a standard conjugate Gamma posterior distribution for *r*:

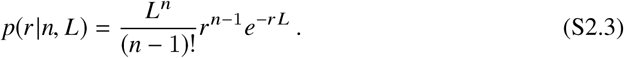

whose posterior mean is just

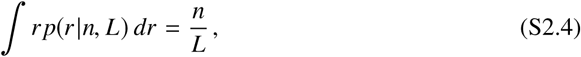

as expected. For all *n* > 0, we define a 1 – α confidence interval to be the α/2 and 1 α/2 percentiles of this posterior distribution. In the case where *n* = 0, the posterior distribution is improper:

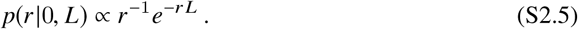

In this case, we define the lower limit of the confidence interval to be 0, and the upper limit to be the point where *e*^-*r L*^ ∼ α/2.

## S2. Within-host evolution in a single-colonization model

In this section, we further explain the assumptions made in computing the expected within-host polymorphism rate for a given species under a simple, single-colonization model. As described in the text, we make conservatively high estimates for the per site mutation rate (µ ∼10^-9^ per generation), generation times (λ ∼10 generations per day), and time since colonization (Δ*t* ∼100 years). We define the within-host polymorphism rate *P* as the fraction of fourfold-degenerate synonymous site mutations with allele frequencies in the range 0.2 ≤*f* ≤0.8. In the single-colonization model, the mutations that contribute to *P* must have reached intermediate frequencies after starting as a *de novo* mutation at some time after colonization.

We assume that the synonymous mutations are effectively neutral over the timespans considered (*s*λΔ*t* ≪ 1). Under this assumption, one of these mutations can only contribute to *P* if it hitchhiked along with a lineage that rose to a frequency in the range 0.2 ≤*f* ≤ 0.8. This can happen either due to neutral drift (i.e., the lineage randomly fluctuated to intermediate frequencies) or selection (i.e., the lineage reached intermediate frequencies because it contains a beneficial mutation). However, if synonymous mutations are neutral, their presence or absence in a lineage is independent of the processes that drive it to intermediate frequency [102]. The probability that a particular neutral mutation arose along the line of descent is simply the product of the per-site mutation rate µ and the total number of generations since the lineage diverged from the common ancestor between it and the rest of the population. By assumption, the latter is bounded by the total number of generations since colonization (λΔ*t*). This yields the conservative estimate for the within-host polymorphism rate,

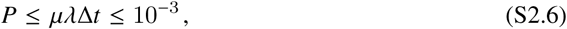

quoted in the main text.

### S3 Text

#### Quasi-phasing metagenomic samples

In this section, we describe the methods used to estimate one of the dominant haplotypes for a given species in a subset of metagenomic samples (the so-called *quasi-phaseable* or QP samples), and to quantify genetic differences between these lineages. The method is similar in spirit to recent work by Ref. [35], but with a greater emphasis on estimating the associated false positive rates.

##### S3.1 Theoretical motivation

To gain intuition for how within-host lineage structure is reflected in the distribution of allele frequencies, it is useful to start by considering the simplest version of the phasing problem, in which the metagenomic reads for a given species in a particular sample are derived one of two clonal lineages mixed in a proportion *f*_mix_ ≥50% (representing the proportion of cells from the more abundant lineage). Within-sample polymorphisms will arise from fixed differences between the two lineages and will segregate at frequency *f*_mix_ or 1 – *f*_mix_, depending on which lineage the mutation arose in and the choice of reference allele. Since this choice is arbitrary, we work with the major allele frequency in each sample. In this case, the distribution of major allele frequencies, *p*(*f*), will then have the simple form

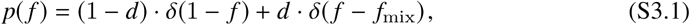

where *d* is the average nucleotide divergence between the two lineages and δ(*z*) is the Dirac delta function. Note that this theoretical distribution is only obtained in the limit of infinite coverage; in practice, the observed distribution of major allele frequencies will be blurred due to sampling noise (see Text S3.2 below). Nevertheless, in the the limit of high coverage, Eq. (S3.1) suggests that we can infer *f*_mix_ and *d* by looking for a peak in the distribution of major allele frequencies (e.g., Fig. 1E). Again, in the idealized case, the two haplotype sequences are easy to recognize: major alleles are assigned to the dominant lineage, while the minor alleles belong to the subdominant type.

This basic idea also extends to mixtures of more than two lineages, but the potential genealogical relationships between them make the problem much more complicated. For example, in a mixture of three strains with frequencies *f*_1_, *f*_2_, and *f*_3_, the distribution of major allele frequencies will now have three characteristic peaks (corresponding to min {*f*_*i*_, 1– *f*_*i*_} for each *i* = 1, 2, 3). This time, however, alleles that segregate at the same frequency do not necessarily belong to the same lineage, since they could also be ancestral to two of the three strains. There are three possible genealogies relating the three strains, which can vary from site-to-site in the presence of recombination. Haplotype estimation then becomes a complicated inference problem, which only grows more difficult as additional lineages are added. Consideration of the combined allele frequency distribution may be helpful for deriving error models for algorithms that attempt to deconvolute strains from metagenomes.

Rather than trying to infer the exact mixture proportions and the haplotypes of each lineage, we developed a set of heuristic rules to identify the haplotype of just *one* of the dominant lineages while controlling the probability of misassigning variants to this haplotype. Suppose that there are within-sample polymorphisms at two sites, with major allele frequencies *f*_1_ and *f*_2_. We denote the four (unobserved) two-locus haplotype frequencies by *f*_*M* *M*_, *f*_*Mm*_, *f*_*mM*_, and *f*_*mm*_, where *M* and *m* denote the major and minor allele at each site. If *f*_1_ = *f*_2_ = 0.5, then there are no constraints on the possible haplotype frequencies, other than the marginal constraints *f*_*M*__*M*_ + *f*_*Mm*_ = *f*_1_ and *f*_*M*__*M*_ + *f*_*mM*_ = *f*_2_. However, in the opposite extreme where *f*_1_ = *f*_2_ = 1, then normalization constraints require that *f*_*M*__*M*_ = 1 (i.e., the major alleles are on the same haplotype). In between these two extremes there is a more general rule that, whenever the allele frequencies satisfy *f*_*i*_ ≥*f*, with log (*f*/1–*f*) = *c* ≳ 1, the minimum possible frequency of the *MM* haplotype is

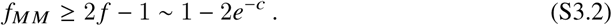

Equation S3.2 represents a worst case scenario in which the haplotypes are specifically assigned to prevent major alleles from segregating together. In practice, a more realistic lower bound for the *f*_*M* *M*_ is attained when the alleles are in linkage equibrium:

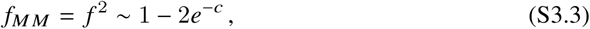

which happens to have the same asymptotic behavior in this two-locus example. In either case, these bounds show that an appreciable fraction of cells in the host must possess both major alleles. This argument can also be extended to larger collections of sites. In the pessimistic case of linkage equilibrium between all polymorphic sites, the number of major alleles per individual is binomially distributed with success probability *f*. In the limit of a large number of sites, this means that the vast majority of the cells will have the major allele at a fraction *f* of the possible sites. However, while the haplotype consisting of all major alleles is the most likely haplotype under linkage equilibrium, its expected frequency can grow quite small, to the point where the haplotype may not even be present in a finite sample. Fortunately, our analysis will primarily focus on one- and two-locus statistics where the stronger bounds in Eq. (S3.2) can be applied.

##### S3.2 False positive rate for SNV phasing

The arguments above suggest that, for many downstream purposes, we can effectively estimate a portion of one of the haplotypes in a metegenomic sample by taking the major alleles present above some threshold freuqency, *f* ^*^≫ 50%, and treating sites with intermediate frequencies as missing data. This is a simple generalization of the consensus method (i.e. taking the haplotype formed by all major alleles) that has been used in previous metagenomic studies [4, 35], and it is similar to methods used to genotype clonal isolates from whole-genome resequencing data [103].

The major difficulty with this approach is that we do not observe the true frequency *f* directly, but rather a sample frequency 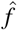 that is estimated from a finite number of sequencing reads. Polarization errors (i.e. errors in determining the major allele) can therefore accumulate when the allele supported by the most reads differs from true major allele in the sample. When sequencing clonal isolates, such false positives are primarily caused by sequencing errors. These occur at a low rate per read (*p*_err_ ∼1% per bp), and become increasingly unlikely at moderate sequencing depths. However, in a metagenomic sample, polarization errors will also arise due to finite sampling noise, when an allele at some intermediate frequency (e.g. 25%) happens to be sampled in a majority of the sequencing reads. As we will show below, for moderate sequencing depths, this will often be the dominant source of error.

To model this process, let (*A*_*𝓁*_, *D*_*𝓁*_) denote the number of alternate alleles and total sequencing depth at a given site t in the genome, and let 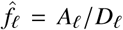 denote the corresponding sample frequency. We assume that the number of alternate reads follows a binomial distribution,

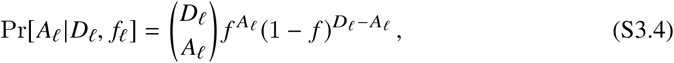

for some true frequency *f*_*𝓁*_, so that the probability of observing 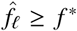 is simply

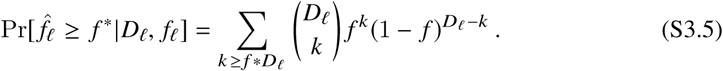

A polarization error will occur when we observe 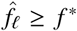 even though *f*_*𝓁*_ < 50%. Equation (S3.5) shows the probability of such an error will strongly depend on *f*_*𝓁*_. For a sequencing depth of *D* = 10 and a frequency threshold of *f* ^*^ = 80%, the error probability ranges from essentially negligible (∼10^-14^) when *f* is on the order of the sequencing error rate (∼1%), to ∼1 per bacterial genome when *f* ≈ 10%, to an error rate of 5% when *f* ≈ 50%.

The average false positive rate across the genome will therefore depend on an average over the possible values of *f* and *D*:

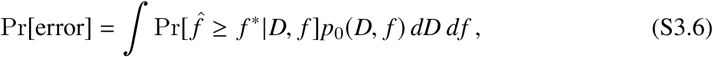

where *p*_0_(*D, f*) is the prior distribution of *D* and *f* at a randomly chosen site (Fig. S1A). In the absence of any additional information, this joint distribution with the product of empirical distributions,

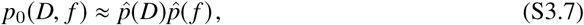

which we estimate for a given sample by binning the observed values of *D* and the allele frequencies across the *L* sites under consideration (blue distribution in Fig. S1A). The expected number of polarization errors in a given sample across all *L* sites is given by

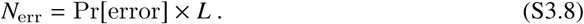

This calculation holds for any large collection of sites where the empirical distribution, 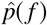, provides a reasonable approximation to the prior distribution, *p*_0_(*f*). For example, in the following section, we consider the set of all synonymous sites in the core genome.

##### S3.3 Quasi-phaseable (QP) samples

The basic idea behind our approach is that we wish to restrict our attention to samples where *N*_err_ is small compared to the total number of sites under consideration. This number will vary depending on the particular analysis that we wish to carry out. But for population-genetic purposes, it will always be related to the number of sites that actually vary between samples. As a simple proxy for this number, we therefore consider a measure of the average genetic distance between the dominant haplotype in a given sample and the lineages in the remainder of our panel.

Specifically, we focus on fourfold-degenerate synonymous sites in the core genome. For each sample, let *N*_<_ denote the number of such sites with major allele frequencies less than *f* ^*^, and conversely, let *N*_>_ denote the number of sites with 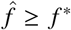. For the sites in the latter group, let 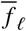 denote the corresponding allele frequency across the entire panel. Then the quantity

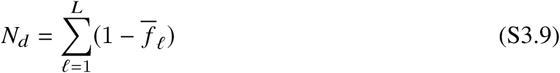

approximates the expected number of differences at these sites for an “average” individual drawn from the panel. A normalized version (*N*_*d*_/*L*) is illustrated for the *B. vulgatus* samples in Fig. S2. We declare the sample to be a **quasi-phaseable (QP)** sample if it passes the coverage thresholds in SI Section S1 Text and *N*_<_/*N*_*d*_ < 0.1.

To see why this is a reasonable definition, we return to our error formula in Eq. (S3.8) and plug in conservative estimates for *p*_0_ (*D, f*). For example, we expect that the number of truly polymorphic sites in the sample will also be of order ∼ *N*_*d*_, with the remaining sites having frequencies near the sequencing error threshold, *f* ∼1%. We then divide the remaining polymorphic sites into the fraction *N*_<_/*N*_*d*_ ≲ 0.1 with major allele frequencies below *f* ^*^, and the remaining fraction (∼100%) with major allele frequencies above *f*^*^. If we make the conservative approximation that all of the sites in the latter group have minor allele frequencies *f* ≈ 1 – *f* ^*^, and all of the sites in the former group have *f* ≈ 50%, then we obtain an approximate prior distribution for *f* :

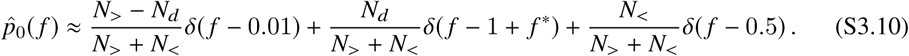

If we make a similarly conservative approximation for the coverage distribution,

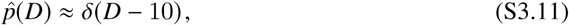

where δ is the Dirac function, then for a threshold of *f* ^*^ = 80%, the realized false positive rate is

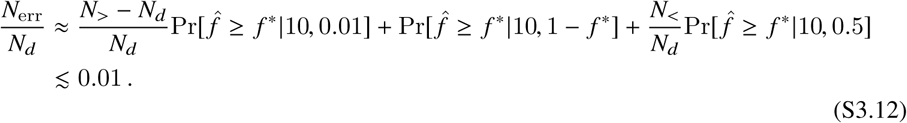

Thus, with these thresholds, we expect that only a small fraction of informative sites (as defined by the average distance between samples) will be susceptible to polarization errors.

##### S3.4 False positive rate for SNV differences

Although the QP sample classification is a good rule of thumb for determining when polarization errors are more or less likely to happen, there are scenarios where we wish to measure genetic distances between samples (e.g. longitudinal samples from the same individual) that are much more closely related than an average pair of individuals in our panel. In these cases, the realized false positive rate can be much higher than the estimate in Eq. (S3.12). To obtain more accurate estimates of the error in these cases, we extend our calculation above to the specific problem of detecting the number of nucleotide differences between two samples.

Generalizing from the phasing problem above, we would conclude that the haplotypes in two samples share the same allele at a given site if that allele is present above frequency *f* ^*^ in both samples. To observe a difference between the two samples, the allele would have to be present above frequency *f* ^*^ in one sample and below 1 – *f* ^*^ in another. If the allele lies between 1 – *f* ^*^ and *f* ^*^ in one of the samples, the site is treated as censored data. Under this definition, a nucleotide difference requires a change in allele frequency of at least

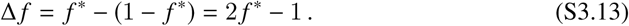

If we rewrite everything in terms of Δ *f*, a nucleotide difference requires the allele frequency to lie below (1 – Δ*f*)/2 in one sample and above (1 + Δ*f*)/2 in another (pink shaded regions in Fig. S1B). We will adopt the latter notation here, as it allows us to easily consider more stringent thresholds for which Δ*f* > 2*f*^*^-1.

Under the null hypothesis, we assume that the true allele frequency *f* is the same in the two samples. If we let *D*_1_ and *D*_2_ denote the coverage of the site in the two samples, then a simple generalization of Eq. (S3.6) shows that the false positive rate for a randomly chosen site is given by

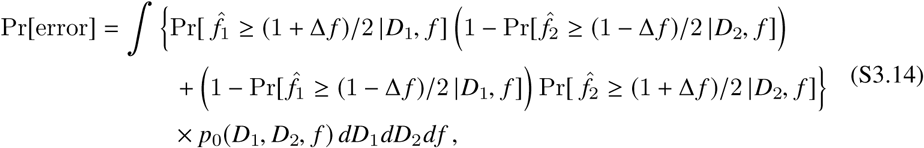

where 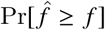 is defined in Eq. (S3.5) and *p*_0_ (*D*_1_, *D*_2_, *f*) is the prior distribution for *D*_1_, *D*_2_, and *f* at a random site. As in Eq. (S3.7) above, we estimate this prior distribution as a product of empirical distributions,

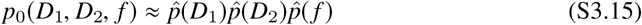

which we estimate by binning the observed values of *D*_1_, *D*_2_, and 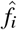 cross the genomes of the two samples (the blue distribution in S1B). The expected number of false positive substitutions is then given by

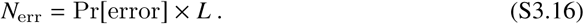

where *L* is the total number of sites compared between the two samples. This will vary depending on the application (e.g. synonymous sites, sites in core genes, all coding sites, etc. are used at various times in the main text).

The error estimate in Eq. (S3.16) is an implicit function of the threshold Δ*f*. Given the typical sequencing coverage and allele frequency distributions of the QP samples in our analyses, we usually obtain sufficiently low error estimates (i.e., *N*_err_ ≪ 1) if we take Δ *f* = 1 – 2*f*^*^ = 0.6, so that an allele transitions from less than 20% to greater than 80% frequency between the two samples, or vice versa. To limit the influence of outliers, we excluded all pairs of samples with *N*_err_ > max {0.5, 0.1*N*_obs_}, where *N*_obs_ is the observed number of SNV differences.

##### S3.5 False positive rate for gene content differences

The false positive rate for gene content differences can be estimated with a similar procedure. In this case, the canonical generative model is one in which a gene *g* with average copy number per cell *c*_*g,i*_ in sample *i* recruits *N*_*g,i*_ reads, which we assume follows a Poisson distribution:

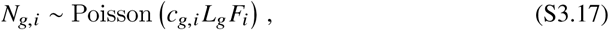

where *L*_*g*_ is the length of gene *g* and *F*_*i*_ is a sample- and species-specific constant that reflects the total number of reads aligned to that species (e.g., by the MIDAS pipeline). The coverage of gene *g* is then defined as

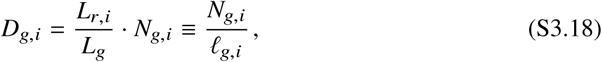

where *L*_*r,i*_ is the average length of reads that align to that gene (typically ≲ 100bp), which can vary in a sample-specific manner. The quantity *𝓁*_*g,i*_ ≡ *L*_*g*_/*L*_*r,i*_ then serves as a conversion factor between the raw number of reads and the coverage. Finally, we assume (as in the MIDAS pipeline) that there is a known panel of marker genes (*g* = *m*) with fixed copy number per cell of *c*_*m*_ ≈ 1 and a large target size, such that *N*_*m,i*_ ≈ 𝔼[*N*_*m,i*_] = *L*_*m*_*F*_*i*_. This allows us to eliminate *F*_*i*_ and rewrite Eq. (S3.17) in terms of the marker coverage *D*_*m,i*_ and the coverage-to-read conversion factor *𝓁*_*g,i*_:

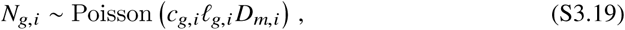

The variables *N*_*g,i*_, *D*_*g,i*_, and *D*_*m,i*_ are all reported by MIDAS, which allowed us to estimate *c*_*g,i*_ and *𝓁*_*g,i*_ for each gene in each sample:

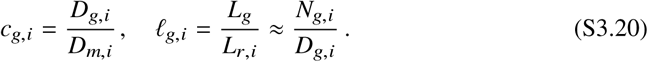

Based on the above error rate calculations, the gene copy number change events we are interested in are those in which a gene transitions from a typical single-copy value (0.6 ≤ *c* ≤ 1.2, see Fig. S6) in one sample to a value close to zero (*c* < 0.05) in another. This does not cover all possible copy number change events, but focuses on the subset that are likely to be (i) statistically significant and (ii) less susceptible to other bioinformatic errors (e.g. read stealing or donating from other species), see Text S1.3.

Given this definition, the probability of an apparent copy number change happening by chance will again depend on the “true” copy number of the gene, *c*, as well as its effective coverage, *𝓁D*. Similar to Eq. (S3.14), the expected false positive rate for a randomly chosen gene is given by

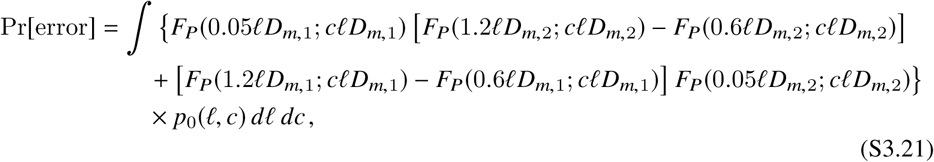

where *F*_*P*_ (*k*; λ) is the Poisson CDF and *p*_0_ (*𝓁, c*) is the null distribution of *𝓁* and *c*. Once again, we estimate this joint distribution with the product of empirical distributions,

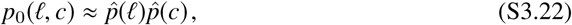

which are estimated by binning the observed values of *𝓁*_*g,i*_ and *c*_*g,i*_ across the two samples. To reduce mapping artifacts, we only bin t-values from genes with copy number in the range 0.6 ≤ *c* ≤ 1.2, which accounts for the bulk of the copy number distribution in a given sample (S6). The expected number of false positive gene changes is therefore given by

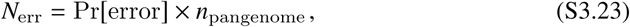

where *n*_pangenome_ is the total number of genes tested (typically of order ∼10^4^). For the typical coverages in our dataset, this number is usually very small (≪ 10^-2^). As above, we excluded all pairs of samples where *N*_err_ ≤ max {0.5, 0.1*N*_obs_}, where *N*_obs_ is the observed number of gene content differences differences between those samples.

##### S3.6 Validation with synthetic data

As a sanity check on our calculations above, we validated our method using synthetic metagenomic data generated by Grinder [104]. To simulate the null hypothesis, we generated synthetic sequencing reads from two *Bacteroides vulgatus* isolates mixed at a 9:1 ratio at both timepoints. We performed these simulations for target coverages of 20x, 50x, and 100x. Two replicate simulations were performed for each coverage value for two difference combinations of isolate genomes, resulting in 4 independent experiments per coverage group. After running these synthetic metagenomic samples through the steps of our pipeline, we found zero SNV or gene changes between the two timepoints for all 12 experiments across the coverage values. This provides further support for the claim that the false positive rate from sampling error is; ≲ 0.1 per genome, and it suggests that this claim is robust to additional noise introduced during the mapping and thresholding steps in SI Section S1 Text.

### S4 Text

#### Population genetic null model of purifying selection for pair-wise divergence across hosts

In this section, we present a minimal model of purifying selection that can account for the varying *d*_*N*_/*d*_*S*_ levels in Fig. 2D as a function of *d*_*S*_. The basic idea is that purifying selection is less efficient at purging deleterious mutations that are very young (in particular, younger than the inverse of the associated fitness cost). To the extent that synonymous divergence can be associated with a characteristic timescale, this line of reasoning implies that anomalously low values of *d*_*S*_ would be associated with less efficient purifying selection (i.e., higher values of *d*_*N*_/*d*_*S*_), while typical values of *d*_*S*_ would be associated with more efficient purifying selection (i.e., lower values of *d*_*N*_/*d*_*S*_). Similar ideas have been employed in previous studies [105, 106].

To make this idea more concrete, suppose that the age of a given mutation is bounded by a time *T*, so that it occured at some point in the last *T* generations. This will result in a genetic difference between two randomly sampled lineages with probability

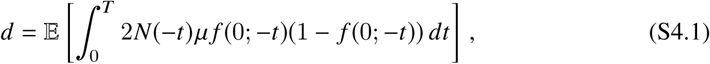

where *N* (*t*) is the effective size of the across-host population, and *f* (*t*; *t*_0_) is the prevalence of an allele that was created at time *t*_0_ and sampled at time *t*, and the expectation is taken over all possible realizations of *f* (*t, t*_0_). If *T* is much smaller than the typical coalescence timescale across hosts, then the mutation cannot rise to a very high prevalence by the time of sampling, and we can neglect the *f* ^2^ term above to obtain

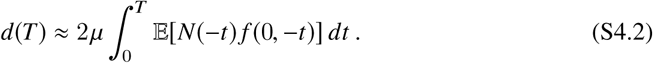

By definition, the new mutation will enter at prevalence 1/*N* (– *t*). If the mutation has a deleterious fitness cost *s*, then its average size is simply

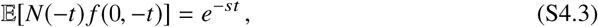

and we have

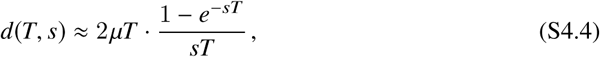

If synonymous mutations are assumed to be neutral, then

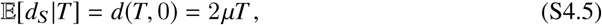

as expected. If we assume that the nonsynonymous sites have a distribution of deleterious fitness costs ρ(*s*), then the nonsynonymous divergence rate satisfies

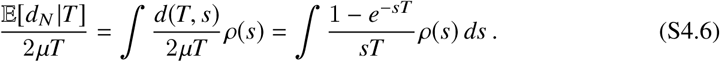

In the simplest case, ρ (*s)* will contain a mixture of truly neutral mutations and a fraction *f*_*d*_ with deleterious fitness cost *s*, for which

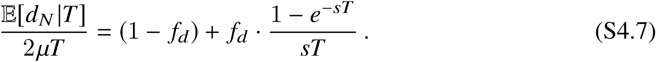

To connect this model with the observed data, we must find a way to estimate *T*. Motivated by the fact that 𝔼[*d*_*S*_ |*T*] = 2µ*T*, we assume that for anomalously low core-genome-wide divergence rates (*T* ≪ *T*_*c*_), the method-of-moments estimator 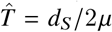 provides a reasonable estimate of the maximum mutation age *T* at most polymorphic loci (otherwise, we would expect a more typical value of *d*_*S*_). However, a complicating factor is that *T* is present on both sides of Eq. (S4.7). Using the same estimator for the *x* and *y* axes in Fig. 3 can lead to spurious correlations that arise from measurement noise, which mimic the true biological signal in Eq. (S4.7). To avoid this issue, we partition the synonymous sites into two artificial categories, which produces two divergence estimates *d*_*S*,1_ and *d*_*S*,2_. By the Poisson thinning property, these are conditionally independent given *T*. Thus, we can use one value of *d*_*S*_ to estimate *T* on the left-hand side of Eq. (S4.7) and one value of *d*_*S*_ to estimate *T* on the right-hand side of Eq. (S4.7), yielding the empirical relation between *d*_*N*_, *d*_*S*,1_, and *d*_*S*,2_,

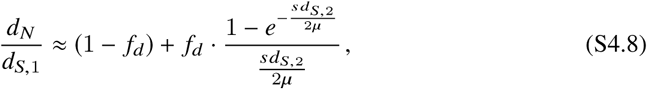

which should be valid for *d*_*S*_ much smaller than the population median. For small *d*_*S*_, this ratio will start to deviate from unity when *d*_*S*_ *≳* 4µ/*s f*. At large *d*_*S*_, the ratio approaches 1 – *f*_*d*_, and will start to deviate from this value when *d*_*S*_ ≲ 2µ *f*_*d*_/*s*(1 – *f*_*d*_). These landmarks allow us to obtain approximate estimates of *f*_*d*_ and *s* by rough inspection of the data in Fig. 3. To obtain the confidence intervals the inset of Fig. 3, we generated bootstrapped datasets by Poisson resampling the synonymous and nonsynonymous counts between each pair of lineages, and applying the same thinning procedure as above.

We note that qualitatively similar behavior is expected in recent models of bacterial evolution proposed by Ref. [75], in which the core genome of closely related strains consists of an asexual “backbone” or “clonal frame” (where synonymous mutations occur at rate µ) interrupted by highly diverged segments of length *𝓁*_*r*_ acquired through recombination. The introgressed segments would enter with low values of *d*_*N*_/*d*_*S*_ associated with the average *d*_*S*_ value. If the common ancestor of the asexual backbone is younger than the typical deleterious fitness cost, we would again expect a transition from essentially neutral behavior (*d*_*N*_/*d*_*S*_ ≈ 1) to the typical between-host value (*d*_*N*_/*d*_*S*_ ≈ 0.1) as a function of *d*_*S*_, where the transition is now informative of the horizontal transfer rate. A formal analysis of this model remains an interesting avenue for future work.

### S5 Text

#### Phylogenetic inconsistency and clade structure across hosts

#### S5.1 Phylogenetic inconsistency

In this section, we describe the methods used to assess phylogenetic inconsistency in Fig. 4A. Traditionally, phylogenetic consistency is measured by first obtaining a genome-wide estimate of the genealogical relationships between lineages, and then asking whether individual SNVs can be explained by a single mutation event on this fixed tree [107, 108]. SNVs that cannot be explained this way are said to be *homoplasic* or *phylogenetically inconsistent*.

The major drawback with this approach is that it requires an accurate estimate of the genome-wide phylogeny. Statistical uncertainties or model misspecification in the genealogical inference step can lead to inflated estimates of inconsistency. More importantly, in cases where significant portions of the genome are phylogenetically inconsistent, it is also difficult to pinpoint the source of the inconsistencies, since they can bias the genome-wide phylogeny in unknown ways. To avoid these issues, we developed a non-parametric approach for quantifying the phylogenetic inconsistency of SNVs directly from the core-genome-wide divergence values in Fig. 2, which eliminates the need to first infer a genome-wide tree.

The idea behind our method is simple. In an infinite sites model, partial information about the genealogy of an individual SNV is encoded in the allelic states of different individuals. In particular, all of the individuals with the derived allele must be more closely related to each other than to individuals with the ancestral allele. Under asexual evolution, the distribution of coalescence times between pairs of individuals (*t*_*ij*_) also encodes information about the genealogy at the SNV site. In particular, the descendents of a coalescent event must have smaller values of *t*_*ij*_ among themselves than they do with individuals in other parts of the tree.

To connect these two pieces of information, we note that all individuals that share a mutation by descent must have coalesced more recently than the age of the mutation. Similarly, individuals with different allelic states must have coalesced further back in time than the age of the mutation (otherwise they would share the mutation by descent). This also implies that the minimum *t*_*ij*_ for individuals with different allelic states must be an upper bound on the age of the mutation, and conversely, the maximum *t*_*ij*_ between derived individuals is a lower bound. If this lower bound exceeds the upper bound, then the SNV is phylogenetically inconsistent (Fig. S10).

To connect this mathematical intuition with the data, we note that the coalescence time is related to the total divergence through the method-of-moments estimator,

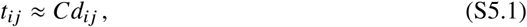

for some species-dependent clock constant *C*. If we let *M* denote the set of individuals with the major allele, and *m* denote the set of individuals with the minor allele, we can then define a critical divergence

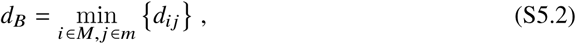

which can be used to infer the upper bound on the age of the mutation:

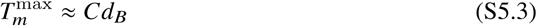

Similarly, we can define a second set of critical divergences for each allele,

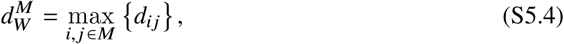

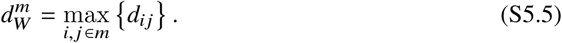

If we knew which allele was the ancestral one, and which was the derived, we could use the corresponding value of *d*_*W*_ to estimate the lower bound on the age of the mutation. Since we do not have this information, we have to take the minimum of these two values,

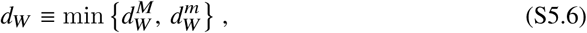

so that

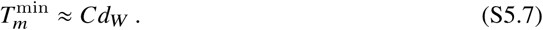

If the ratio between *d*_*W*_ and *d*_*B*_,

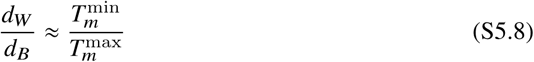

is substantially greater than one, then there is evidence that the SNV is phylogenetically inconsistent.

To implement this logic in Fig. 4A, we chose a threshold divergence 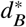 and looked for all SNVs that occured more recently than this (i.e., those with 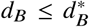) and which had at least two minor alleles (so that *d*_*W*_ is well-defined). We defined the net amount of phylogenetic inconsistency at 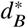 to be the fraction of SNVs in this set with 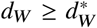, for some threshold 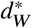 To be conservative, we chose

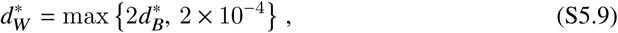

which ensures that all inconsistent SNVs have *d*_*W*_ ≥ 2*d*_*B*_. The factor of 2 was chosen to match traditional notions of sequence similarity clusters (or “ecotypes”) [62].

#### S5. Clustering and identification of top-level clades

In some species, we observed very high levels of phylogenetic consistency for SNVs that separate the most distantly related strains, and a sudden transition to high levels of inconsistency for intermediate levels of divergence. In these species, there is often a second mode in the distribution of core-genome-divergence at the high end of the spectrum. This suggests that the lineages may represent a mixture of two genetically isolated populations, e.g. different subspecies or ecotypes. Given the purely operational species definition used by MIDAS (95% ANI), it is not surprising that genetically isolated populations can sometimes fall below this species threshold and their metagenomic reads can map to the same reference genome.

Mixtures of genetically isolated populations can confound traditional SNV-based estimates of recombination within species, since more SNVs will have accumulated between genetically isolated populations than within them. To account for these biases, we manually partitioned each species into a few “top-level” clades, which we hypothesized could better approximate a genetically cohesive population. Note that this partitioning scheme is conservative for detecting recombination: subsetting individuals cannot create evidence for recombination where there is none, but the lack of evidence for recombination could simply indicate that we chose the clades poorly.

Our approach for identifying clades is based on traditional notions of sequence similarity clusters [62, 79]. We first constructed core-genome dendrograms by hierarchically clustering the matrix of pairwise divergence rates averaged across the core genome, using the UPGMA method from SciPy [109]. Based on these dendrograms, lineages were assigned to one or more “top-level” clades using a manual procedure, loosely designed to maximize the difference between inter- and intra-clade divergence at the most deeply diverged branches (Table S2). We adopted this manual procedure to capture clade structure that is inconsistent with a single cut through the dendrogram at a given level of divergence.

In Fig. S11, we plot the fixation index, *F*_*st*_ for these manually defined clades:

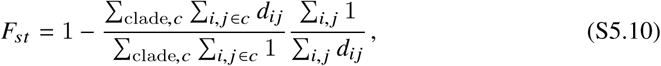

where *c* indexes the clades and *d*_*ij*_ is the average nucleotide divergence across core genes in hosts *i* and *j*. Several of the prevalent species have top-level clades with high *F*_*st*_. *B. vulgatus* serves as one of the more extreme cases, owing to the fact that the *B. vulgatus* and *B. dorei* clades are both clustered to the *B. vulgatus* reference genome. However, this is not a universal pattern across gut bacteria: some species, even other *Bacteroides* like *Bacteroides xylanisolvens*, have lineage phylogenies and recombination patterns that are more consistent with a single clade (Fig. 4C).

### S6 Text

#### Population genetic null model for the decay of linkage disequilibrium

In principle, the rate of decay of linkage disequilibrium in Fig. 4 contains information about the average recombination rate between pairs of loci [16]. For example, in a neutral panmictic population of size *N*, Ref. [60] have shown that

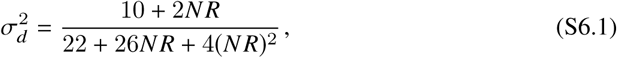

where *R* is the recombination rate between two loci. Similar functional forms are expected for related measures of linkage disequilibrium (e.g. *r*^2^ [110]). To obtain a relation between the recombination rate *R* and the genomic distance *𝓁* between two loci, we assume that recombination occurs through the exchange of DNA fragments of with average length *𝓁* _*r*_, which are exponentially distributed around this mean value and occur uniformly across the genome. Two loci undergo a recombination event when there is a genetic exchange that involves only one of the two loci. This happens with probability

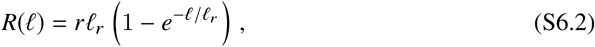

where *r* is a rate constant. Thus, for distances much shorter than *𝓁*_*r*_, this recombination model resembles a linear chromosome with a crossover rate *r* per site. For larger distances, Eq. (S6.2) shows that the effective recombination rate saturates at *r𝓁*_*r*_. Substituting *R* (*𝓁*) into Eq. (S6.1), the decay of linkage disequilibrium will have the characteristic shape

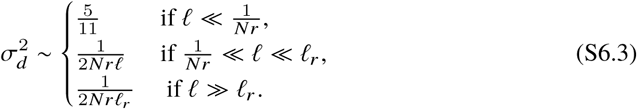

To estimate 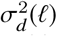 for a given species, we focused on lineages from the largest top-level clade defined in Table S2. Since Fig. 2D suggests that evolutionary forces may be different for closely related strains, we chose only a single lineage from each subclade defined by cutting the core genome tree at divergence *d* = 10^-3^. For pairs of SNVs in the same gene, we assigned a coordinate distance *𝓁* based on their relative position on the reference genome. For a given value of t, we then estimated 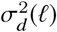 via

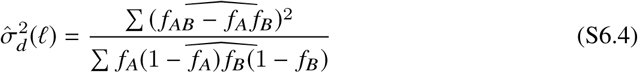

where the sum runs over all pairs of synonymous sites with distances within the range (*𝓁* – Δ*𝓁, 𝓁*+Δ*𝓁*), as described in Fig. 4. Here, *f*_*A*_ = *f*_*Ab*_ + *f*_*AB*_, and *f*_*B*_ = *f*_*aB*_ + *f*_*AB*_, where *f*_*AB*_, *f*_*Ab*_, and *f*_*aB*_ denote the frequencies of the gametic combinations in the across-host population. The hat symbols denote unbiased esimators for the respective quantities underneath, based on the observed gamete counts *n*_*AB*_, *n*_*Ab*_, *n*_*aB*_, and *n*_*ab*_ in our sample of hosts. We assume that the counts are sampled from the frequencies through the multinomial distribution,

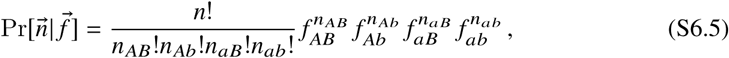

where *n* = *n*_*AB*_ + *n*_*Ab*_ + *n*_*aB*_ + *n*_*ab*_ is the total sample size. The estimate for the hat symbols above are constructed via linear combinations of polynomials in the *n*’s chosen to have the same expected value as the quantity underneath the hat. These expressions are somewhat unwieldy, but are provided in the associated computer code.

After applying this method, we obtain estimates of within-gene σ^2^ (*𝓁*) as a function of *𝓁*, and a core-genome-wide value estimated from SNVs in different genes (Fig. 4), which can be compared with the theoretical prediction in Eq. (S6.3). Because the core-genome-wide value of 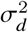 is usually much lower than its intragenic counterpart, we assume that *𝓁*_*r*_ is much larger than the ∼3000bp intragenic window we consider, so we formally set *𝓁*_*r*_ = ∞. However, it is also clear from Fig. 4 that 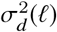 does not always approach the neutral expectation as *𝓁* → 0. As is common practice, we therefore consider an expanded class of models of the form

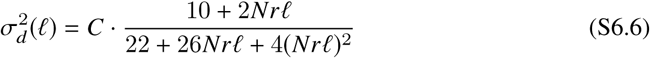

for some arbitrary normalization constant *C*, which must be jointly estimated from the data. (The introduction of *C* is equivalent to focusing on the percentage change in 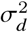, rather than its absolute value.)

This model has two free parameters (*Nr* and *C*), which can be estimated from the observed values of 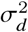 at any two values of *𝓁*. We fix one of these at a reference location *𝓁*_1_ = 9bp, which was chosen to balance the desire to have *𝓁*_1_ ≪ 1/*Nr*, but also to be as large as possible to minimize contamination from compound mutation events. For the second value of 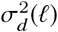, we focus on distances of the form

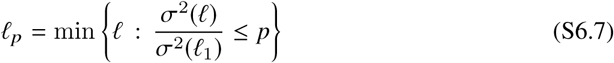

for some fraction *p* (e.g., *p* = 1/2, *p* = 1/4, etc.). In other words, *𝓁* _*p*_ is the distance at which the observed value of σ^2^ (*𝓁*) first falls to a percentage *p* of its value at *𝓁* _1_. According to the model in Eq. S6.6, these distances should satisfy

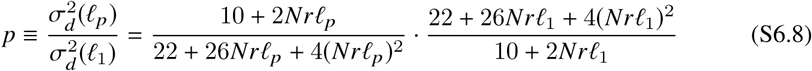

which depends only on *Nr*, in addition to the observed values of *p, 𝓁* _1_, and *𝓁* _*p*_. Solving this function numerically, we obtain estimates for *Nr* for different values of *p*.

In the neutral model that leads to Eq. S6.1, the population size *N* can be estimated from the average pairwise divergence, *d*_*S*_ = 2*N* µ. Thus, we normalize the estimated values of *Nr* by *d*_*S*_/2 to obtain an estimate of the ratio *r*/µ for different values of *p*. As long as the model is a good description of the data, these estimates should be approximately independent of the choice of *p*. The observed deviations in *r*/µ as a function of *p* (Fig. S14) point to fundamental deviations from the model in Eq. (S6.6) that cannot be accounted for by simply varying the parameters. This suggests that the decay of 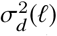 may hold power for investigating departures from the simple neutral model above (e.g. to include hitchhiking, population structure, variation in recombination rate within genes, etc.).

### S7 Text

#### Validation of between-host patterns using isolate sequences

A major practical advantage of our metagenomic approach is that it can resolve a large number of quasi-phased genomes across many species, using data from a much smaller number of host metagenomes (Fig. 1F). These large sample sizes enabled our between-host population genetic analyses in Fig. 2-Fig. 4. In principle, many of these analyses could be performed equally well using traditional isolate-based approaches, in cases where comparably large numbers of isolates have been sequenced. However, as noted by Ref. [35], there are currently few isolate sequences available for many of the most prevalent human gut bacteria. To validate our approach, we therefore repeated our between-host analyses for the handful of species in Fig. 1F where larger sample sizes are available.

We downloaded isolate genomes from the PATRIC database [99] (as of May 1, 2018) that were annotated as belonging to one of three bacterial species: *Bacteroides vulgatus* (*n* = 15 genomes), *Bacteroides fragilis* (*n* = 107), and *Parabacteroides distastonis* (*n* = 17). Of these, only *B. fragilis* has a sample size approaching those available in our metagenomic study. We simulated metagenomic reads from each these isolate genomes at 100x coverage using the software Grinder [104]. These synthetic metagenomes therefore constitute simple versions of the QP samples we have analyzed above. We processed these synthetic metagenomes using the same MIDAS-based pipeline described in SI Section S1 Text, and we repeated our between-host analyses using the same code that we used to analyze the true metagenomic samples in the main text. The results largely recapitulate our findings in the main text (Fig. S13), particularly the observation of recombination within genes (compare Figs. S13D and S12A). This provides an important validation of our quasi-phasing approach.

We note, however, that we observe a somewhat larger number of closely related strains among the *B. fragilis* isolates than among the quasi-phased samples in Fig. 2. This could arise from the fact that many of the isolates in the PATRIC database were collected from a study in the same hospital [57], where they are more likely to have arisen from the same clonal expansion. This highlights the benefits of the large cohort studies that we have utilized (e.g., Refs. [42, 44, 45]), which were designed with an eye toward obtaining a representative random sample from a population.

### S8 Text

#### Quantifying prevalence of within-host SNV and gene changes

##### S8.1 Excess of high-prevalence SNVs

To interpret the SNV prevalence distribution in Fig. 5C, we compared the observed data to a null model of random *de novo* mutation. In such a model, within-host SNVs are assumed to occur uniformly along the genome of the resident population. If the resident population is fixed for the cohort-wide consensus allele, than the derived allele of the within-host sweep will be the cohort-wide minor allele, whose prevalence we denote by *p*_*i*_. On the other hand, if the resident population is fixed for the cohort-wide minor allele, than the derived allele of the within-host sweep will be the cohort-wide consensus, which has prevalence 1 – *p*_*i*_. To a first approximation, a random resident population can be formed by replacing the consensus genotype at each site with the cohort wide minor allele with probability *p*. Thus, under a model of random *de novo* mutation, the null distribution of prevalence is given by

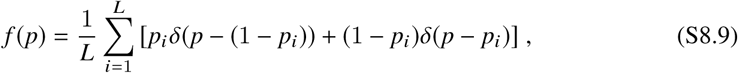

where δ (·) is the Dirac delta function, and the sum is over all *L* sites in the genome.

To compare this model with the observed data, we generated null expectations for the prevalence bins in Fig. 5C, using the database of private SNVs to populate the first and last bins. Different species genomes were weighted according to the number of within-host SNV differences observed in each species. Under the null hypothesis, the observed counts follow a multinomial distribution with these expected weights. We quantified deviations from this null model using the log-likelihood of the observed data as our test statistic:

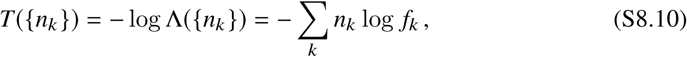

where *n*_*k*_ denotes the observed number of SNVs in prevalence bin *k*, and *f*_*k*_ denotes the expected weight in that bin. Significance was assessed numerically by resampling the null distribution for *n* = 10^4^ bootstrap iterations, and calculating the fraction of bootstrap samples with *T* greater than or equal to the observed value.

Null distributions for the prevalence of gene gains and losses are obtained using a similar procedure. We assume that *de novo* mutations cannot produce a gene gain by definition, so we only consider the distribution of prevalence within the set of gene losses. We assume that random *de novo* gene losses occur uniformly throughout the genome of the resident population, and that a given gene is present in the resident population with probability proportional to its prevalence *p*_*i*_. The null distribution for gene loss prevalence is therefore given by

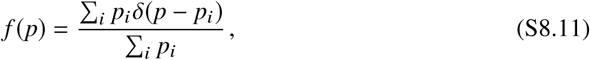

where the sum is over all genes in a species’ pangenome. The null expectations in Fig. 5D are obtained by summing this null distribution within each prevalence bin, and multiplying by the same total number of losses.

##### S8.2 Non-uniform distribution of synonymous and nonsynonymous mutations

To quantify the relationship between prevalence and the inferred strength of natural selection, we examined the differences in the relative fraction of synonymous (4D) and nonsynonymous (1D) in the different prevalence bins in Fig. S16C. We compared the observed distribution against a null model prevalence and amino acid impact are independent of each other. The null model is chosen so that it has the same overall prevalence distribution and fraction of nonsynonymous and synonymous mutations as the observed data. If we let 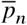 denote the fraction of nonsynonymous mutations across all prevalence bins, then under the null model, the number of nonsynonymous mutations in bin 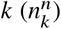 should be binomially distributed with success probability 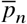 As above, we quantified deviations from this model using the log-likelihood as a test statistic,

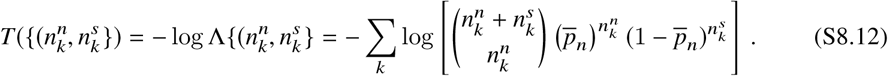

Significance was assessed numerically by resampling the null distribution for *n* = 10^4^ bootstrap iterations, and calculating the fraction of bootstrap samples with *T* greater than or equal to the observed value.

To demonstrate this result is robust to the choice of prevalence bins, we directly compared the raw prevalence values of synonymous and nonsynonymous mutations using the Kolmogorov-Smirnov (KS) test [111]. In particular, we calculated the KS distance *D* between the empirical prevalence distributions of synonymous and nonsynonymous mutations (Fig. S17B). To assess significance, we compared the observed value of *D* against a null model where the synonymous and nonsynonymous labels are randomly permuted across the different prevalence values. We performed *n* = 10^4^ bootstrap iterations, and calculated a *P*-value as the fraction of bootstrap samples with *D* greater than or equal to the observed value.

##### S8.3 Time-reversal asymmetry

To provide further support for the hypothesis that modification events represent evolutionary changes, we examined the temporal asymmetry of the prevalence distributions in Fig. 5C,D. If these genetic differences were primarily driven by equilibrium processes like (i) replacement by extremely closely related strains or (ii) bioinformatic artifacts like read donating described in SI Section S1 Text, then the statistical features of these changes should be independent of the labeling of the initial and final timepoints. This is a form of *time-reversal symmetry* [112]. In contrast, evolutionary processes often violate time-reversal symmetry, particularly when natural selection is involved, since less-fit ancestors are continually replaced by fitter descendents (and only rarely the other way around).

To see how time-reversal symmetry applies in the context of Fig. 5, we note that if we reverse the intial and final timepoints, then gene gains become gene losses and vice versa, while their prevalence values (and the overall number of gene changes) are preserved. Similarly, for within-host SNV differences, reversing the order of time switches the roles of the ancestral and derived alleles, so that the prevalence of the derived allele switches from *p →*1 *- p*. Thus, reversing the order of time reflects the distributions in Fig. 5C,D across the central axis of each panel. Time-reversal symmetry therefore requires that these prevalence distributions are symmetric about this central axis.

We tested for violations of time-reversal symmetry using a Kolmogorov-Smirnov (KS) procedure [111], similar to the one employed in Text S8.2. For the SNVs in Fig. 5C, we calculated the KS distance *D* between the observed distribution of (unbinned) prevalence values, and a corresponding symmetrized version, in which every prevalence value *p* is duplicated with its time-reflected value 1 – *p* (Fig. S17A). To assess significance, we compared the observed value of *D* against a null model in which the initial and final timepoints of each resident population are randomly permuted. We carried out this procedure for *n* = 10^4^ bootstrap iterations, and calculated a *P*-value as the fraction of bootstrap samples with *D* greater than or equal to the observed value. We used a similar procedure to test for deviations of time-reversal symmetry for the gene gains and losses in Fig. 5C, except with the KS distance *D* calculated using the prevalence distributions of gains and losses (Fig. S17C). For both SNV and gene changes, we observed significant deviations from the null model of time-reversal symmetry (*P* < 10^-4^ and *P* ≈ 2 × 10^-3^, respectively). This suggests that evolutionary processes, rather than strain replacement or bioinformatic errors, provide a better explanation for the data.

**Fig S1.**
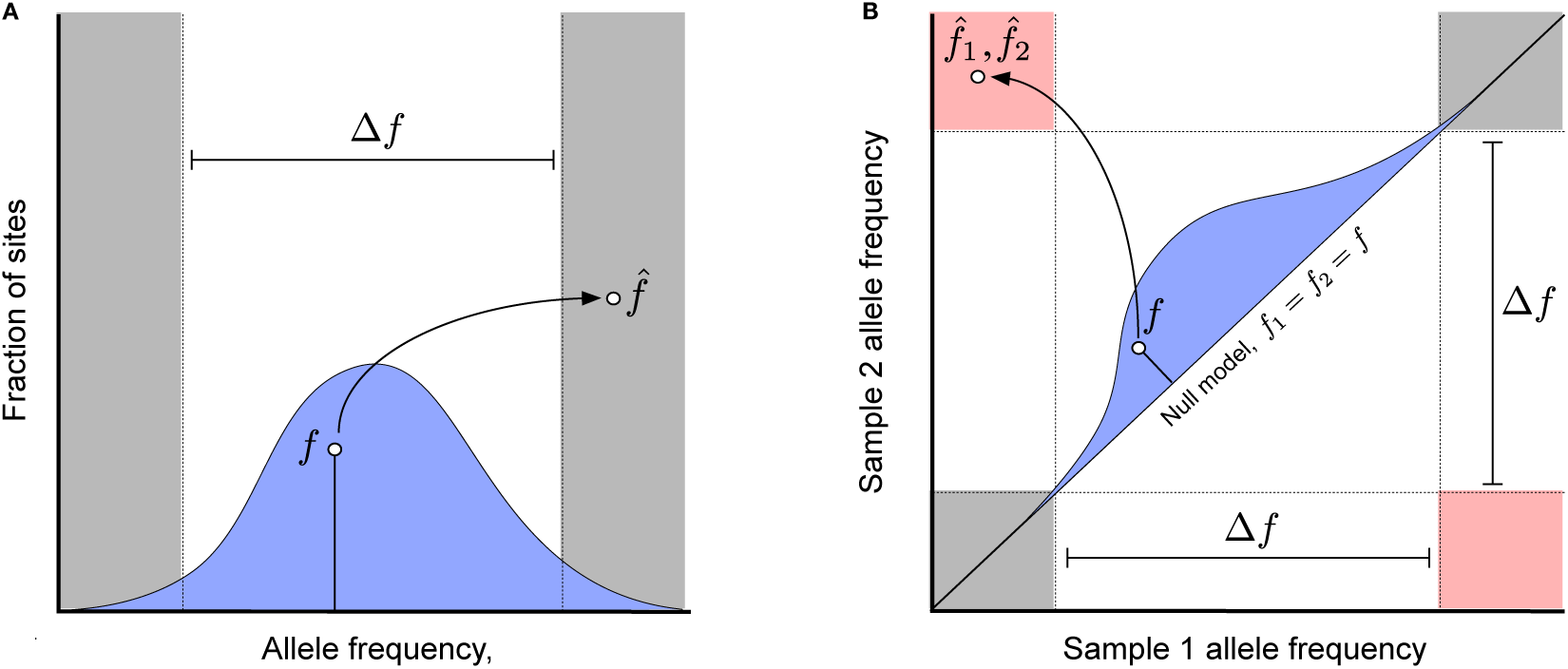
Schematic depiction of phasing and substitution errors. (a) An example of a haplotype phasing error, where an allele with true within-host frequency *f* [drawn from a hypothetical genome-wide prior distribution, *p*_0_ (*f*), blue] is observed with a sample frequency 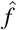 with the opposite polarization. (b) An example of a falsely detected nucleotide substitution between two samples, where an allele with true frequency *f*_1_ = *f*_2_ = *f*[drawn from a hypothetical genome-wide null distribution, *p*_0_ (*f*), blue] is observed with a sample frequency 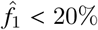 in one sample and 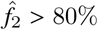 in another. Allele frequency pairs that fall in the pink region are counted as nucleotide differences between the two samples, while pairs in the grey shaded region are counted as evidence for no nucleotide difference; all other values are treated as missing data.

**Fig S2.**
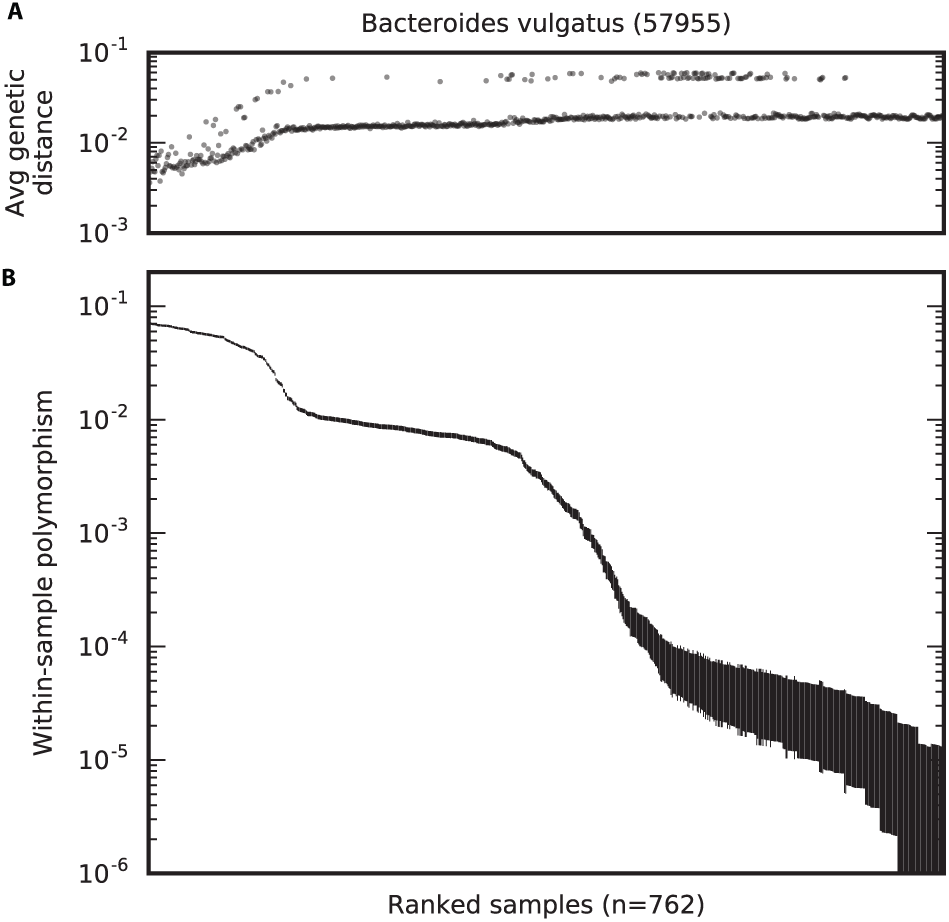
Average genetic distance between *B. vulgatus* metagenomes. (a) The fraction of fourfold degenerate synonymous sites in the core genome that have major allele frequencies ≥ 80% and differ in a randomly selected sample (see Text S3.3 for a formal definition). (b) The corresponding rate of intermediate-frequency polymorphism for each sample, reproduced from Fig. 1B. In both panels, samples are plotted in the same order as in Fig. 1B.

**Fig S3.**
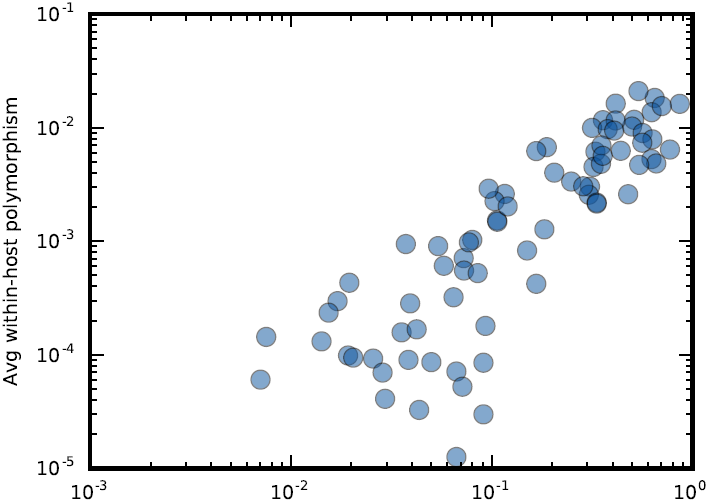
Correlation between within-host diversity and the fraction of non-QP samples per species. Circles denote the average rate of within-host polymorphism (as defined in Fig. 1E) for each species as a function of the fraction of non-QP samples in that species.

**Fig S4.**
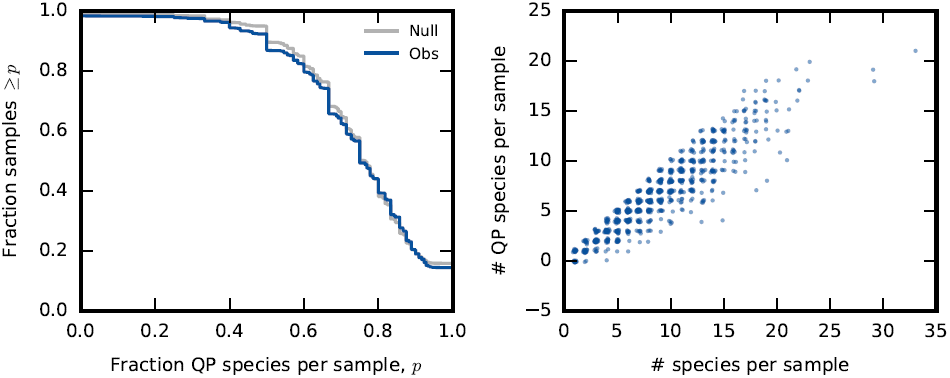
Distribution of the number of QP species per sample. Left: The distribution of the fraction of QP species per sample (blue line). The grey line denotes the corresponding null distribution obtained by randomly permuting the QP classifications across the samples. We conclude that QP species are not strongly enriched within specific hosts. Right: The number of species classified as QP in each sample on the left as a function of the number of species with sufficient coverage in that sample. A small amount of noise is added to both axes to enhance visibility.

**Fig S5.**
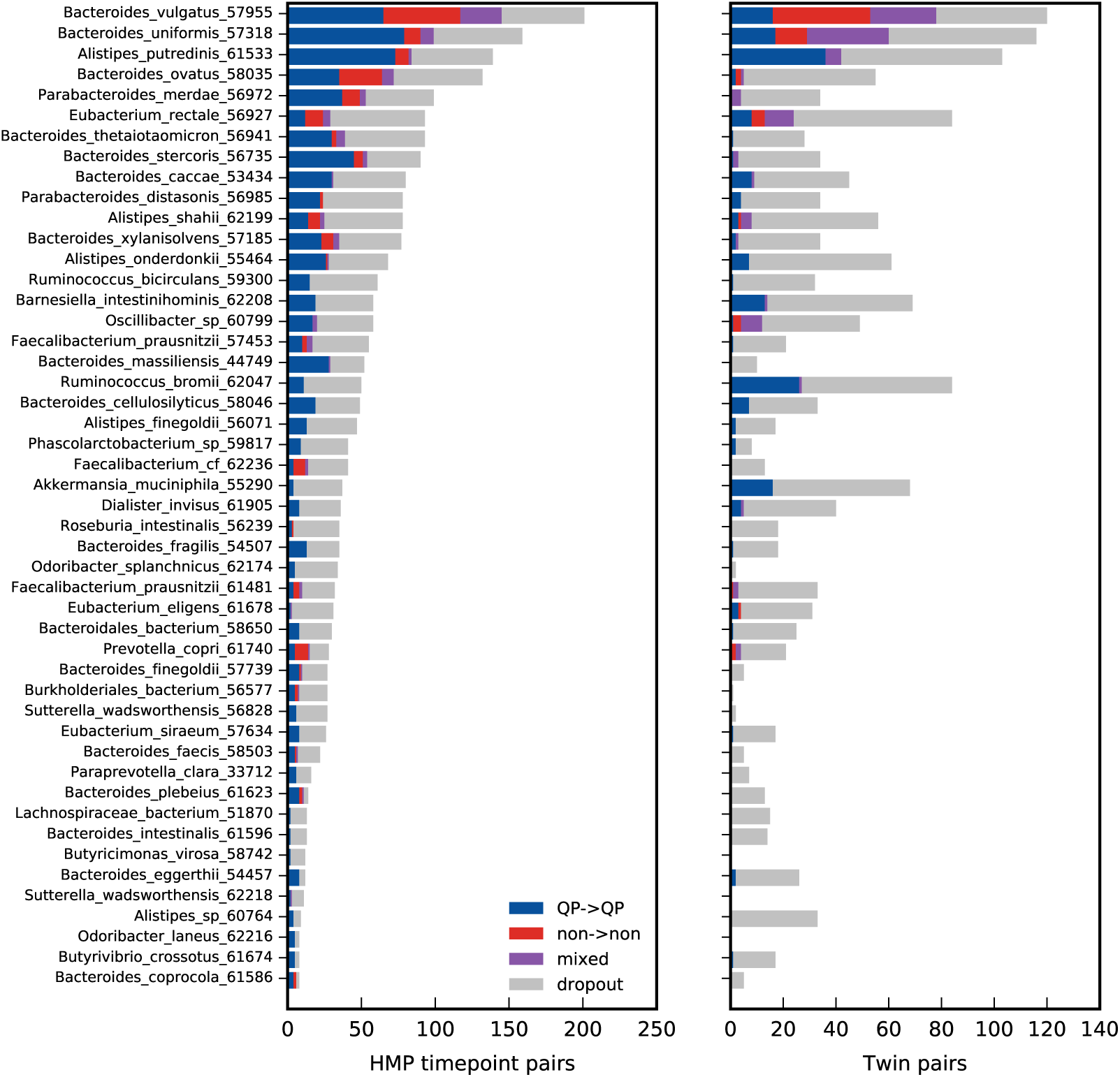
Distribution of quasi-phaseable (QP) samples in longitudinal samples and adult twin pairs. Bars show the number of sample pairs for each species that are quasi-phaseable for both samples (QP→QP), non-quasi-phaseable for both samples (non→non), mixed samples (QP→non or non→QP), and pairs where the species did not have sufficient coverage in one of the two timepoints (dropout). The left panel shows data from longitudinally sampled individuals in the HMP cohort [42, 44], while the right panel compares contemporary samples from pairs of adult twins [45]. Species are ordered in decreasing order of prevalence in the HMP cohort. Species are only included if they have at least 10 QP samples and at least 3 QP timepoint pairs.

**Fig S6.**
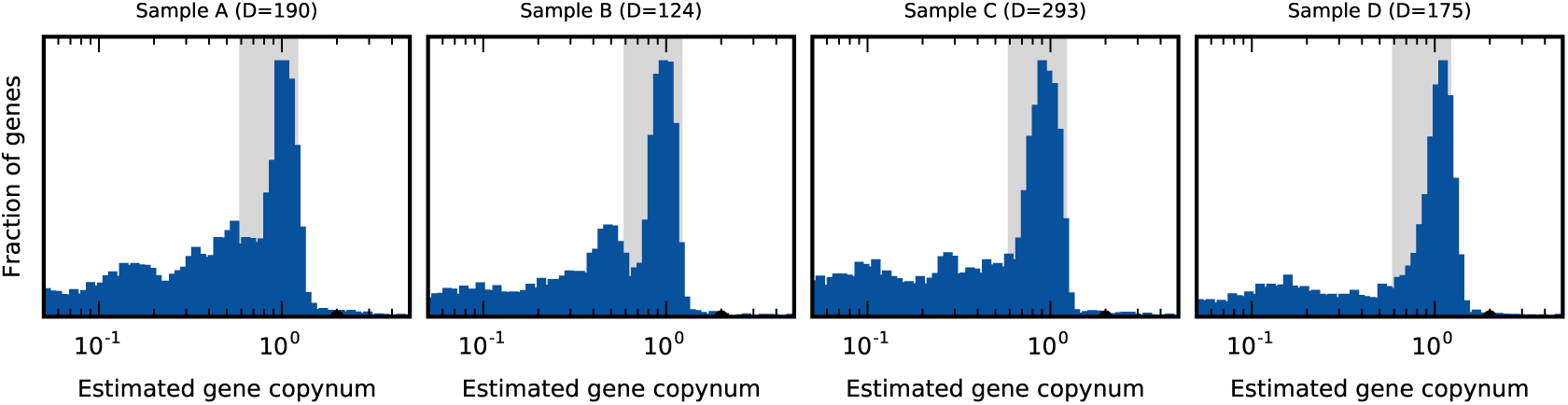
Distribution of estimated gene copy numbers for the four samples in Fig. 1. The grey region denotes the copy number range required in at least one sample to detect a difference in gene content between a pair of samples (see Text S3.5).

**Fig S7.**
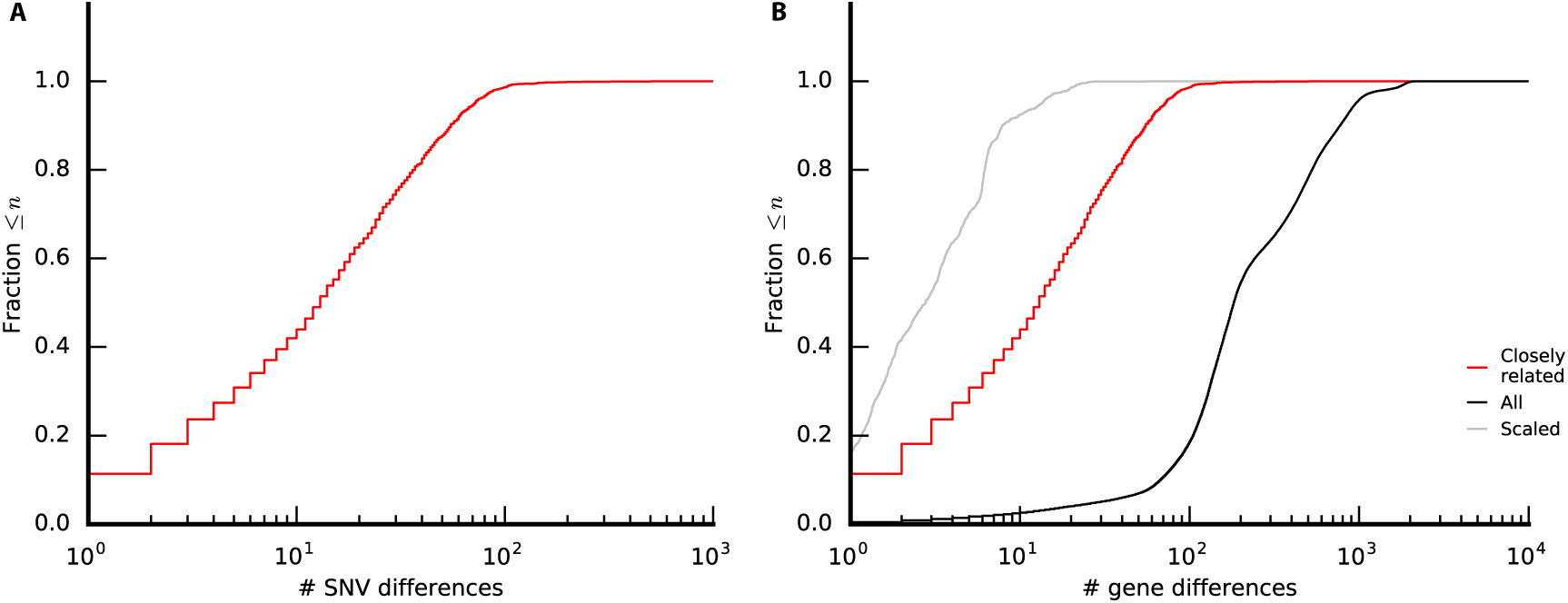
SNV and gene content differences between closely related strains. (a) Cumulative distribution of the total number of core genome SNV differences between closely related strains in Fig. 2. (b) Cumulative distribution of the number of gene content differences for the closely related strains in panel a (red line). For comparison, the corresponding distribution for all pairs of strains in Fig. 2 is shown in black, while the grey line denotes a ‘clock-like’ null distribution for the closely related strains, which assumes that genes and SNVs each accumulate at constant rates.

**Fig S8.**
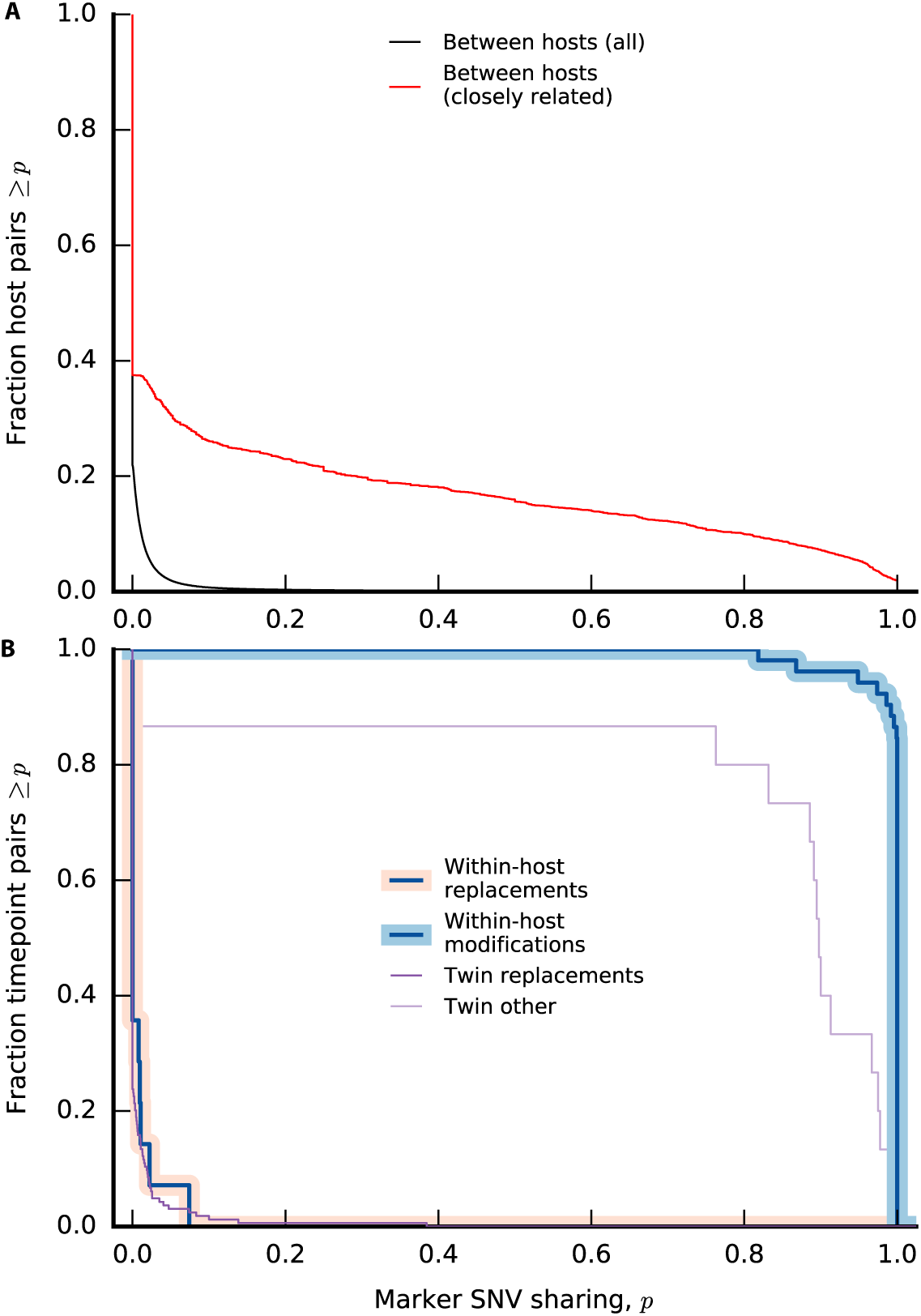
Private marker SNV sharing within and between hosts. Given an ordered pair of QP strains, we define private marker SNVs to be core genome SNVs that (i) are phaseable in both strains, (ii) have the derived allele in strain 1, and (iii) do not have the derived allele in any other host outside the pair. The marker sharing fraction *p* is then defined as the fraction of private marker SNVs that also have the derived allele in strain 2. (a) Private marker SNV sharing between unrelated hosts. Solid lines show the distribution of marker sharing fraction *p* between all pairs of strains in Fig. 2 (black) and between the subset of closely related strains (red). Separate sharing fractions are calculated for both orderings of a given strain pair, and we only include pairs with at least 10 marker SNVs. (b) Distribution of marker SNV sharing for replacement and modification events in longitudinally sampled HMP hosts (blue lines), using the replacement and modification thresholds in Fig. 5A. For comparison, the distribution of marker SNV sharing between strains in pairs of adult twins is shown in purple. For twins, we use modified definitions of replacement (>10^3^ SNV differences) and modification (<10^3^ SNV differences). As above, sharing fractions are only computed for samples with at least 10 marker SNVs.

**Fig S9.**
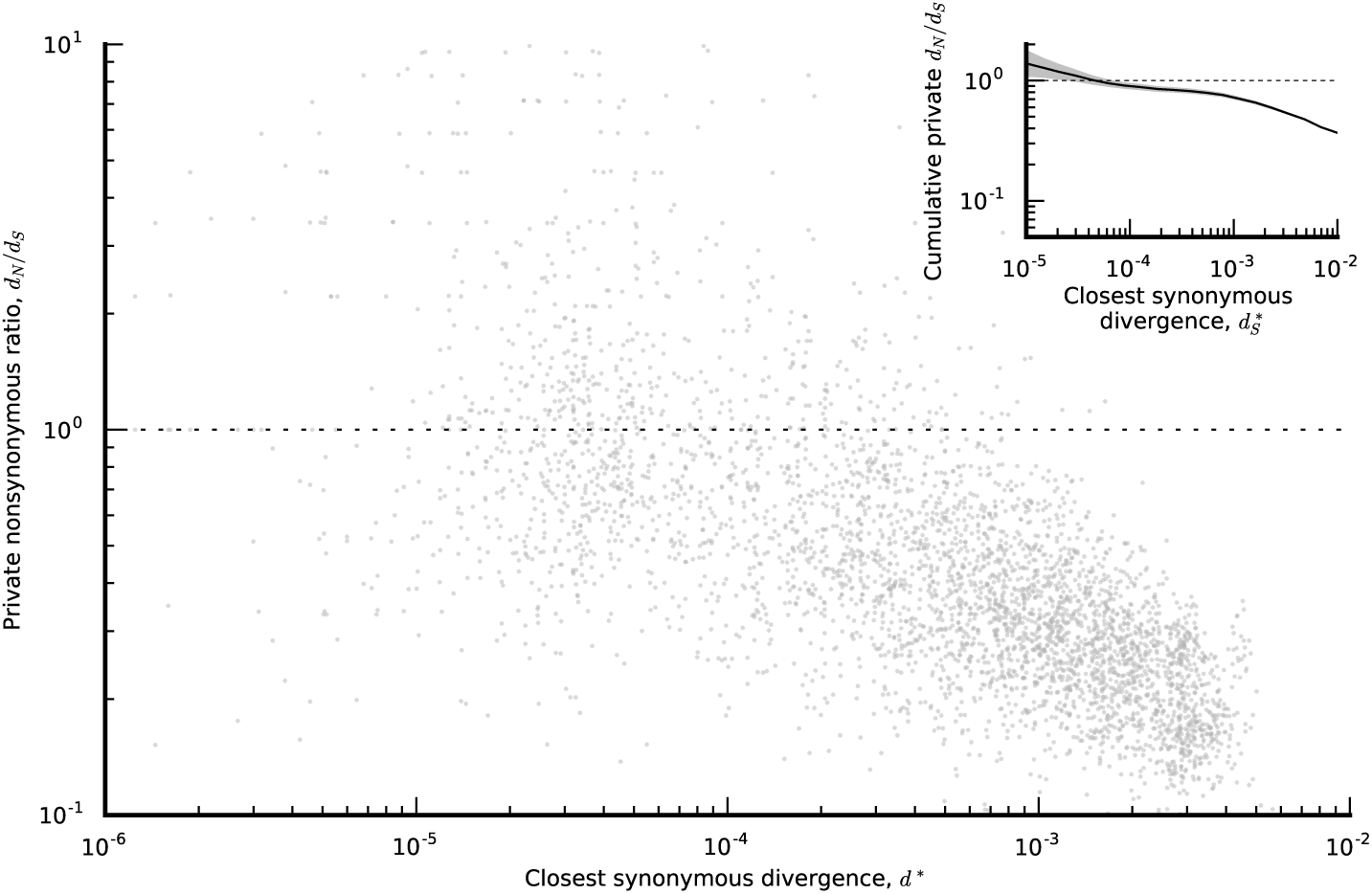
Signatures of selective constraint within private SNVs. An analogous version of Fig. 3 computed for private SNVs. For each QP species x host combination, *d*_*N*_/*d*_*S*_ is computed for the subset of alleles that are not found in any other hosts. These private *d*_*N*_/*d*_*S*_ ratios are plotted as a function of 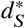, an estimate of the minimum synonymous divergence from other QP lineages of that species. (inset) Ratio between the cumulative *d*_*N*_ and *d*_*S*_ values for all lineages with 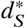 less than the indicated value. The narrow shaded region denotes 95% confidence intervals estimated by Poisson resampling. The resampling procedure uses an analogous version of the thinning scheme employed in Fig. 3 to ensure that the *x* and *y* axes are statistically independent (see SI Section S4 Text).

**Fig S10.**
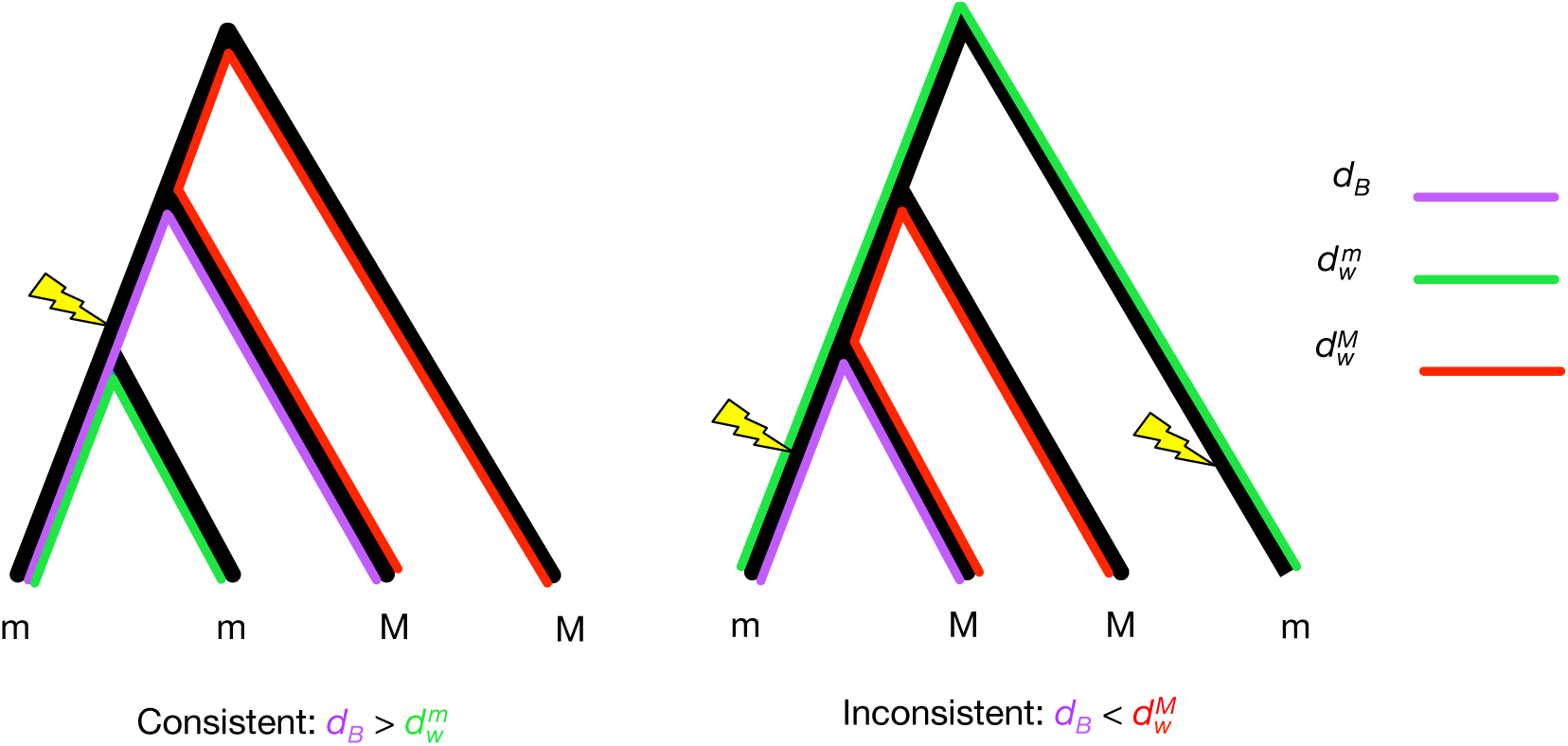
Schematic illustration of phylogenetic inconsistency between individual SNVs and core-genome-wide divergence. Two examples are shown, illustrating phylogenetically consistent and inconsistent SNVs, respectively, in a sample of four lineages. The lineages at the leaves of each tree are labeled according to whether they have the major (*M*) or minor (*m*) allele. Thunderbolts depict the most parsimonious introduction of the derived allele on the genealogy. Different colors indicate the core-genome-wide divergence between lineages with different combinations of alleles, as described in Text S5.1. Highlighted in purple is *d*_*B*_, which is the minimum divergence between two lineages bearing different alleles. Highlighted in red and green are 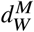 and 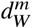, which are the maximum divergence between individuals bearing the same allele (major and minor, respectively). In practice, we do not know which allele is ancestral and which is derived, so we define 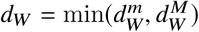. If *d*_*W*_ ≫ *d*_*B*_, we say that the SNV is phylogenetically inconsistent.

**Fig S11.**
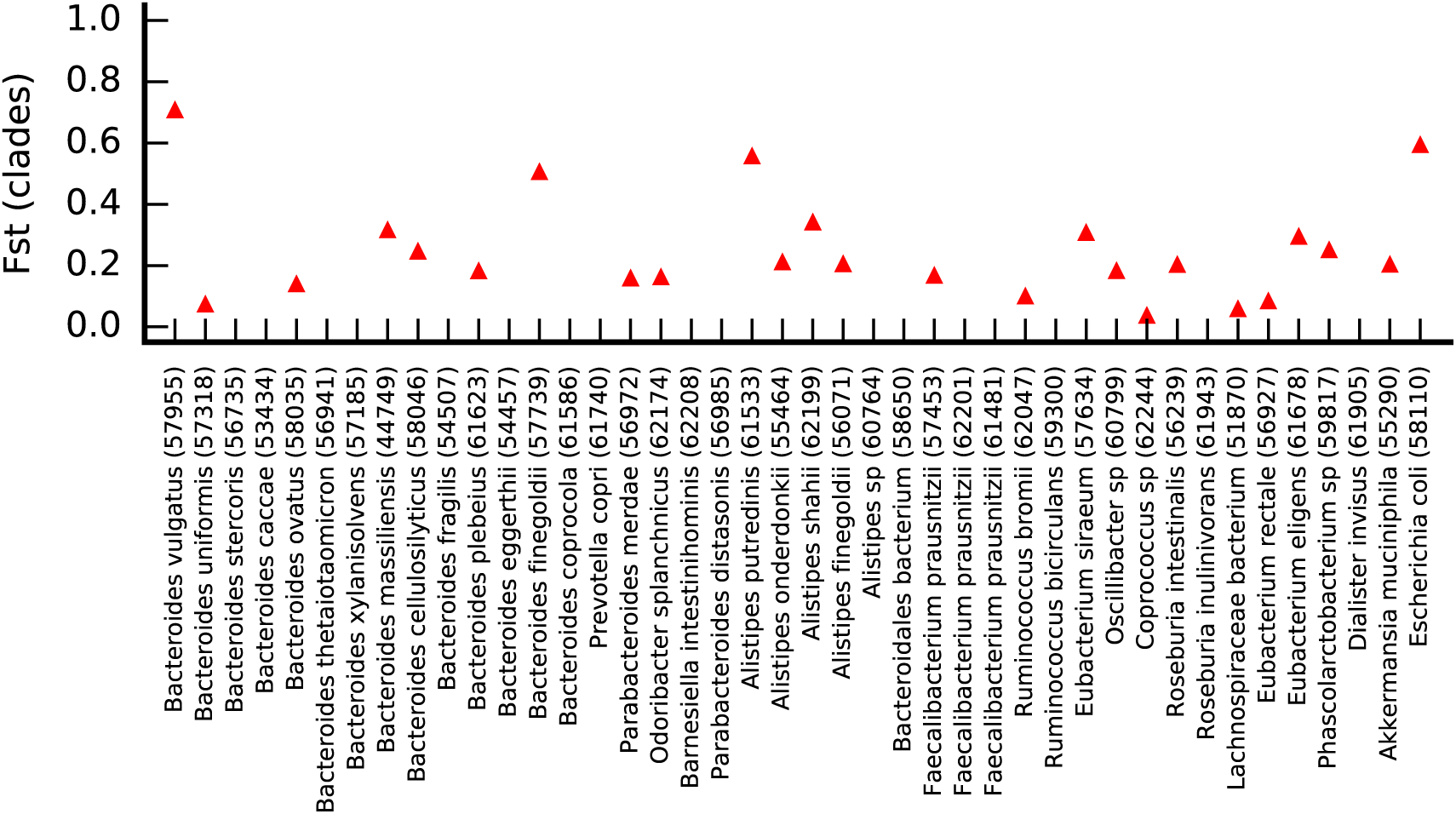
Top-level clade structure among lineages in different QP hosts. Core-genome-wide *F*_*st*_ between manually assigned top-level clades in each species (Table S2, Text S5.2). Species are only included if there are at least two clades with more than two individuals in each of them.

**Fig S12.**
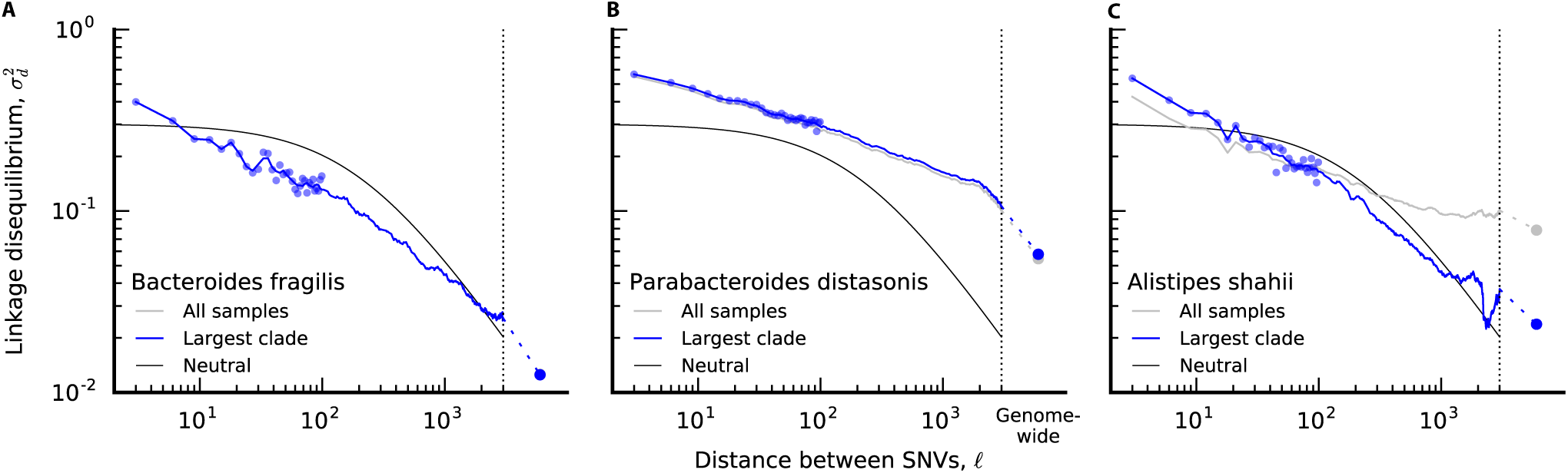
Decay of linkage disequilibrium in three example species. Analogous versions of Fig. 4B for *Bacteroides fragilis, Parabacteroides distasonis* and *Alistipes shahii*.

**Fig S13.**
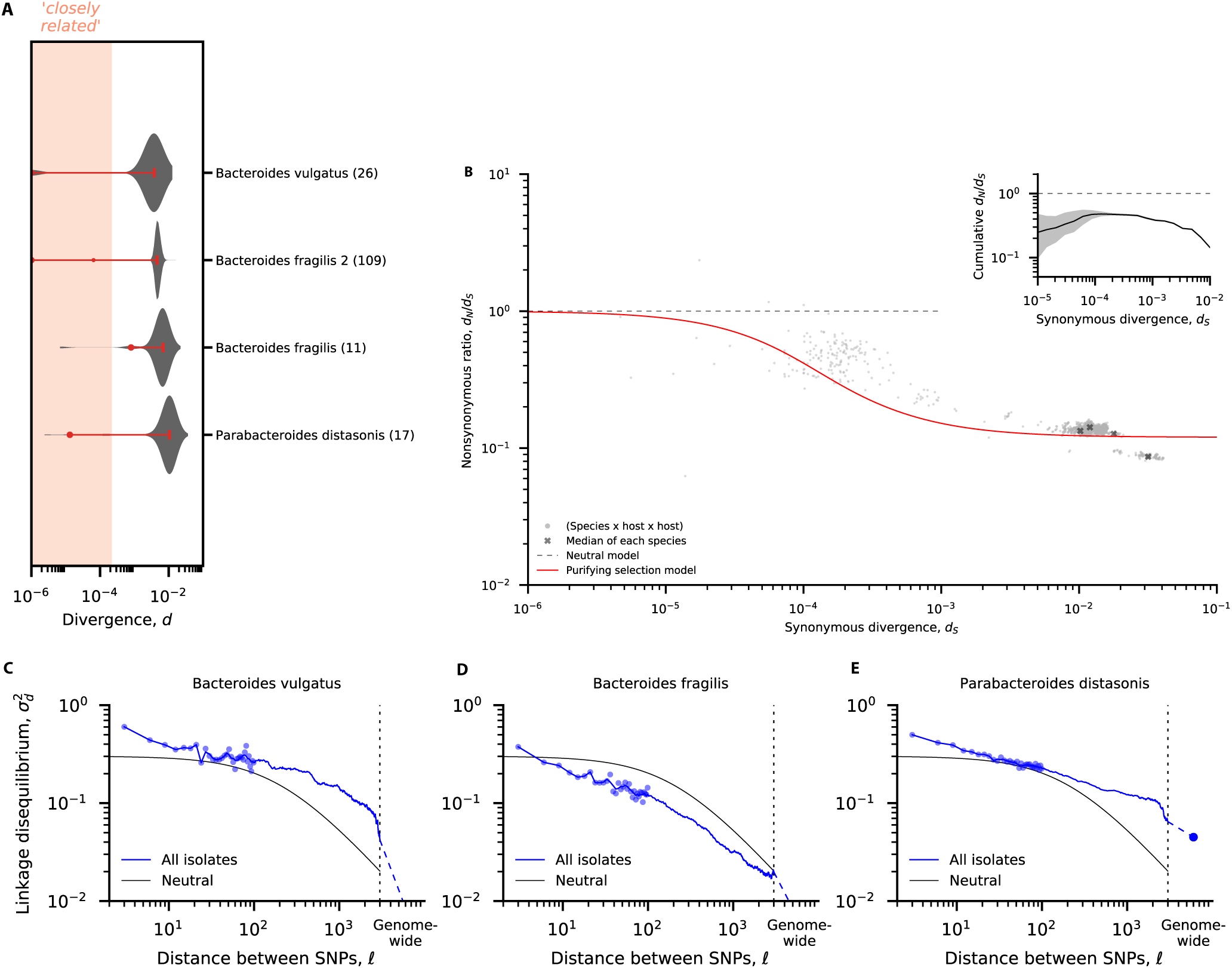
Recapitulating patterns of between-host evolution from sequenced isolates. (a) An analogous version of Fig. 2B constructed from the genomes of sequenced isolates in three representative species, as described in SI Section S7 Text. (b) An analogous version of Fig. 3 constructed from the pairs of isolate genomes in panel a. (c-e) Analogous versions of Fig. 4B for the three species in (a) with the largest number of isolates.

**Fig S14.**
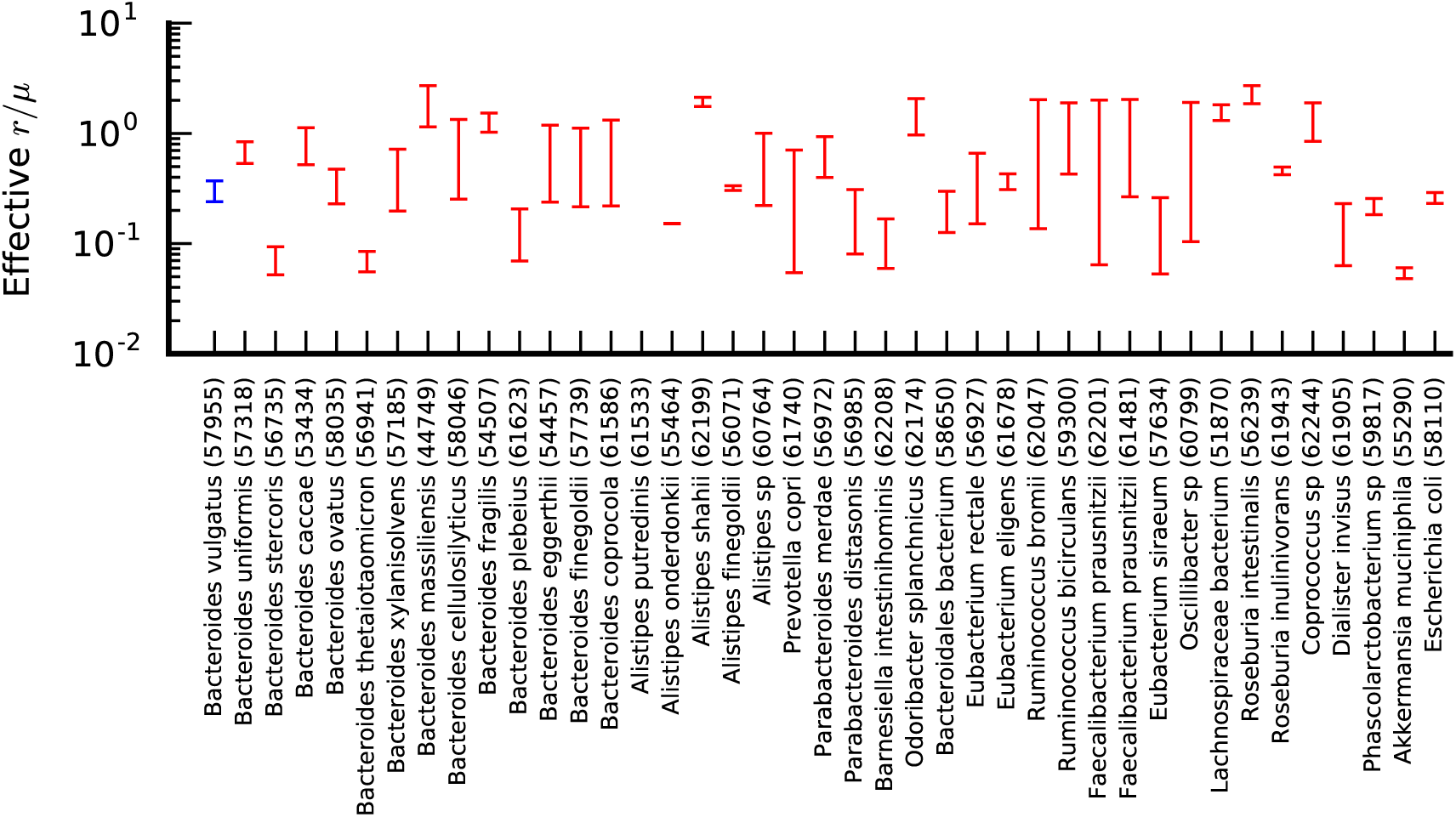
Recombination rate estimates based on the decay of linkage disequilibrium. For each species, the two dashes represent effective values of *r*/µ estimated from the neutral prediction for the decay of 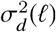, using the half-maximum and quarter-maximum decay lengths (see SI Section S6 Text). The two estimates are connected by a vertical line for visualization.

**Fig S15.**
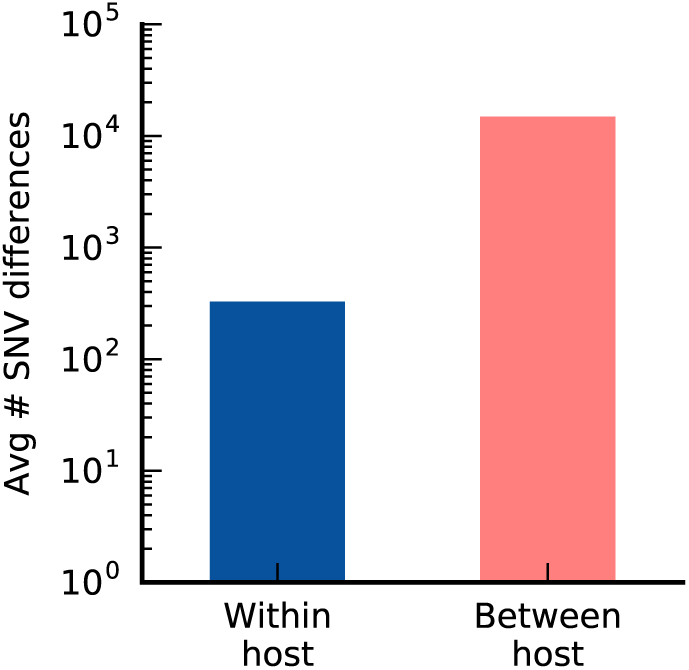
Average number of SNV differences within and between hosts. Blue and red bars denote the average of the within- and between-host distributions in S16A. Consistent with previous work [31, 44], the within-host average is ∼100-fold lower than the between-host average. However, the average is a poor summary of the *typical* values in the between-host distribution in Fig. S16A. Instead, the within-host average is well approximated by the product of the typical number of SNV differences per replacement, and the overall fraction of replacement events.

**Fig S16.**
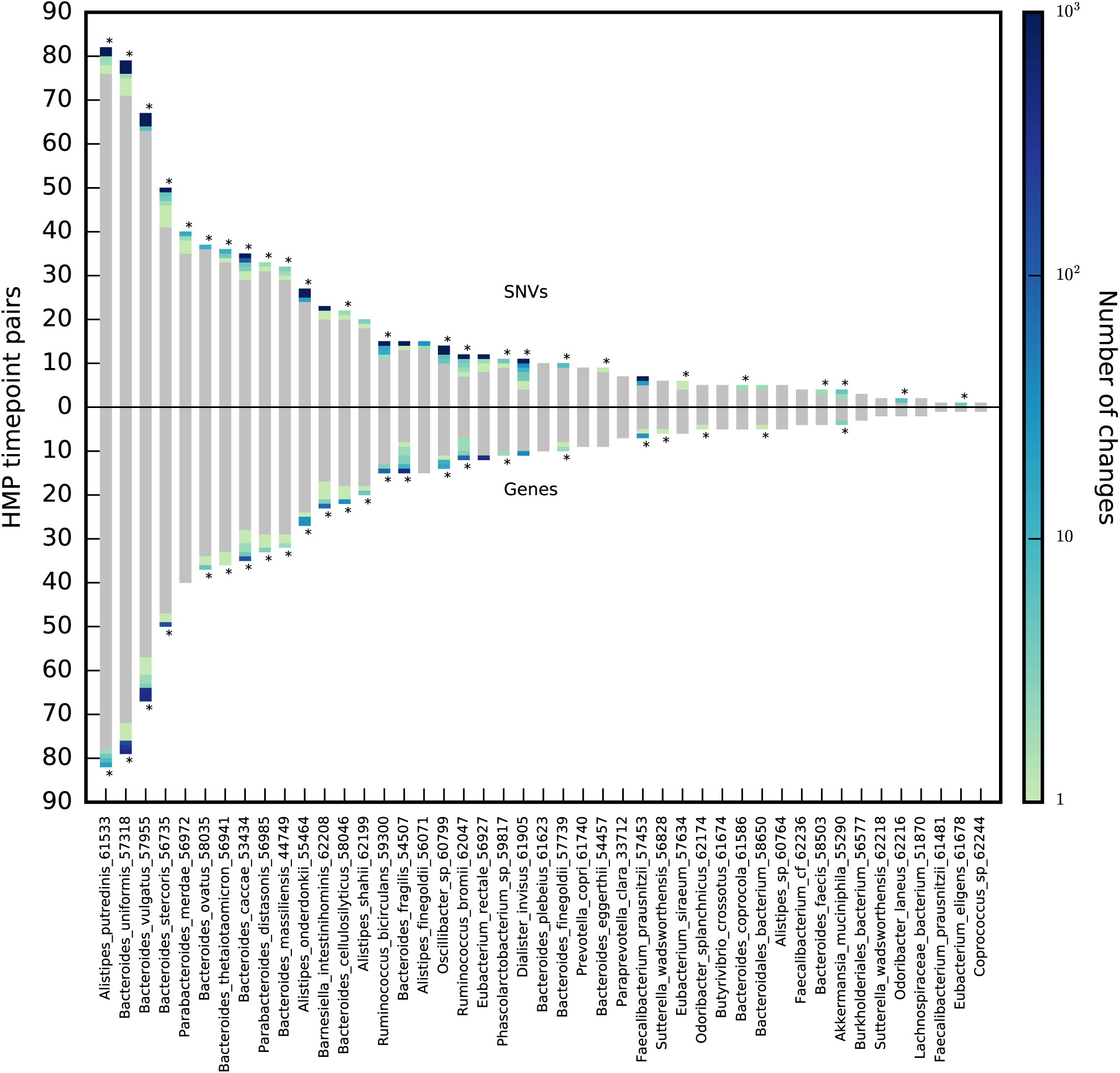
Comparable rates of within-host SNV and gene changes across prevalent species. Summary of within-host SNV changes (top) and gene changes (bottom) across all species with at least 10 QP samples and at least 3 pairs of longitudinal QP samples. Each row in each bar represents a different longitudinal pair from the HMP cohort, and rows are colored according to the total number SNV changes (top) and gene changes (bottom), with grey indicating no detected changes. A star indicates that the total number of non-replacement changes is ≥ 10 times the total estimated error rate across samples from that species (see Text S3.4 and S3.5), where replacements are defined as in Fig. 5.

**Fig S17.**
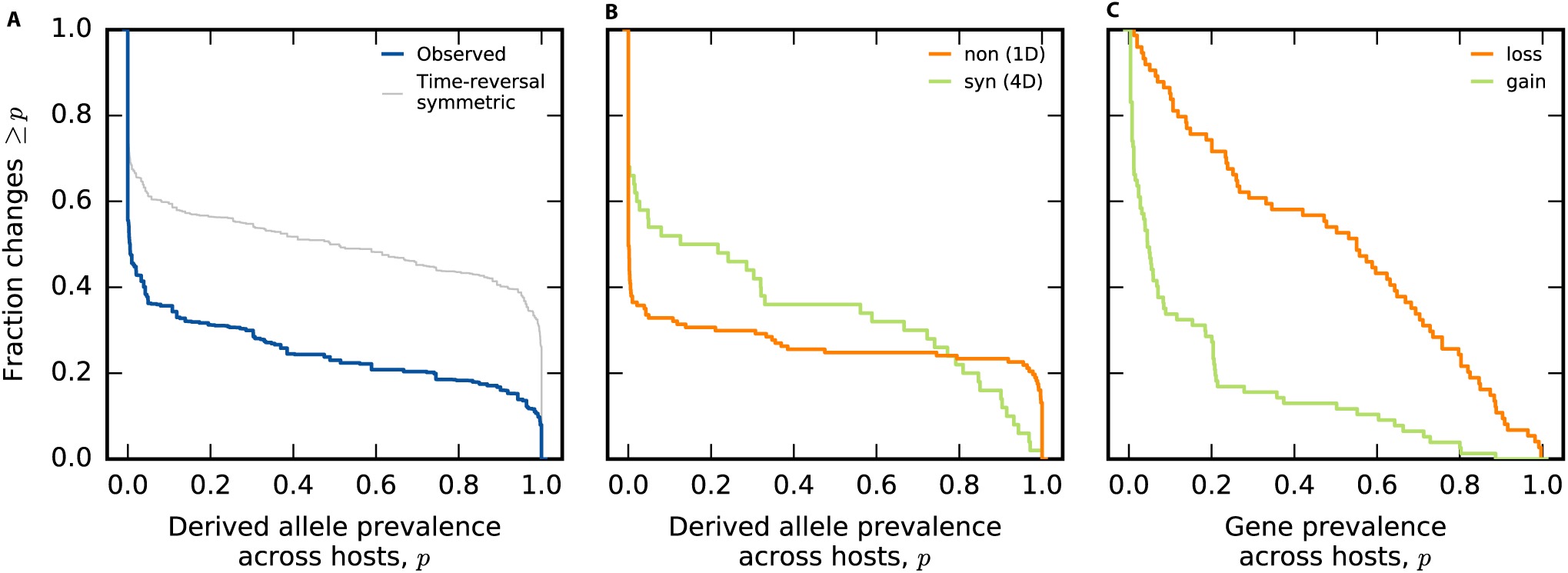
Prevalence distributions of within-host SNV and gene content differences without binning. (a) The empirical survival function for the raw prevalence values Fig. 5C. For comparison, the grey line shows the time-reversal symmetric version described in Text S8.3. (b) Empirical prevalence distributions for synonymous (1D) and nonsynonymous (4D) differences in Fig. 5C. (c) Empirical prevalence distributions for gene gains and losses in Fig. 5D.

**Fig S18.**
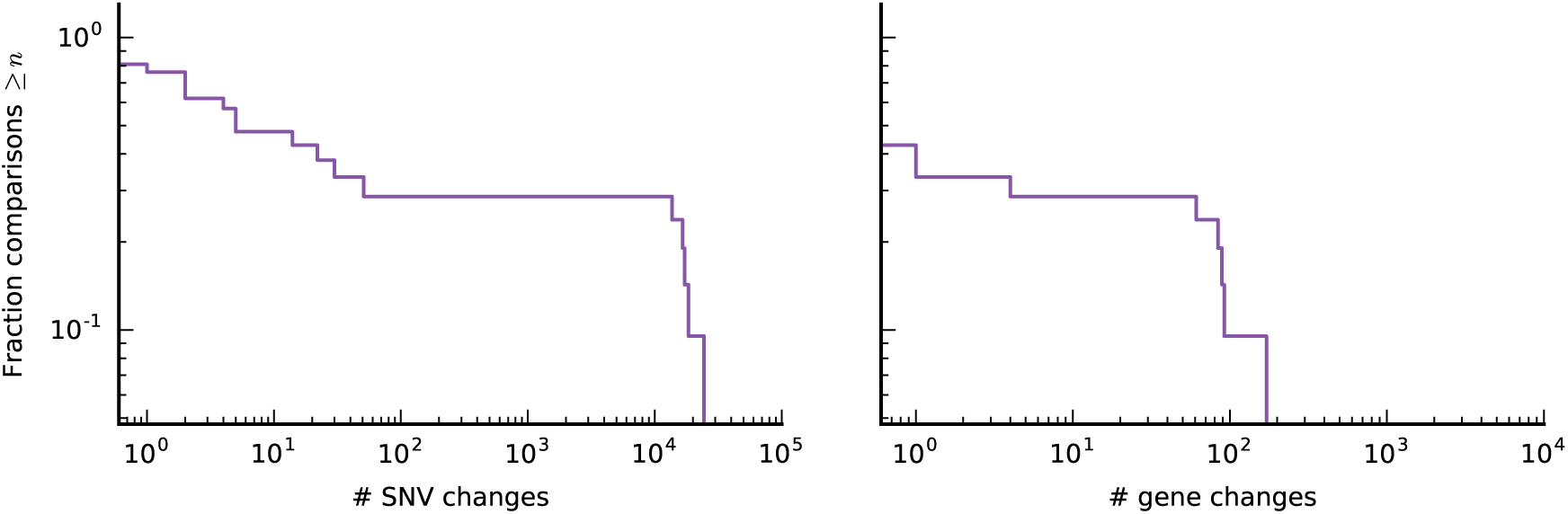
SNV and gene content differences between younger twins. Analogous versions of Fig. 5A,B for four twin pairs from Ref. [46], which range from ∼ 5 to ∼ 20 years of age. The results are consistent with the original findings in Ref. [46].

**Fig S19.**
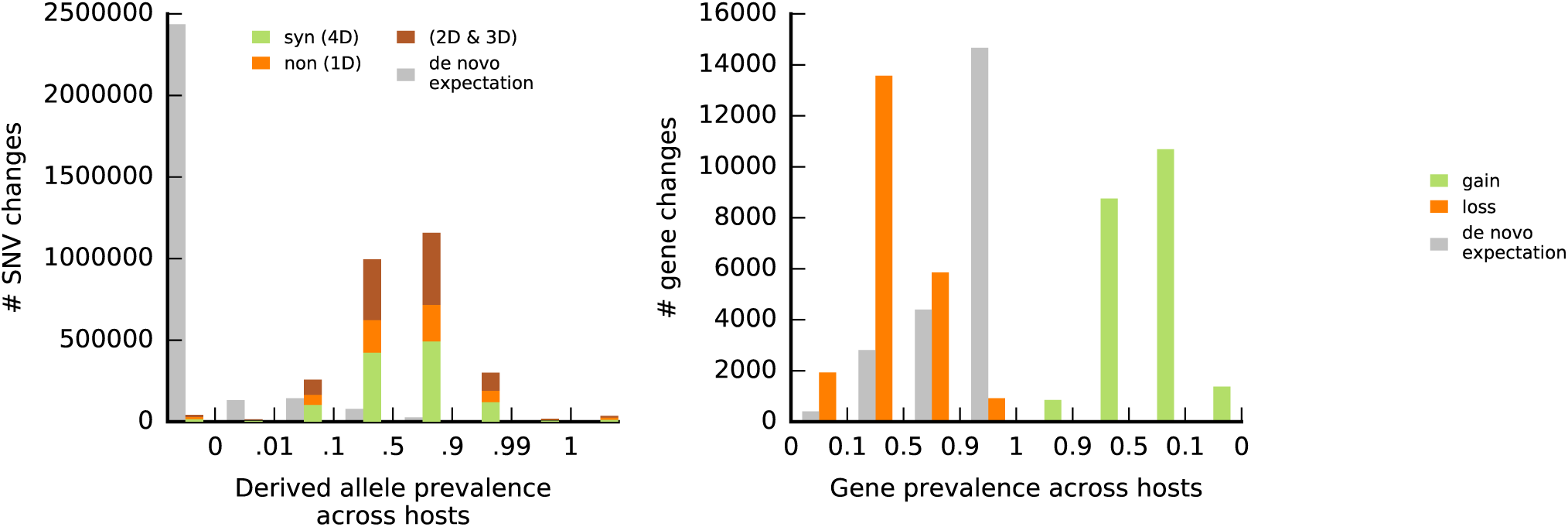
Prevalence of SNV and gene content differences between adult twins. Analogous versions of Fig. 5C,D computed using the SNV and gene content differences observed between the adult twins (purple lines in Fig. 5A,B). In contrast to the within-host changes in Fig. 5C,D, the prevalence distributions and the relative fraction of nonsynonymous differences between twins are more consistent with replacement by a distantly related strain.

## References

1. David LA, Maurice CF, Carmody RN, Gootenberg DB, et al. Diet rapidly and reproducibly alters the human gut microbiome. Nature. 2013;505:559–563.

2. Seedorf H, Griffin NW, Ridaura VK, Reyes A, et al. Bacteria from Diverse Habitats Colonize and Compete in the Mouse Gut. Cell. 2014;159:253–266.

3. Rakoff-Nahoum S, Foster KR, Comstock LE. The evolution of cooperation within the gut microbiota. Nature. 2016;533:255–259.

4. Verster AJ, Ross BD, Radey MC, Bao Y, et al. The Landscape of Type VI Secretion across Human Gut Microbiomes Reveals Its Role in Community Composition. Cell Host & Mirobe. 2017;22:411–419.

5. Bradley PH, Nayfach S, Pollard KS. Phylogeny-corrected identification of microbial gene families relevant to human gut colonization. biorXiv. 2017;.

6. Matamouros S, Hayden HS, Hager KR, Brittnacher MJ, Lachance K, Weiss EJ, et al. Adaptation of commensal proliferating Escherichia coli to the intestinal tract of young children with cystic fibrosis. Proceedings of the National Academy of Sciences. 2018; p. 201714373.

7. Barroso-Batista J, Sousa A, Lourenço M, Bergman ML, Sobral D, Demengeot J, et al. The first steps of adaptation of Escherichia coli to the gut are dominated by soft sweeps. PLoS genetics. 2014;10(3):e1004182.

8. Barroso-Batista J, Demengeot J, Gordo I. Adaptive immunity increases the pace and predictability of evolutionary change in commensal gut bacteria. Nature Communications. 2015;6:8945.

9. Lescat M, Launay A, Ghalayini M, Magnan M, Glodt J, Pintard C, et al. Using long-term experimental evolution to uncover the patterns and determinants of molecular evolution of an Escherichia coli natural isolate in the streptomycin-treated mouse gut. Molecular ecology. 2017;26(7):1802–1817.

10. Goodman AL, McNulty NP, Zhao Y, Leip D, et al. Identifying Genetic Determinants Needed to Establish a Human Gut Symbiont in Its Habitat. Cell Host & Microbe. 2009;6:279–289.

11. Wu M, McNulty NP, Rodionov DA, Khoroshkin MS, et al. Genetic determinants of in vivo fitness and diet responsiveness in multiple human gut Bacteroides. Science. 2015;350:aac5992.

12. Didelot X, Walker AS, Peto TE, Crook DW, Wilson DJ. Within-host evolution of bacterial pathogens. Nat Rev Microbiol. 2016;14:150–162.

13. Jerison ER, Desai MM. Genomic investigations of evolutionary dynamics and epistasis in microbial evolution experiments. Current Opinion in Genetics & Development. 2015;35:33–39.

14. Denef VJ, Banfield JF. In Situ Evolutionary Rate Measurements Show Ecological Success of Recently Emerged Bacterial Hybrids. Science. 2012;336:462–466.

15. Shapiro BJ, Friedman J, Cordero OX, Preheim SP, et al. Population genomics of early events in the ecological differentiation of bacteria. Science. 2012;336:48–51.

16. Rosen MJ, Davison M, Bhaya D, Fisher DS. Fine-scale diversity and extensive recombination in a quasisexual bacterial population occupying a broad niche. Science. 2015;348:1019–1023.

17. Bendall ML, Stevens SLR, Chan LK, Malfatti S, et al. Genome-wide selective sweeps and gene-specific sweeps in natural bacterial populations. The ISME Journal. 2016;10:1589–1601.

18. Herron MD, Doebeli M. Parallel Evolutionary Dynamics of Adaptive Diversification in *Escherichia coli*. PLoS Biology. 2013;11:e1001490.

19. Lang GI, Rice DP, Hickman MJ, Sodergren E, Weinstock GM, Botstein D, et al. Pervasive genetic hitchhiking and clonal interference in forty evolving yeast populations. Nature. 2013;500:571–574.

20. Tenaillon O, Barrick JE, Ribeck N, Deatherage DE, Blanchard JL, et al. Tempo and mode of genome evolution in a 50,000-generation experiment. Nature. 2016;536:165–170.

21. Lieberman TD, Flett KB, Yelin I, Martin TR, McAdam AJ, Priebe GP, et al. Genetic variation of a bacterial pathogen within individuals with cystic fibrosis provides a record of selective pressures. Nature Genetics. 2014;46:82–87.

22. Zanini F, Brodin J, Thebo L, Lanz C, Bratt G, Albert J, et al. Population genomics of intrapatient HIV-1 evolution. eLife. 2015;4:e11282.

23. Traverse CC, Mayo-Smith LM, Poltak SR, Cooper VS. Tangled bank of experimentally evolved Burkholderia biofilms reflects selection during chronic infections. Proc Natl Acad Sci USA. 2013;110:E250–E259.

24. Xue KS, Stevens-Ayers T, Angela P, Campbell JA, et al. Parallel evolution of influenza across multiple spatiotemporal scales. eLife. 2017;6:e26875.

25. Good BH, McDonald MJ, Barrick JE, Lenski RE, Desai MM. The dynamics of molecular evolution over 60,000 generations. Nature. 2017; p. in press.

26. Moeller AH, Caro-Quintero A, Mjungu D, Georgiev AV, et al. Cospeciation of gut microbiota with hominids. Science. 2016;353:380–382.

27. Tikhonov M, Monasson R. A collective phase in resource competition in a highly diverse ecosystem. Phys Rev Lett. 2017;118:048103.

28. Taillefumier T, Posfai A, Meir Y, Wingreen NS. Microbial consortia at steady supply. eLife. 2017;6:e22644.

29. Koskella B, Hall LJ, Metcalf CJE. The microbiome beyond the horizon of ecological and evolutionary theory. Nat Ecol Evol. 2017;1(11):1606–1615. doi:10.1038/s41559-017-0340-2.

30. Tikhonov M, Leach RW, Wingreen NS. Interpreting 16S metagenomic data without clustering to achieve sub-OTU resolution. The ISME journal. 2015;9(1):68.

31. Schloissnig S, Arumugam M, Sunagawa S, Mitreva M, Tap J, Zhu A, et al. Genomic variation landscape of the human gut microbiome. Nature. 2013;493:45–50.

32. Segata N, Waldron L, Ballarini A, Narasimhan V, Jousson O, Huttenhower C. Metagenomic microbial community profiling using unique clade-specific marker genes. Nature methods. 2012;9(8):811.

33. Nayfach S, Rodriguez-Mueller B, Garud N, Pollard KS. An integrated metagenomics pipeline for strain profiling reveals novel patterns of bacterial transmission and biogeography. Genome Res. 2016;26:1612–1625.

34. Scholz M, Ward DV, Pasolli E, Tolio T, Zolfo M, Asnicar F, et al. Strain-level microbial epidemiology and population genomics from shotgun metagenomics. Nature methods. 2016;13(5):435.

35. Truong DT, Tett A, Pasolli E, Huttenhower C, Segata N. Microbial strain-level population structure and genetic diversity from metagenomes. Genome Res. 2017;27:626–638.

36. Costea PI, Coelho LP, Sunagawa S, Munch R, Huerta-Cepas J, Forslund K, et al. Subspecies in the global human gut microbiome. Mol Syst Biol. 2017;13(12):960. doi:10.15252/msb.20177589.

37. Nayfach S, Pollard KS. Population genetic analyses of metagenomes reveal extensive strain-level variation in prevalent humanassociated bacteria. bioRxiv. 2015;.

38. Zolfo M, Tett A, Jousson O, Donati C, Segata N. MetaMLST: multi-locus strain-level bacterial typing from metagenomic samples. Nucleic Acids Res. 2017;45(2):e7.

39. Luo C, Knight R, Siljander H, Knip M, et al. ConStrains identifies microbial strains in metagenomic datasets. Nat Biotechnol. 2015;33:1045–52.

40. Smillie CS, Sauk J, Gevers D, Friedman J, Sung J, Youngster I, et al. Strain Tracking Reveals the Determinants of Bacterial Engraftment in the Human Gut Following Fecal Microbiota Transplantation. Cell Host & Microbe. 2018;23(2):229–240.

41. Kuleshov V, Jiang C, Zhou W, Jahanbani F, Batzoglou S, Snyder M. Synthetic long-read sequencing reveals intraspecies diversity in the human microbiome. Nature Biotechnology. 2016;34(1):64–69.

42. Consortium HMP. A framework for human microbiome research. Nature. 2012;486:215–221.

43. Qin J, Li Y, Cai Z, Li S, et al. A metagenome-wide association study of gut microbiota in type 2 diabetes. Nature. 2012;490:55–60.

44. Lloyd-Price J, Mahurkar A, Rahnavard G, Crabtree J, et al. Strains, functions and dynamics in the expanded Human Microbiome Project. Nature. 2017;in press.

45. Xie H, Guo R, Zhong H, Feng Q, Lan Z, Qin B, et al. Shotgun Metagenomics of 250 Adult Twins Reveals Genetic and Environmental Impacts on the Gut Microbiome. Cell Syst. 2016;3(6):572–584 e3. doi:10.1016/j.cels.2016.10.004.

46. Korpela K, Costea P, Coelho LP, Kandels-Lewis S, Willemsen G, Boomsma DI, et al. Selective maternal seeding and environment shape the human gut microbiome. Genome Res. 2018;28(4):561–568. doi:10.1101/gr.233940.117.

47. Sung W, Ackerman MS, Miller SF, Doak TG, Lynch M. Drift-barrier hypothesis and mutation-rate evolution. Proc Natl Acad Sci USA. 2012;109:18488–18492.

48. Poulsen LK, Licht TR, Rang C, Krogfelt KA, Molin S. Physiological state of *Escherichia coli* BJ4 growing in the large intestines of streptomycin-treated mice. J Bacteriol. 1995;177:5840–5845.

49. Russell SL, Cavanaugh CM. Intrahost Genetic Diversity of Bacterial Symbionts Exhibits Evidence of Mixed Infections and Recombinant Haplotypes. Molecular Biology and Evolution. 2017;msx188.

50. Olm MR, Brown CT, Brooks B, Firek B, et al. Identical bacterial populations colonize premature infant gut, skin, and oral microbiomes and exhibit different *in situ* growth rates. Genome Research. 2017;27:601–612.

51. Faith JJ, Guruge JL, Charbonneau M, Subramanian S, Seedorf H, Goodman AL, et al. The long-term stability of the human gut microbiota. Science. 2013;341(6141):1237439.

52. Wakeley J. Coalescent Theory, an Introduction. Greenwood Village, CO: Roberts and Company; 2009.

53. Cohan FM. What are bacterial species? Annual Reviews in Microbiology. 2002;56(1):457–487.

54. Cohan FM. Transmission in the Origins of Bacterial Diversity, From Ecotypes to Phyla. Microbiology spectrum. 2017;5(5).

55. Voight BF, Pritchard JK. Confounding from cryptic relatedness in case-control association studies. PLoS genetics. 2005;1(3):e32.

56. Asnicar F, Manara S, Zolfo M, Truong DT, Scholz M, Armanini F, et al. Studying vertical microbiome transmission from mothers to infants by strain-level metagenomic profiling. MSystems. 2017;2(1):e00164–16.

57. Roach DJ, Burton JN, Lee C, Stackhouse B, Butler-Wu SM, Cookson BT, et al. A year of infection in the intensive care unit: prospective whole genome sequencing of bacterial clinical isolates reveals cryptic transmissions and novel microbiota. PLoS genetics. 2015;11(7):e1005413.

58. Browne HP, Forster SC, Anonye BO, Kumar N, Neville BA, Stares MD, et al. Culturing of ‘unculturable’ human microbiota reveals novel taxa and extensive sporulation. Nature. 2016;533(7604):543–546. doi:10.1038/nature17645.

59. Smith JM, Smith NH, O’Rourke M, Spratt BG. How clonal are bacteria? Proc Natl Acad Sci USA. 1993;90(10):4384–4388.

60. Ohta T, Kimura M. Linkage disequilibrium at steady state determined by random genetic drift and recurrent mutation. Genetics. 1969;63:229–238.

61. Slatkin M. Linkage disequilibrium – understanding the evolutionary past and mapping the medical future. Nature Reviews Genetics. 2008;9:477–485.

62. Palys T, Nakamura L, Cohan FM. Discovery and classification of ecological diversityin the bacterial world: the role of DNA sequence data. International Journal of Systematic and Evolutionary Microbiology. 1997;47(4):1145–1156.

63. Zhu A, Sunagawa S, Mende DR, Bork P. Inter-individual differences in the gene content of human gut bacterial species. Genome Biol. 2015;16:82.

64. Voigt AY, Costea PI, Kultima JR, Li SS, Zeller G, Sunagawa S, et al. Temporal and technical variability of human gut metagenomes. Genome Biol. 2015;16:73. doi:10.1186/s13059-015-0639-8.

65. Greenblum S, Carr R, Borenstein E. Extensive strain-level copy-number variation across human gut microbiome species. Cell. 2015;160:583–94.

66. Donaldson GP, Lee SM, Mazmanian SK. Gut biogeography of the bacterial microbiota. Nature Reviews Microbiology. 2016;14(1):20.

67. Tenaillon O, Rodríguez-Verdugo A, Gaut RL, McDonald P, Bennett AF, Long AD, et al. The Molecular Diversity of Adaptive Convergence. Science. 2012;335:457–461.

68. Good BH, Desai MM. Deleterious passengers in adapting populations. Genetics. 2014;198:1183–1208.

69. Smillie CS, Smith MB, Friedman J, Cordero OX, David LA, Alm EJ. Ecology drives a global network of gene exchange connecting the human microbiome. Nature. 2011;480(7376):241–4. doi:10.1038/nature10571.

70. Brito IL, Yilmaz S, Huang K, Xu L, Jupiter SD, Jenkins AP, et al. Mobile genes in the human microbiome are structured from global to individual scales. Nature. 2016;535(7612):435–439. doi:10.1038/nature18927.

71. Vos M, Didelot X. A comparison of homologous recombination rates in bacteria and archaea. ISME J. 2009;3(2):199–208. doi:10.1038/ismej.2008.93.

72. Guttman DS, Dykhuizen DE. Clonal divergence in *Escherichia coli* asa result of recombination, not mutation. Science. 1994;266:1380–1383.

73. Suerbaum S, Smith JM, Bapumia K, Morelli G, Smith NH, Kunstmann E, et al. Free recombination within Helicobacter pylori. Proc Natl Acad Sci U S A. 1998;95(21):12619–24.

74. Didelot X, Meric G, Falush D, Darling AE. Impact of homologous and non-homologous recombination in the genomic evolution of Escherichia coli. BMC Genomics. 2012;13:256. doi:10.1186/1471-2164-13-256.

75. Dixit PD, Pang TY, Studier FW, Maslov S. Recombinant transfer in the basic genome of Escherichia coli. Proc Natl Acad Sci U S A. 2015;112(29):9070–5. doi:10.1073/pnas.1510839112.

76. Neher RA, Shraiman BI. Statistical genetics and evolution of quantitative traits. Rev Mod Phys. 2011;83:1283–1300.

77. Huynen MA, Bork P. Measuring genome evolution. Proceedings of the National Academy of Sciences. 1998;95(11):5849–5856.

78. Dixit PD, Pang TY, Maslov S. Recombination-driven genome evolution and stability ofbacterial species. Genetics. 2017;207:281–295.

79. Koeppel A, Perry EB, Sikorski J, Krizanc D, Warner A, Ward DM, et al. Identifyingthe fundamental units of bacterial diversity: a paradigm shift to incorporate ecology into bacterial systematics. Proceedings of the National Academy of Sciences. 2008;105(7):2504–2509.

80. Ochman H, Lawrence JG, Groisman EA. Lateral gene transfer and the nature of bacterial innovation. Nature. 2000;405:299–304.

81. Whitlock MC. Fixation probability and time in subdivided populations. Genetics. 2003;164:967–779.

82. Majewski J, Cohan FM. Adapt globally, act locally: the effect of selective sweeps on bacterial sequence diversity. Genetics. 1999;152(4):1459–1474.

83. Shapiro BJ, Polz MF. Ordering microbial diversity into ecologically and geneticallycohesive units. Trends in microbiology. 2014;22(5):235–247.

84. Cohan FM. Bacterial speciation: genetic sweeps in bacterial species. Current Biology. 2016;26(3):R112–R115.

85. Karasov T, Messer PW, Petrov DA. Evidence that adaptation in *Drosophila* is not limited by mutation at single sites. PLoS Genetics. 2010;6:e1000924.

86. Ghalayini M, Launay A, Bridier-Nahmias A, Clermont O, Denamur E, Lescat M, et al. Evolution of a dominant natural isolate of *Escherichia coli* in the human gut over a year suggests a neutral evolution with reduced effective population size. Applied and environmental microbiology. 2018; p. AEM–02377.

87. Zhao S, Lieberman TD, Poyet M, Groussin M, Gibbons SM, Xavier RJ, et al. Adaptive evolution within the gut microbiome of individual people. bioRxiv. 2017; p. 208009.

88. Press MO, Wiser AH, Kronenberg ZN, Langford KW, Shakya M, Lo CC, et al. Hi-C deconvolution of a human gut microbiome yields high-quality draft genomes and reveals plasmid-genome interactions. bioRxiv. 2017; p. 198713.

89. Moss E, Bishara A, Tkachenko E, Kang JB, Andermann TM, Wood C, et al. De novo assembly of microbial genomes from human gut metagenomes using barcoded short read sequences. bioRxiv. 2017; p. 125211.

90. Abubucker S, Segata N, Goll J, Schubert AM, Izard J, Cantarel BL, et al. Metabolic reconstruction for metagenomic data and its application to the human microbiome. PLoS Comput Biol. 2012;8(6):e1002358. doi:10.1371/journal.pcbi.1002358.

91. Scholz M, Ward DV, Pasolli E, Tolio T, Zolfo M, Asnicar F, et al. Strain-level microbial epidemiology and population genomics from shotgun metagenomics. Nat Methods. 2016;13(5):435–8. doi:10.1038/nmeth.3802.

92. Nielsen HB, Almeida M, Juncker AS, Rasmussen S, Li J, Sunagawa S, et al. Identification and assembly of genomes and genetic elements in complex metagenomic samples without using reference genomes. Nature biotechnology. 2014;32(8):822.

93. Quince C, Delmont TO, Raguideau S, Alneberg J, Darling AE, Collins G, et al. DESMAN: a new tool for de novo extraction of strains from metagenomes. Genome biology. 2017;18(1):181.

94. Langmead B, Salzberg S. Fast gapped-read alignment with Bowtie 2. Nature Methods. 2012;9:357–359.

95. Wu D, Jospin G, Eisen JA. Systematic identification of gene families for use as “markers” for phylogenetic and phylogeny-driven ecological studies of bacteriaand archaea and their major subgroups. PLoS One. 2013;8(10):e77033. doi:10.1371/journal.pone.0077033.

96. Korem T, Zeevi D, Suez J, Weinberger A, et al. Growth dynamics of gut microbiota in health and disease inferred from single metagenomic samples. Science. 2015;349(6252):1101–1106.

97. Wexler AG, Goodman AL. An insider’s perspective: Bacteroides as a window into the microbiome. Nature microbiology. 2017;2(5):17026.

98. Coyne MJ, Zitomersky NL, McGuire AM, Earl AM, Comstock LE. Evidence of extensive DNA transfer between bacteroidales species within the human gut. MBio. 2014;5(3):e01305–14. doi:10.1128/mBio.01305-14.

99. Wattam AR, Davis JJ, Assaf R, Boisvert S, Brettin T, Bun C, et al. Improvements to PATRIC, the all-bacterial Bioinformatics Database and Analysis Resource Center. Nucleic Acids Res. 2017;45(D1):D535–D542. doi:10.1093/nar/gkw1017.

100. Li H, Handsaker B, Wysoker A, Fennell T, et al. The Sequence Alignment/Map format and SAMtools. Bioinformatics. 2009;25:2078–2079.

101. Edgar RC. Search and clustering orders of magnitude faster than BLAST. Bioinformatics. 2010;26(19):2460–2461.

102. Birky CW Jr, Walsh JB. Effects of linkage on rates of molecular evolution. Proc Natl Acad Sci USA. 1988;85:6414–6418.

103. Deatherage DE, Barrick JE. Identification of mutations in laboratory-evolved microbes from next-generation sequencing data using breseq. Methods Mol Biol. 2014;1151:165–188.

104. Angly FE, Willner D, Rohwer F, Hugenholtz P, Tyson GW. Grinder: a versatile amplicon and shotgun sequence simulator. Nucleic Acids Res. 2012;40(12):e94. doi:10.1093/nar/gks251.

105. Rödelsperger C, Neher RA, Weller AM, Eberhardt G, Witte H, Mayer WE, et al. Characterization of Genetic Diversity in the Nematode *Pristionchus Pacificus* from Population-Scale Resequencing Data. Genetics. 2014;196:1153–1165.

106. Rocha EP, Smith JM, Hurst LD, Holden MT, et al. Comparisons of dN/dS are time dependent for closely related bacterial genomes. J Theor Biol. 2006;239:226–235.

107. Everitt RG, Didelot X, Batty EM, Miller RR, Knox K, Young BC, et al. Mobile elements drive recombination hotspots in the core genome of Staphylococcus aureus. Nature communications. 2014;5:3956.

108. Bobay LM, Ochamn H. Biological species are universal across Life’s Domains. Genome Biol Evol. 2017;9:491–501.

109. Jones E, Oliphant T, Peterson P, et al.. SciPy: Open source scientific tools for Python; 2001–. Available from: http://www.scipy.org/.

110. McVean GAT. A Genealogical Interpretation of Linkage Disequilibrium. Genetics. 2002;162:987–991.

111. Sprent P, Smeeton NC. Applied nonparametric statistical methods. Chapman and Hall/CRC; 2000.

112. Landau L, Lifshitz E. Statistical Physics, Part 1: Volume 5. Butterworth-Heinemann; 1980.

